# Dimensionality and modularity of adaptive variation: Divergence in threespine stickleback from diverse environments

**DOI:** 10.1101/2021.10.31.466679

**Authors:** Grant E. Haines, Louis Moisan, Alison M. Derry, Andrew P. Hendry

## Abstract

In nature, populations are subjected to a wide variety of environmental conditions that affect fitness and induce adaptive or plastic responses in traits, resulting in phenotypic divergence between populations. The dimensionality of that divergence, however, remains contentious. At the extremes, some contend that populations diverge along a single axis of trait covariance with greatest availability of heritable variation, even if this does not lead a population directly to its fitness optimum. Those at the other extreme argue that selection can push populations towards their fitness optima along multiple phenotype axes simultaneously, resulting in divergence in numerous dimensions. Here, we address this debate using populations of threespine stickleback (*Gasterosteus aculeatus*) in the Cook Inlet region of southern Alaska from lakes with contrasting ecological conditions. We calculated effective dimensionality of divergence in several trait suites (defensive, swimming, and trophic) thought to be under correlated selection pressures, as well as across all traits. We also tested for integration among the trait suites and between each trait suite and the environment. We found that populations in the Cook Inlet radiation exhibit dimensionality of phenotype high enough to preclude a single axis of divergence.

## Introduction

Evolutionary biologists have long grappled with uncertainty regarding the dimensionality of evolutionary processes and patterns, which has important implications for understanding rates and patterns of evolution, as well as their constraints. Although the dimensionality of evolution can be considered from selection oriented or genetic variance oriented perspectives, it can be broadly simplified to two extremes. At one extreme, divergence is constrained by the major axis of the genetic covariance matrix (the “**G** matrix”), to a “genetic line of least resistance” (**G_max_**), irrespective of the strength, complexity, or direction of selection. At the other extreme, high-dimensional divergence is expected to be facilitated both by a complex selection landscape influencing mostly uncorrelated traits with genetic variance distributed across them (Mezey and Houle 2005; Nosil et al. 2009). (Schluter 1996; Hine et al. 2014).

The dimensionality of evolution has important implications for rates of evolutionary change, the directions in trait space in which adaptation is possible, and the number of niche spaces that can be filled. For instance, if the dimensionality of traits is low and correlated traits experience opposing selection pressures, the rate of evolution will be lower than if selection was focused in a single direction similar to the main axis of genetic variation (Agrawal and Stinchcombe 2009). By contrast, high dimensionality of traits could mean fewer constraints on the number of possible combinations of trait values, and thus a greater number of ecological niches that can be rapidly filled by adaptation (Kirkpatrick 2009). Further, multidimensionality of diversification also promotes many-to-one mapping of phenotype to function; that is, the existence of multiple possible phenotypic “solutions” to selection that results in similar fitness (Alfaro et al. 2005; Wainwright et al. 2005), and accommodates the ability of populations to adapt to different peaks on a rugged fitness landscape (Langerhans 2018). In this sense, high-dimensional phenotypic divergence is more aligned with the empirical evidence on the limits of parallel evolution to similar selection pressures than unidimensional divergence (Oke et al. 2017; Langerhans 2018).

One way of conceptualizing and quantifying evolutionary dimensionality is through the related concepts of modularity and integration. Modules are defined as groups of traits that are strongly correlated with each other, but less strongly correlated with other groups of traits (Wagner and Altenberg 1996; Mitteroecker and Bookstein 2007; Wagner et al. 2007). Integration describes correlations between traits or between trait groups that do not permit independent variation (Cheverud 1982; Wagner and Altenberg 1996; Mitteroecker and Bookstein 2007; Adams and Collyer 2016). Integration between modules can constrain evolution by limiting the number of possible evolutionary trajectories without reorganization of the **G**-matrix (Arnold 1992). A high degree of modularity allows for more flexibility in evolutionary trajectories because of the relative independence of each trait group (Wagner and Altenberg 1996), although it has been predicted that dimensionally complex evolution may proceed at a slower rate than evolution along a single axis (Welch and Waxman 2003). However, measuring dimensionality of evolutionary change is made more difficult by patterns of modularity, which results in its uneven distribution across traits.

Most previous studies have tended to focus on associations within one or two trait modules of functionally similar traits likely experiencing similar selection pressures, eliminating the possibility of exploring how integration and modularity expand or contract the potential adaptive range of the phenotypes. For example, Schluter (1996) focused on traits related to trophic ecology in sticklebacks, Darwin’s finches, and sparrows. Hine et al. (2014) considered only cuticular hydrocarbons in *Drosophila.* Hansen & Houle (2008) used only different measurements of *Drosophila* wing shape. Kirkpatrick (2009) used – in separate analyses – fat and muscle traits for beef cattle, body shape measurements in the fish *M. eachamensis,* wing shape measurements in *Drosophila*, lactation curves for dairy cows, and growth trajectories for mice. We suggest that these previous assessments of associations among functionally similar traits are perhaps more likely to represent one or only a few trait modules, which would result in an underestimate of evolutionary dimensionality and an overestimate of the constraints on evolution. Hence, our goal in the present study is to assess the extent of the dimensionality of evolutionary divergence within and between hypothesized modules (hereafter “trait suites”) with different selection pressures, and the degree to which the measurement of dimensionality is influenced by trait modularity and integration. We did this by considering multiple traits in each of three trait suites of different functional categories in evolutionarily divergent populations, making it unlikely that apparent restriction of phenotypic divergence primarily to a single dimension is a consequence of the traits being under similar selection pressures.

### Study system and questions

Threespine stickleback (*Gasterosteus aculeatus*) are an excellent species for studying evolutionary dimensionality through the lens of modularity and integration because they are highly variable within and among populations, and because multiple freshwater populations are independently derived from ancestral marine populations (Bell and Foster 1994). Many of their variable traits are known to be correlated with environmental variables (Bell et al. 1993; Reimchen 1994; MacColl et al. 2013; McGee et al. 2013) that can be used to classify traits into functional suites based on the source(s) of selection they experience. Furthermore, stickleback divergence between habitat types (e.g., lake-stream (Lavin and McPhail 1993), benthic-limnetic (McPhail 1993), and marine-freshwater (Gelmond et al. 2009)) has been described as occurring along axes corresponding to habitat, but whether a single axis can accurately describe any of these divergence patterns is uncertain.

Here, we ask how much variation in phenotype can be explained by a single axis of divergence, and whether phenotypic variation among variable habitats is better explained by a multifarious or unidimensional model of trait divergence. If trait divergence is close to unidimensional, then it may be constrained either by similarly low-dimensional selection, or by strong genetic or developmental integration between traits. Our study focuses on lacustrine environments, so the benthic-limnetic axis of divergence is likely to represent a major axis of phenotypic differentiation, and will occasionally be referred to give an impression of the differences being described. This axis can be defined by a stouter and deeper body shape, a shortened head with a downturned jaw, and fewer gill rakers at the benthic end, and more elongate, zooplanktivory-adapted fish at the limnetic end (Bentzen et al. 1984; McPhail 1993; Willacker et al. 2010). If divergence is multidimensional, it will occur along multiple axes that are selected upon by different environmental variables.

We collected stickleback and conducted environmental surveys at 14 lakes in the Cook Inlet region of southern Alaska. We divided trait measurements of the collected stickleback – *a priori* – into four functional suites (defensive, trophic, swimming, body shape) that could be under differing selection pressures or respond plastically to different environmental conditions. We then used these morphological data to answer four main questions (Figure 1).

**Figure 1.**
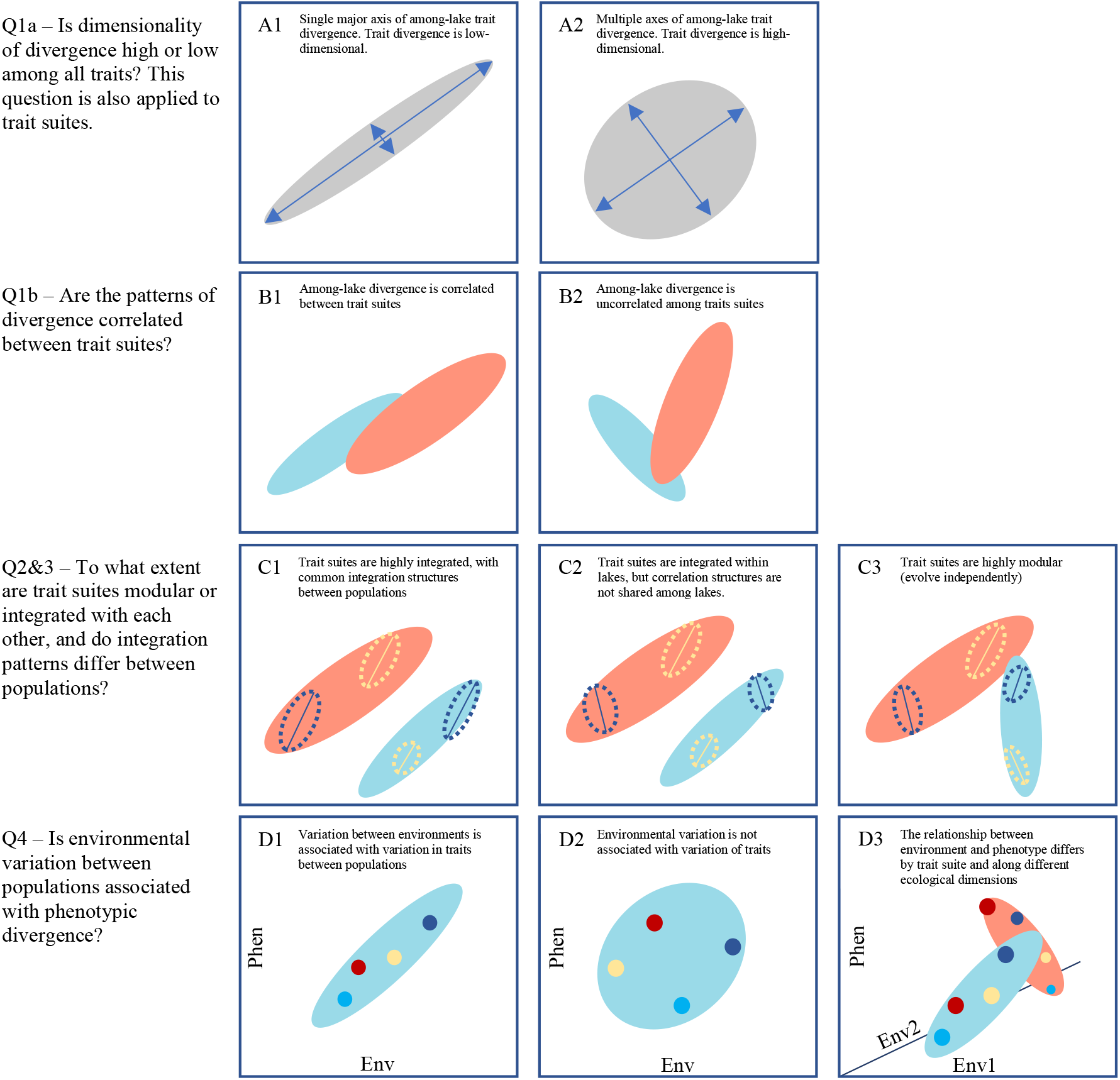
Conceptual diagrams illustrating the potential results of the questions addressed in this paper. Grey ellipses represent total variation in phenotype between populations. Filled, colored ellipses represent variation within trait suites between populations, open dashed ellipses represent populations, and solid circles represent population means. Lines bisecting population ellipses indicate the major axis of variation within that population for the trait suite in which it is shown. Panel axes, except for in D1 -D3, are arbitrary dimensions in a shared trait space, but trait suite ellipses are separated in some panels for illustrative purposes. For the tests that correspond with the panels, please see Table A1.

1) To what extent do lacustrine stickleback diversify along a single axis of divergence among populations, and what is the effective dimensionality of that divergence? Although there is no formal null expectation of effective dimensionality for this question, we predicted that populations of stickleback confined to lakes diversified along a single axis of correlated variation between phenotypes, and that divergence is largely a unidimensional process. We applied this question to each of the pre-defined functional trait suites, as well as a single dataset including all traits. If there is a single primary axis of divergence, the primary axes of the trait suites will be correlated. If trait suites are uncorrelated, then dimensionality of divergence is not constrained to the major axis of the **G** matrix unless the trait suites themselves are highly multidimensional.

2) To what extent do lakes share common within-lake correlation structures between trait modules? We predict that if trait suites are closely correlated, then lakes will share common within-lake correlation structures between trait suites, reflecting conserved covariance among traits suites.

3) To what extent do functional trait suites covary as adaptive modules? Trait suites that constitute modules would strongly covary internally, and be representative of axes of divergence that could be predicted based on the environmental conditions imposing selection on the module, irrespective of pressures on other traits or trait suites.

4) To what extent are different environmental variables, or groups of environmental variables, associated with variation in all phenoptypes and in the trait suites? Phenotypic divergence corresponding with environmental variation would indicate that the phenotypic variation is likely adaptive or shares common plastic responses, and that it is only weakly constrained by correlations between trait suites.

From previous work, we do have some *a priori* expectations of trait responses to the environment. For defensive traits, low dissolved calcium concentration in the absence of predators is strongly predictive of reduction of the pelvis and spines (Giles 1983; Bell et al. 1993). Although some of the lakes we sampled have stocked or native salmonids that could act as predators, only limited information about the predator communities of most of the lakes is available (ADF&G 2015). Robustness of defensive traits is also associated with larger, clearwater lakes with higher visibility for predators (Reimchen et al. 2013), and these lakes tend to have fish with narrower caudal peduncles and body shapes with more anteriorly-shifted dorsal spines and dorsal and anal fins – traits that are not directly related to defense (Spoljaric and Reimchen 2007). Willacker et al. (2010) also found that depth-related variables were significantly correlated with the benthic-limnetic axis for skull morphology. One population – that of G Lake – was known to have an introduced population of Muskellunge (*Esox masquinongy*; Lipka et al. 2020; Massengill et al. 2020), a congener of Northern Pike (*E. lucius*), which Willacker et al. (2010) found was associated with more limnetic cranial morphologies in stickleback than those in populations from otherwise comparable habitats. We expect that lakes with environmental variables that depart from the mean lake in more dimensions will have stickleback that are divergent in more directions. We expect that trophic traits could depend on nutrient availability, mediated by the prey community. However, because of the dynamic feedbacks between prey availability and trophic morphology, the direction of causality between morphology and some environmental variables that influence predator-prey eco-evo relationships is not clear (Brooks and Dodson 1965; Palkovacs and Post 2008, 2009; Schmid et al. 2019).

## Methods

### Study system

In May and June of 2018, we surveyed 14 lacustrine populations of threespine stickleback from the Cook Inlet region of southwest Alaska. The lakes surveyed were: Corcoran, Finger, Long, Ruth, South Rolly, and Walby Lakes in the Matanuska-Susitna Valley, and Echo, Engineer, G, Jean, Spirit, Tern, Wik, and Watson Lakes on the Kenai Peninsula (Figure 2). These populations were targeted in an effort to maximize morphological diversity along the benthic-limnetic axis of divergence, while including some intermediate populations, using a combination of Willacker et al. (2010) and Alaska Department of Fish & Game (personal communication). This design increased the likelihood that populations had sufficient phenotypic diversity to detect multidimensionality of divergence if it is present in wild populations more generally, and although some of these populations are close to one another geographically, they do not share surface connections that are plausible routes of dispersion. Generally, in the Cook Inlet region of Alaska (Willacker et al. 2010) – as elsewhere in the species range (Schluter and McPhail 1992), “benthic” stickleback are adapted to feeding on benthic macroinvertebrates and have shorter heads and deeper bodies, whereas “limnetic” stickleback are adapted to feeding on zooplankton, and have more elongate body and head shapes.

**Figure 2.**
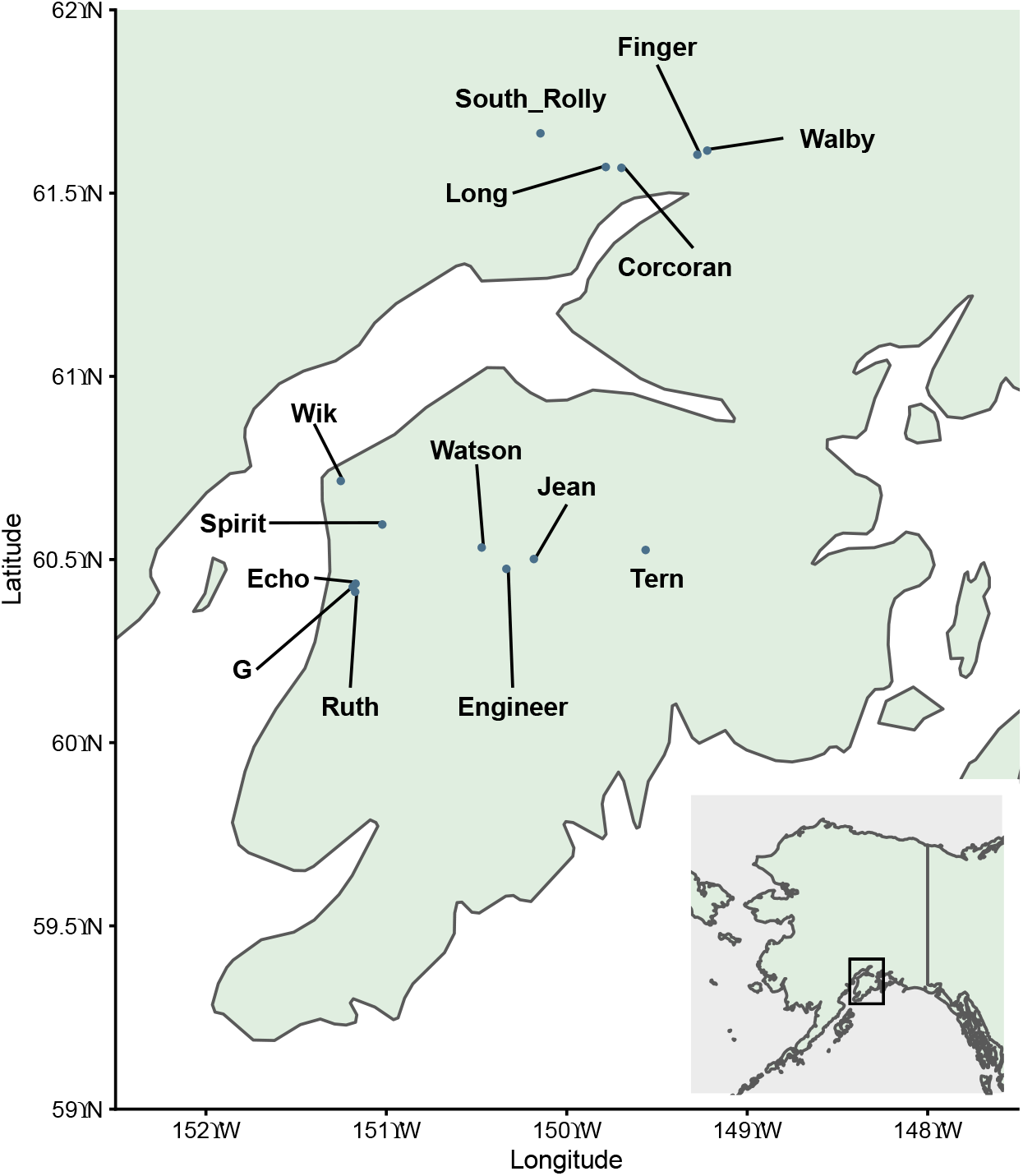
Locations of lakes in the Cook Inlet region of southern Alaska that were sampled for this study.

### Linear measurements and meristic traits

Stickleback were captured in unbaited minnow traps and then killed with an overdose of clove oil, following the animal use protocol obtained from McGill University. Each fish was photographed immediately after death on 1mm grid paper with a Nikon D800 (Finger, Walby, and 31 Long Lake fish) or a Canon PowerShot G11 (all others), and then preserved in 95% EtOH. Upon their return to the lab, the following measurements were taken on fish by means of digital callipers with resolution to a hundredth of a millimeter: standard length (SL), body depth (BD), gape width (GW), buccal cavity length (BC) and the length of the pelvic and first and second dorsal spines (PS_L_, DS1_L_, and DS2_L_, respectively; Figure 3). We also measured maximum width of the posterior process of the pterotic (PW), which articulates with the latero-anterior portion of the post-temporal as an alternative to epaxial width. Following staining with alizarin red-S dye as described by Springer & Johnson (2000), we counted lateral gill rakers of the first right branchial arch (GR), and the number of lateral plates (LP) on the both sides of the body (the average of the two sides was used for analysis). Using the digital photos, the following measurements were made in ImageJ (Rasband 1997-2018): caudal peduncle width (CP), upper jaw length (JL), snout length (SN), head length (HL), and eye diameter (ED) (Figure 3).

**Figure 3.**
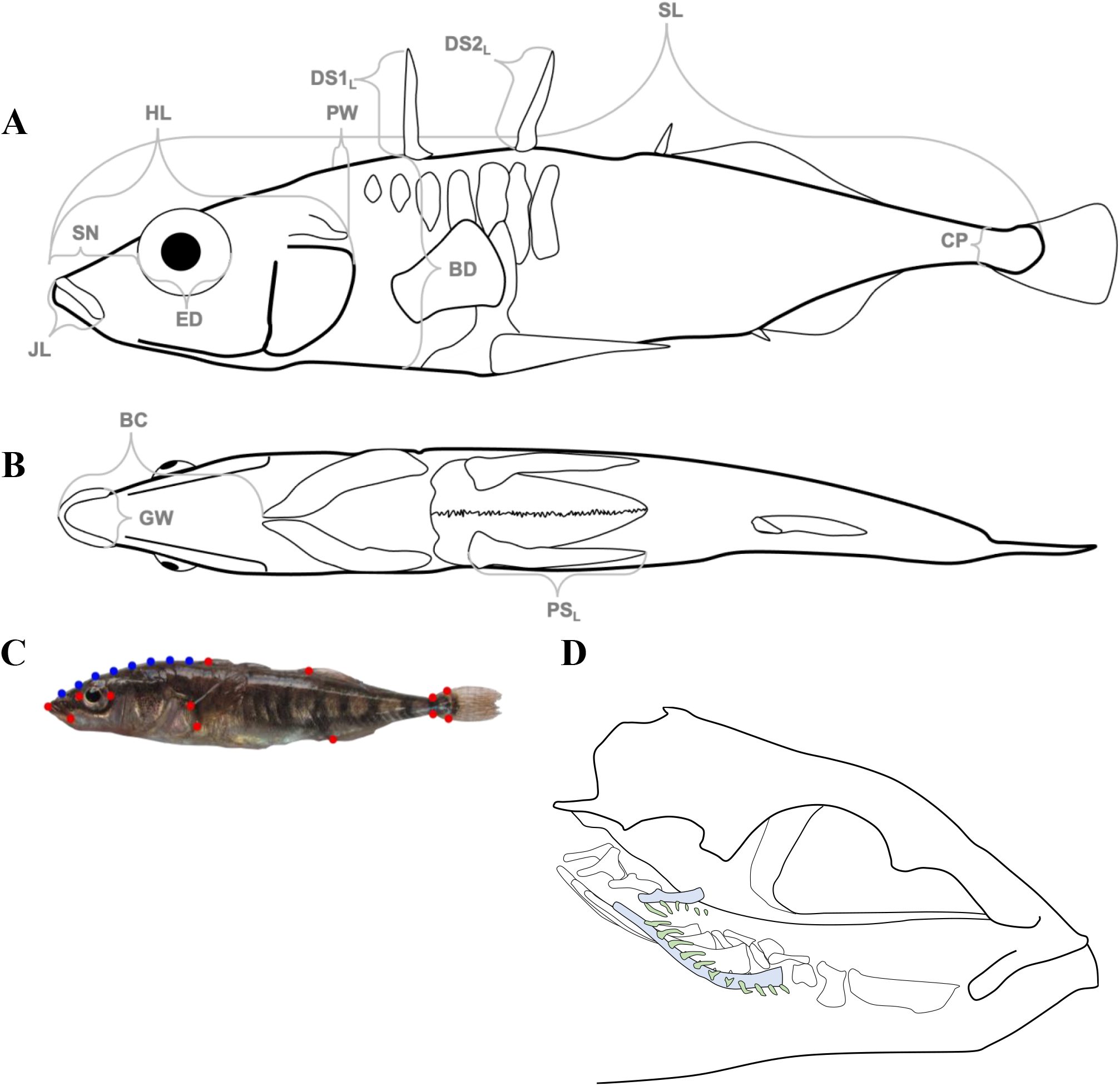
Univariate morphometric measurements from lateral (A) and ventral (B) views, geometric morphometric landmarks (red) and semilandmarks (blue; C) and the 1^st^ branchial arch gill rakers (GR) that were counted (D, in green). Measurement shown are: JL – upper jaw length; SL – snout length; HL – head length; PW – pterotic width (width measured at widest point of pterotic posterior process); DS1_L_ and DS2_L_ – dorsal spine lengths; BD – standard length; CP – caudal peduncle width; BC – buccal cavity length; GW – gape width; PS_L_ – pelvic spine length. Medial rakers on the first right branchial arch (blue) were not counted (C). Only ossified lateral rakers (green) were counted.

Linear measurements were adjusted for allometry using the formula 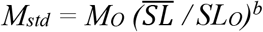, where *Mo* and *SLo* are the observed trait length and standard length, respectively, 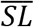 is the grand mean of all standard lengths (46.50 mm), and *b* is the slope of an ANCOVA of the form *M* ~ *SL* + *Lake* (Reist 1986; Hendry and Taylor 2004). GR, LP, and LP_M_ were not adjusted for allometry because all but 2 of our 792 fish had standard lengths < 30 mm, the approximate size by which all gill rakers, lateral plates, and pelvic bones have formed (Bell 1981, 1987, 2001; Glazer et al. 2014). Spearman’s rank correlations, with p-values adjusted for multiple comparisons using the false detection rate (FDR) method (Benjamini and Hochberg 1995), confirmed that GR and LP were not correlated with SL in any lake.

Following Stuart et al. (2017), univariate measurements were then assigned to one of three trait suites (Defense, Trophic, Swimming) based on the type of selection they are expected to experience most directly (Table S1). Although some traits could be associated with more than one trait suite based on selection in the environment, each had to be assigned to a single trait suite for methodological reasons. Additionally, the tests of correlation between trait suites described below implicitly acknowledge the potential for functional overlap between trait suites. Missing values of traits, except GR, which had by far the lowest *N* of all traits recorded (*N_GR_* = 276, or 34% of specimens), were estimated using the regression formula of the most correlated trait with *r* > 0.1 within the same trait suite (numbers of individuals for which traits were thus estimated are provided in Table S2). Units of traits were then standardized for multivariate analyses by transforming units into standard deviations of the total sample, and centered about zero (Z-transformation). All data analyses were performed using the *R* statistical environment, version 3.6 (R Core Team 2019).

### Geometric morphometrics

Body shape was quantified using landmark-based geometric morphometric (GM) methods, with thirteen homologous anatomical landmarks placed from mouth to tail, to which eight semi-landmarks were added to capture the curvature of the head from the tip of the snout to the first dorsal spine (Figure 3). Landmarks and semi-landmarks were placed using tpsDig2 2.31 (Rohlf 2018). A Procrustes superimposition was performed on all specimens using the *gpagen* function in the *R* package *geomorph* (Adams et al. 2019), with curves defined using minimum bending energy of the semi-landmarks. Photographs of Long Lake specimens taken with the two different cameras were found to be significantly different with a Procrustes ANOVA, so landmark data of specimens photographed with the Nikon D800 was excluded from geometric morphometric analysis. Specimens for which the location of the landmark at the base of the first dorsal spine was uncertain were excluded, as this landmark was used to place the series of eight semi-landmarks. Repeatability of shape was assessed to be 92% using variance ratios from Procrustes ANOVA as described by Zelditch et al (2012).

Because the GM data was available for two fewer lakes than the data describing the other trait suites, the subsequent analyses were conducted both including and excluding the GM data, with results provided in the Supplementary Tables when they are not provided in the Results section. Although shape is described as a trait suite throughout this paper, it is expected to be highly integrated with all three of the other trait suites. This is because it is associated with Defense from gape-limited predators (Reimchen 1991), with Defense and Swimming traits via its association with the fast-start response for predator avoidance (Taylor and McPhail 1986; Bergstrom 2002), and with Swimming and Trophic traits via the relationship between body depth and maneuverability in spatially complex foraging environments (Walker 1997; Feilich 2016). Additionally, linear measurements included in other trait suites, especially BD, ED, CP, and HL describe aspects of shape, and thus these morphological traits are inextricably linked with the shape data.

### Limnological Conditions and Dietary Indicators

Lake physico-chemical properties were measured in June of 2018 and June of 2019. These included total nitrogen (TN), total phosphorus (TP), chlorophyll-*a* (Chl-*a*), dissolved calcium (Ca), dissolved organic carbon (DOC), conductivity (Cond.), and pH. These were all measured from epilimnetic water samples at the deepest point in each lake. Total surface area and maximum depth were obtained from previous Alaska Department of Fish & Game (ADF&G) surveys. The proportion of littoral area (defined as area < 3 m deep) was calculated using ADF&G bathymetric maps when such maps were available. Zooplankton and benthic invertebrate community samples were collected using vertical Wisconsin net tows, and D-frame kick net sampling, respectively, and were used to estimate the densities of a large limnetic grazer, *Daphnia* sp., in the pelagic zone and the densities of a dominant benthic macroinvertebrate family, gammarids, in the littoral zone of each lake. *Daphnia* sp. and gammarids are important components of the stickleback diet when they are present. Details of these field and laboratory methods are provided in Appendix A.

### Statistical Analyses

#### Question 1 Dimensionality of divergence, and correlations of trait suites

We examined the extent to which trait divergence among lake populations was multidimensional by first applying a linear discriminant analysis (LDA) including traits from all trait suites (Venables and Ripley 2002; Ripley et al. 2020). The LDA determined the axis of greatest variation among lakes in relation to the variation within them. It is important here to note that unlike PCA axes, LDA axes are not orthogonal because LDA scales groups by their variances. For our rationale behind using LDAs, please see Appendix A: Supplementary Methods. Because Canonical Variate Analysis (CVA) and multivariate LDA are identical operations, this analysis was performed on shape data using the *CVA* function in the *R* package *Morpho* (Schlager 2017; Schlager et al. 2020), which provided an output that was more convenient for shape visualization, but yielded the same results as *lda* in *MASS* (Ripley et al. 2020).

We next calculated the number of effective number of dimensions of multivariate divergence in all traits using several methods reviewed by Del Giudice (2020), shown in Table 1. The third and fourth of these indices are very conservative and consistently overestimate effective dimensionality (*D_e_*), respectively, but we include them for purposes of comparison to their usage elsewhere in the evolutionary biology literature. Because LDA is performed by singular value decomposition, we use the squares of the singular values (Ripley et al. 2020; Watanabe 2021) – which are the canonical F-stats and used to calculate proportions of the trace explained by each LD axis – in place of eigenvalues in the *D_e_* calculations. Despite our above objections to the use of eigenvectors to calculate divergence, we also calculated effective dimensionality of the population-level trait-space using eigenvalues of population means as well the trait-space using eigenvalues of individual-level data for the purposes of comparison to other work and to place the *D_e_* calculations from LDs in context. Dimensionality is a continuous variable, and so cut-offs for categorizing low, moderate, and high values are somewhat arbitrary. However, because these categories ease discussion, we regard “low” effective dimensionality as < 2, “moderate” from 2 – 3, and “high” > 3. These values were chosen because the dimensionality of a circle projected onto a plane is 2, and the dimensionality of a sphere is 3, making their interpretation somewhat more intuitive than other thresholds. For convenience, we will refer mostly to the Dn1 effective dimensionality index unless otherwise noted. Because we are testing the extent to which this radiation aligns with to hypotheses along a continuum of dimensionality, there is no null hypothesis, and expectations of dimensionality are likely to vary by system and environmental complexity.

**Table 1.** Indices of effective dimensionality and variables used in their calculation. Details of these indices and their properties are described further and reviewed in Del Giudice (2020). All indices are bounded by a minimum of 1 and a maximum of *n*.

| $D_e$ Index | Formula | Reference(s) | Notes |
| --- | --- | --- | --- |
| $D_{n1}$ | $\prod_{i=1}^n \left( \frac{\lambda_i}{\sum_{i=1}^n \lambda_i} \right)^{-\frac{\lambda_i}{\sum_{i=1}^n \lambda_i}}$ | (Cangelosi and Goriely 2007; Roy and Vetterli 2007) | |
| $D_{n2}$ | $\frac{(\sum_{i=1}^n \lambda_i)^2}{\sum_{i=1}^n \lambda_i^2}$ | (Pirkel et al. 2012) | |
| $D_{n\infty}$ | $\sum_{i=1}^n \lambda_i / \lambda_1$ | (Kirkpatrick 2009) | excessively conservative |
| $D_{nC}$ | $n - \frac{n^2}{(\sum_{i=1}^n \lambda_i)^2} \text{var}(\lambda)$ | (Cheverud 2001) | overestimates $D_e$ |
| variable | Description |  |  |
| $\lambda$ | squared singular values of LD axes, or eigenvalues for full data and population means | | |
| $\lambda_1$ | squared singular value of LD1, or largest eigenvalue | | |
| $n$ | the number of squared singular values or eigenvalues | | |

Because gill rakers were counted in only approximately one third of specimens, the above discriminant analysis for all traits was conducted with and without gill rakers. Additionally, this discriminant analyses excluded geometric morphometric landmark coordinates, both because landmarks excluded two lakes and because some components of shape are redundant with other variables.

As with the dataset of all traits, we conducted LDAs within Defense, Swimming, and Trophic, and Shape trait suites, and calculated effective number of dimensions for each. We tested for correlations between the trait suites using pairwise Pearson correlations of the primary discriminant axes and mantel tests of within-suite distance matrices of the specimens in the calculated directly from standardized trait values. Each mantel test used 5000 iterations. Because the distribution of defensive traits violated the assumptions of Pearson’s *r*, univariate and multivariate correlations including the defensive trait suite were performed using Spearman’s rank correlation test, in addition to the Pearson correlation test.

The effective dimensionality is limited by the number of traits included in each trait suite (e.g., the defensive trait suite, which includes 5 traits, and cannot have an effective dimensionality greater than 5), and so we implemented a rarefaction-like permutation procedure to visualize changes in effective dimensionality of trait suites as traits are accumulated at random, and compare suites of different numbers of traits. For details on this procedure and results, see Appendices A and B, respectively.

#### Question 2 Comparisons of within-lake correlation structures

We used two-block partial least squares correlation (2B-PLS), to test whether the trait suites were correlated within populations. This technique takes two data matrices from the same individuals, for example our data for swimming and defensive trait suites, and identifies the linear vectors through each dataset that maximize covariation explained between the datasets (Rohlf and Corti 2000). We conducted 2B-PLS tests for all pairwise comparisons of trait suites. Each test yielded both a correlation coefficient (*r_PLS_*), and a *Z*-score (Adams et al. 2019). The *Z*-score serves as an effect size of integration between the two blocks (here, trait suites) for each population, and which is comparable between 2B-PLS tests on populations with different sample sizes (Adams and Collyer 2016). We used these standardized effect sizes to compare integration strength of trait suites between populations with pairwise two-sample *Z*-tests (Adams and Collyer 2016; Adams et al. 2019), and characterized the orientation of those differences by calculating the angle between vectors describing the most variance through data blocks between lakes. 2B-PLS tests and between-population *Z*-tests were performed using the *two.b.pls* and *compare.pls* functions, respectively, in the *R* package *geomorph* (Adams et al. 2019). Reported p-values were adjusted for multiple comparisons using the FDR method (Benjamini and Hochberg 1995).

By way of a simple example, in a radiation of three populations (A, B, and C), we could conduct 2B-PLS tests between defensive traits and swimming traits in each population. The *r_PLS_* statistic would give us the level of correlation between the defensive traits and swimming traits. Because *r_PLS_* is sensitive to sample size, however, we compare the strength of correlations between trait suites using *Z*-scores which describe the strength of the effect relative to the null expectation of no association. If, after conducting pairwise *Z*-tests between populations, the comparisons between populations A and B yield a low effect size, but the A-C and B-C comparisons yield high effect sizes, then the strength of integration between defensive and swimming traits is similar between A and B but C differs from both. In this example radiation, then, the strength of the relationship between defense-associated and swimming-associated traits in population C diverged from that of the other two populations. Calculating vector angles between data blocks of the same trait suite for pairwise comparisons between lakes allowed us to compare orientation of integration structures, which are not taken into account in *Z*-tests for differences in strength of effects between lakes.

Because PLS methods are sensitive to both vector magnitude and orientation, our use of correlation matrices does produce different results in our 2B-PLS tests than use of covariance matrices would, (and indeed, we did use unstandardized GM landmarks in these analyses, to the same effect). However, use of a correlation matrix for instances in which data do not all exist on the same scale, as is the case here, is recommended by Rohlf and Corti (2000), and even required for uses of 2B-PLS in which a block of morphological data is associated with a block of environmental data or other non-morphological data related to niche occupation, as environmental data are rarely on the same scale (Corti et al. 1996; Fadda and Corti 1998; Felice et al. 2019).

#### Question 3 Evolutionary modularity of trait suites

We tested the extent of modularity of trait suites, both within each population and across all populations using the *modularity. test* function of *geomorph,* which tests whether the ratio of covariation between to within designated trait suites is less than if traits had been assigned to suites randomly (Adams 2016; Adams and Collyer 2019). When this test is applied to multiple trait suites simultaneously, the average of pairwise covariance ratios (CR) is used as a test statistic (Adams and Collyer 2019), which provides a better picture of overall modularity, but provides less information about the independence of each trait suite. Consequently, these tests were executed in two ways, involving how partitions between trait suites were set. First, we performed the test after defining a partition separating a target trait suite from all other traits. These tests are referred to in results as “focused” tests. We then performed a test of modularity with all trait suites partitioned from one another, which we refer to in results as a “complete” test.

Because the measurements in the swimming trait suite were important components of shape, modularity tests were performed both with and without the inclusion of shape as a module to be sure that this redundancy did not alter results. All modularity tests were performed with 5000 permutations. The p-values reported for these tests are FDR-adjusted for multiple comparisons.

#### Question 4 Responses to the environment

We quantified environmental differences between lakes using principal components analysis (PCA). This was first performed first using physico-chemical variables – pH, DOC, Ca, TN, TP, Maxium Depth, chl-*a*, and surface area – for which all data were available for all but two lakes (Echo and Ruth). This was followed by a PCA including both physico-chemical variables and stickleback foraging-related characteristics of the lakes, for which only a subset of lakes were surveyed. Both environmental PCAs used scaled environmental variables, and thus were calculated using correlation matrices rather than covariance matrices. Stickleback foraging-related lake characteristics included the areal proportion of littoral habitat (< 3 m depth), *Daphnia* sp. abundance in the pelagic zone (number of individuals per L), and gammarid abundance in the littoral zone (number of individuals per m^2^). We also performed Spearman’s correlations between each environmental variable and stickleback trait suite linear discriminants axes to determine the relationship between different environmental variables and axes of divergence (see Appendix B).

We subsequently executed 2B-PLS tests using normalized population-mean morphological variables as one block, and environmental variables as the other. As with PCAs, we performed 2B-PLS tests twice, using both physico-chemical and full environmental datasets, the first including more lakes (n = 12), but the second included data related to foraging (n = 9). 2B-PLS is a balanced analysis which does not assume a direction of causality. It therefore may be a more accurate characterization of our system, in which eco-evo feedbacks potentially play a role, than another multivariate regression technique like redundancy analysis (RDA), which includes multivariate regression as its first step (Legendre et al. 2011). Additionally, 2B-PLS was more suited to our limited sample size because it uses a resampling procedure to generate the distribution for calculation of significance and effect size, and it is thus not limited by available degrees of freedom. Each 2B-PLS test yielded vectors through both the phenotype and environmental blocks representing axes of greatest correlation between the two. The environmental data used in 2B-PLS tests is identical between tests, irrespective of trait suite phenotype. Because of this, the primary environmental vector angles between tests for different trait suites were calculated to quantify the divergence between environmental gradients along which divergence occurs.

## Results

### 1. Axes of divergence for all traits and within trait suites

In the linear discriminant analysis including all defensive, trophic, and swimming traits, dimensionality was calculated to be high (*D*_*n*1_ = 3.26 and *D*_*n*2_ = 2.01) when gill rakers were included (*N* = 241; Tables S3, 2). In this LDA, the primary axis explained 69.0% of the variance between populations, and the first three axis explained 87.2% of the variance. The traits onto which the first axis were most heavily loaded were standard length and pelvic spine length (Table S4). The most important trait in the LDA was pelvic spine length, with a loading on LD1 four times the magnitude of the next largest coefficient, SL. Results were similar when gill rakers were excluded to increase the sample size. In this discriminant analysis (*N* = 549), dimensionality was only slightly lower (*D*_*n*1_ = 3.09 and *D*_*n*2_ = 1.90). The first LD axis described 71.4% of interpopulation variance, and the first three axes described 88.1%.

An LDA of defensive traits showed that variation between populations was driven mostly by a single axis that accounted for 92.1% of this variation. Pelvic spine length (PS_L_) was the variable most heavily loaded onto this axis, and had a loading nearly 8 times higher than DS2_L_, the trait with the next highest loading. this result seems to be driven by the G Lake population, which was entirely lacking pelvic spines, and Echo Lake, which had only a single fish with pelvic spines. Dorsal spine lengths (DS1_L_ and DS2_L_) drove most of the remaining variance in defensive traits between populations, and dimensionality of defensive trait divergence was low (Tables 2 and S5, Figures 5 & S1).

**Table 2.** Dimensionality of trait suites calculated using inertia of linear discriminants, as well as eigenvalues of the full trait matrix and population means calculated from PCAs.

| $D_e$ type | Defense | | | Trophic | | | Swimming | | | Shape | | | Total non-shape | | |
| --- | --- | --- | --- | --- | --- | --- | --- | --- | --- | --- | --- | --- | --- | --- | --- |
|  | LDA | full matrix | pop. means | LDA | full matrix | pop. means | LDA | full matrix | pop. means | LDA | full matrix | pop. means | LDA | full matrix | pop. means |
| **$D_{n1}$** | **1.43** | **4.04** | **3.37** | **4.57** | **2.6** | **2.32** | **2.22** | **1.73** | **1.68** | **7.16** | **10.32** | **5.28** | **3.26** | **5.31** | **4.45** |
| **$D_{n2}$** | **1.18** | **3.63** | **3.00** | **3.47** | **1.75** | **1.66** | **1.90** | **1.39** | **1.36** | **5.27** | **6.63** | **4.23** | **2.01** | **2.99** | **2.99** |
| $D_{n\infty}$ | 1.09 | 2.59 | 2.34 | 2.25 | 1.35 | 1.31 | 1.48 | 1.19 | 1.18 | 2.84 | 3.57 | 2.93 | 1.45 | 1.81 | 1.87 |
| $D_{nc}$ | 1.75 | 4.62 | 4.34 | 6.7 | 4.43 | 4.17 | 2.42 | 1.85 | 1.80 | 9.91 | 36.66 | 33.07 | 7.54 | 11.64 | 11.65 |

Notes: The final group of values represents all traits excluding shape, because a) two populations were excluded from geometric morphometric analysis and b) our swimming traits are largely redundant with our shape data. **$D_{n1}$** and **$D_{n2}$** are shown in bold typeface to indicate that they are the $D_e$ measures that are recommended for consideration, as $D_{n\infty}$ and $D_{nc}$ systematically underestimate and overestimate dimensionality, respectively.

**Figure 5.**
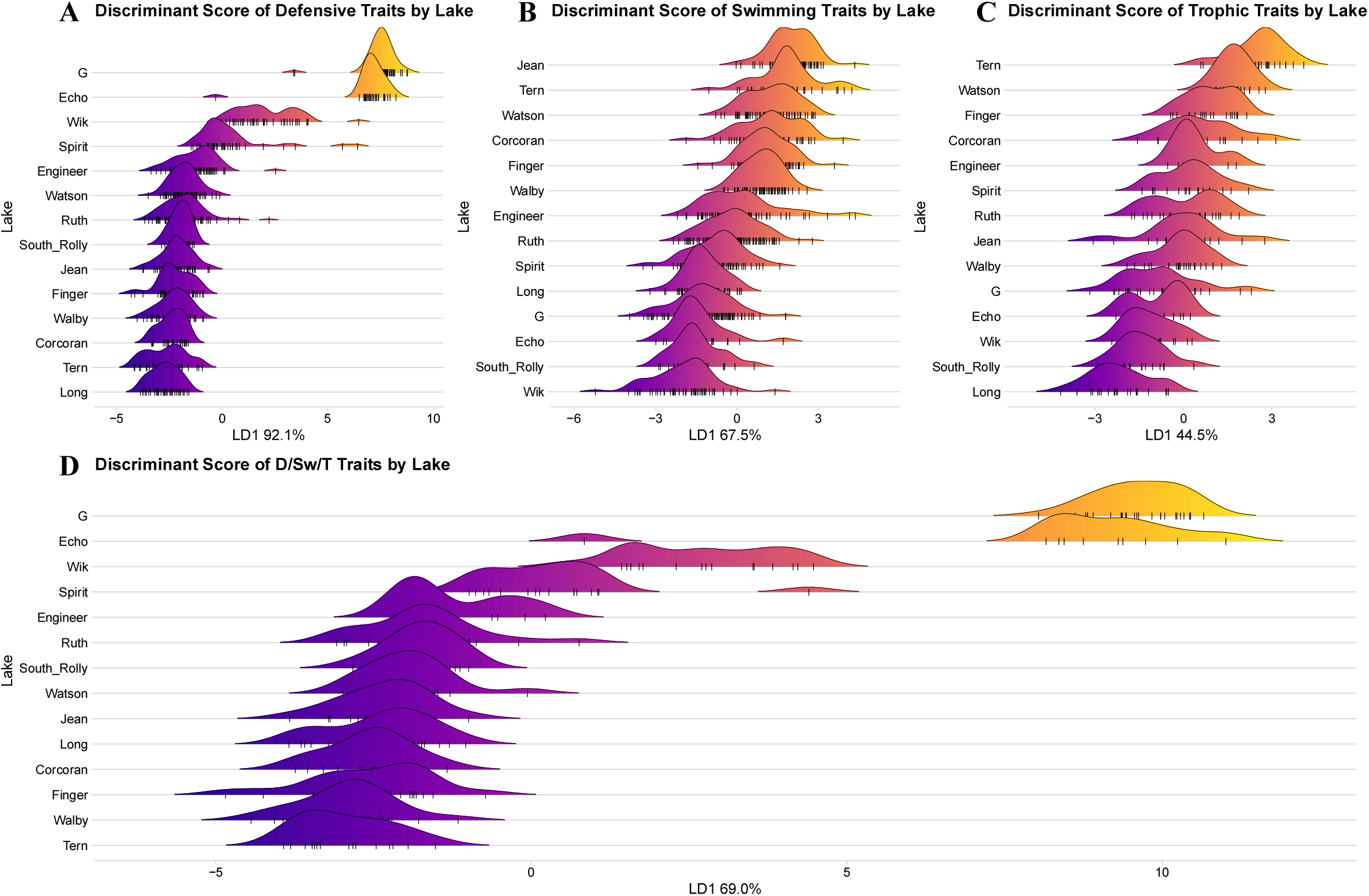
Distribution of individuals along primary linear discriminant axes for Defensive. Swimming, Tropine, and combined traits. Positions of individual fish on each axis are indicated by the tick marks. Primary linear discriminant axes account for 92.1% (Defense), 67.5% (Swiimning), 44.5% (Trophic), and 69.0% (D/Sw/T) of variances in their respective trait suites. Shape is excluded here because divergence is distributed evenly enough across CV axes that visualizing only the first axis would contribute little to comprehension of results, and it is excluded from the combined discriminant score because variables in other trait suites (particularly the swiimning suite) are redundant with important components of shape.

Swimming traits, by contrast, had higher effective dimensionality of divergence than defensive traits, despite the swimming trait LDA including only three variables (SL, BD, and CP; Tables 2 and S6). LD1 of this discriminant function described 67.5% of variation between population, with CP and BD being loaded approximately equally onto this axis. This placement of populations along this axis conformed fairly well to *a priori* expectations of the benthic-limnetic axis, with more apparently limnetic populations, like G, Wik, and South Rolly Lakes, being assigned more negative values – and more benthic populations, like Watson and Tern Lakes, being assigned more positive values.

Trophic trait divergence was much more multidimensional than swimming or defensive trait divergence, with effective dimensionality of *D*_*n*1_ = 4.57 and *D*_*n*2_ = 3.47 (Table 2). This was also much higher than *D_e_* calculated from eigenvalues of individual-level or population mean trait data. In the trophic trait suite, LD1 described only 44.5% of variation between populations with PW, followed closely by HL and SN, being the most heavily loaded variable onto this axis (Table S7).

Body shape was the most multidimensional of the trait suites, with high effective dimensionality of *D*_*n*1_ = 7.16 and *D*_*n*2_ = 5.27, albeit calculated from data with a much higher number of traits, with each landmark having an X and Y coordinate (Table 2). For shape, the first discriminant axis only described 35.2% of variation between populations, and five LD axes were required to explain at least 80%. The first axis here describes mostly body depth, but also angle of the head and the posterior extent of the dorsal fin. As with the swimming and to some extent the trophic suite, LD1 appears to reflect the benthic-limnetic continuum (Figures S1 and S2).

Primary axes of all trait suite LDAs were significantly correlated with all other LD1s (Figure S3; Table S8). The least correlated LD1s were those of the defensive and trophic trait suites (Pearson’s *r* = −0.21, p = 0.001; Spearman’s ρ = −0.10, p = 0.109). The most correlated trait suite LD1s were those of shape and swimming traits (Pearson’s *r* = 0.71, p < 0.001; Spearman’s ρ = 0.72, p < 0.001). Because of its intensely bimodal distribution, however, correlations between the defensive trait LD1 and the other trait suites’ primary LD axes should be viewed skeptically. Removing G and Echo Lakes, which included most of the individuals included in the smaller peak in defense LD1’s distribution, reduced the strength of correlations between defense LD1 and the LD1 of other trait suites, including lowering the correlation with trophic LD1 to *r* = −0.06, p = 0.40. Even after removing these populations, the apparent relationships between defensive and other trait suites remained idiosyncratic.

Mantel tests between full-dimensional data of all trait suites revealed that there was no significant multivariate correlation between the defensive trait suite and shape (Pearson’s *r* = 0.033, p = 0.110; Spearman’s ρ = 0.037, p = 0.055), and a relatively weak correlation between defensive and trophic trait suites (Pearson’s *r* = 0.078, p = 0.005; Spearman’s ρ = 0.082, p < 2×10^-4^; Table S9). By far the strongest correlation shown in mantel tests was between trophic and swimming traits (Pearson’s *r* = 0.789, p < 2×10^-4^; Spearman’s ρ = 0.705, p < 2×10^-4^), and all other multivariate correlations had correlation coefficients less than 0.13. Analyzed using Spearman’s rank correlation to accommodate severe violations of the Pearson correlation’s assumption of normality, results of multivariate correlations of the defensive suite with other trait suites changed little. Although all trait suites did violate the assumption of multivariate normality as assessed using the Royston test for multivariate normality – as might be expected of samples drawn from multiple distinct populations – this assumption was violated most severely by the defensive trait suite (Royston’s H = 308.1, p = 7.9 x 10^-65^; H = 229.3, p = 6.9 x 10^-48^ when partially-plated individual from Spirit L. excluded).

### 2. Comparisons of within-lake correlation structures

Two-block PLS correlations between trait suites within populations show significant integration for each combination of trait suites in at least some lakes. Of these, the swimming x trophic suite tests show the most persistent trends (*r_PLS_* > 0.855 in all cases, *Z*_range_ = 2.36 – 7.01, p ≤ 0.001 in all lakes except Engineer: p = 0.011; Table S10). The defensive trait suite and shape had by far the least consistent, and typically weakest, relationship. Their *r_PLS_* correlation coefficients ranged from 0.33 (Wik Lake, *Z* = −1.71, p = 0.962) to 0.79 and 0.80 (Echo Lake, *Z* = 3.24, p = 0.006; and Long Lake, *Z* = 0.99, p = 0.362, respectively). The other pairs of trait suites mostly had clear integrative relationships, although the patterns of shared variation between defensive and trophic traits were also somewhat erratic between lakes.

Pairwise comparisons between lakes for difference of PLS effect size in each pair of trait suites revealed several cases of clusters of populations that share similar integration strengths when compared with other lakes, albeit typically with relatively moderate effect sizes, and none higher than *Z* = 3.89 (Figure S4). These groups do not clearly reflect populations’ position along the benthic-limnetic axis, and in some cases, lakes on opposite ends of the traditional benthic-limnetic axis share similar integration strengths. Watson and Wik Lakes, for instance, have quite similar patterns of integration between defensive and trophic traits, despite Watson fish being very benthic and Wik fish being very limnetic.

More revealing were the patterns of relative orientation of PLS axes between lakes (Figure S5). In the swimming x trophic PLS test, axes of correlation between these trait suites were strikingly parallel across all lakes, with the angles between trophic suite axes never exceeding 26° between lakes and the angles between swimming suite axes never exceeding 24°. Integration structures between swimming and defensive traits or shape also resulted in quite parallel PLS axes for the swimming suite, although the orientations of the defensive trait especially the shape axes were much more muddled in these cases. The integration patterns between some trait suites, in contrast, sometimes clustered the lakes into more or less poorly-defined groups, within which lakes shared similar PLS axis orientations and differed in similar ways from lakes in other clusters (Figure S5). However, the lakes composing these clusters were not necessarily the same between integration structures. Additionally and surprisingly, as was the case with integration effect sizes, the lakes in the PLS axis orientation clusters did not map clearly onto the fairly obvious, but apparently superficial, benthic-limnetic continuum. One of the clusters in the defensive block defensive x trophic suite PLS angle matrix included Wik and G Lakes, both of which are extremely limnetic, but also Tern Lake, the most benthic of any of the lakes surveyed.

### 3. Modularity of trait suites

The complete modularity test excluding shape showed that there is a statistically significant modular signal of relatively modest strength in every population except Engineer, which was only marginally insignificant, and differed only slightly from other populations (Table S11). However, the focused tests on individual trait suites revealed some patterns in modularity of trait suites. In particular, the defensive trait suite exhibited moderate to high levels of modularity in all lakes, with the effect being strongest in G, Wik, and Spirit Lakes, and weakest in Long, Finger, and Engineer (Table S12). When gill raker data is excluded from the trophic trait suite to increase sample size, the signal of modularity in defensive traits generally becomes even stronger, even exceeding *Z* = −10 in Wik and Spirit Lakes. Modularity in trophic traits was not consistent between lakes and exhibited a significant, moderately sized signal in four lakes (Corcoran, G, Long, and Spirit). Swimming traits were not modular in any population, and had consistently weak effect sizes. That swimming traits are not modular was not surprising given the importance of swimming performance in both feeding and avoiding predation.

The inclusion of shape in a complete modularity test lowered covariance ratios, and also slightly strengthened effect sizes of modularity (Table S13). When shape was considered in a focused test of modularity, the signal of modularity was significant in all lakes, and with strong effect sizes. Its inclusion in other focused tests reduced the effect size of the modular signal, particularly for the defensive trait suite, indicating shared variation between defensive traits and some components of shape (Table S14).

### 4. Responses to the environment

The surveyed lakes are mostly of small to moderate size (min. surface area = 2.7 ha, Ruth Lake – although Ruth was excluded from PCAs due to lack of Ca data; max. surface area = 135.7 ha, Finger Lake). However, these lakes exhibit a wide range of environmental characteristics. Lakes ranged from 2.1 (Corcoran) – 24.4 m (Wik) in maximum depth. All lakes had low to very low biomass of primary producers (max. Chl-*a* = 3.30 μgL^-1^, Finger Lake; min. Chl-*a* = 0.42 μgL^-1^, Wik Lake). The pH in the lakes ranged from circumneutral (pH = 6.6, Ruth Lake) to alkaline (pH = 8.8, Corcoran Lake), and Ca and conductivity were both very low in some lakes. The two lakes with the lowest Chl-*a*, Wik and G Lakes, also had the lowest Ca (1.3 and 0.6 mgL^-1^, respectively) and cond. (14.9 and 11.3 μScm^-1^, respectively). Jean Lake had the highest Ca (36.9 mgL^-1^), and Finger Lake had the highest cond. (239.0 μScm^-1^). Principal components biplots showing lakes and their relationships with environmental variables are presented in Figure 6, with PC loadings and eigenvalues in Tables S15-S16. See Appendix B for detailed results on foraging-related environmental variables.

**Figure 6.**
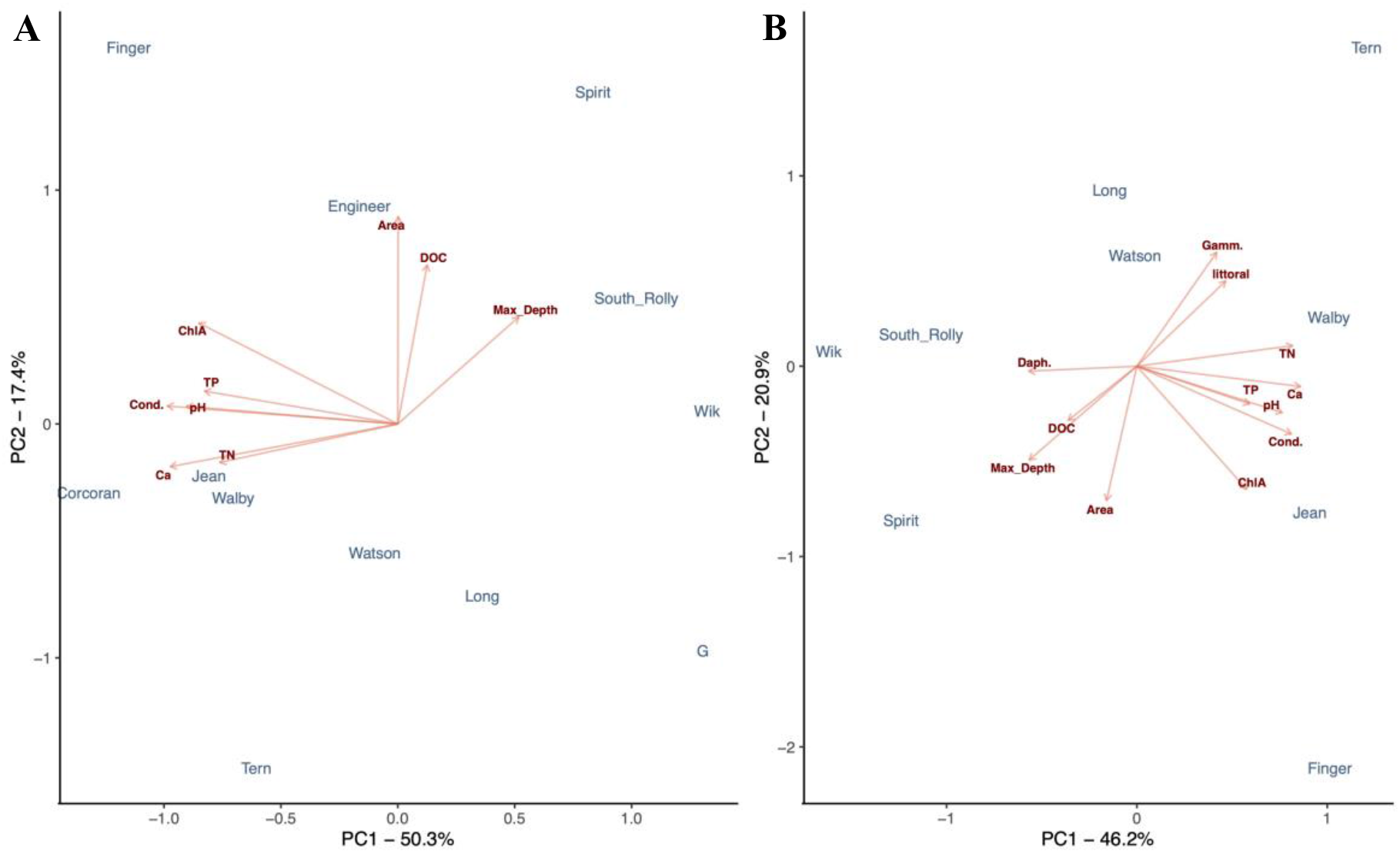
PCA biplots of lakes and environmental variables. Panel A shows physico-chemical environmental variables. Panel B shows a PCA biplot additionally including *Daphnia* abundance, Gammarid abundance and percent littoral area, all foraging-related environmental variables. Ruth Lake and Echo Lake are excluded from both panels because some environmental data were unavailable.

Two-block partial least squares regressions between trait suites and environmental variables did not reveal any significant correlations after p-values were adjusted for multiple comparisons (Figure 7; Table S17). However, physico-chemical environmental variables alone were correlated with stickleback defensive and swimming trait suites at levels only slightly higher than the p = 0.05 threshold. While the trophic trait suite was not strongly correlated with physico-chemical characteristics of the lakes, this association was strengthened substantially by the inclusion of environmental variables related to foraging (*Daphnia* sp. abundance and gammarid density, and % littoral area), even though this reduced the number of lakes for which data was available to only eight lakes, and the correlation remained statistically insignificant. Axis loadings and singular values for all PLS tests are shown in Tables S18-S21.

**Figure 7.**
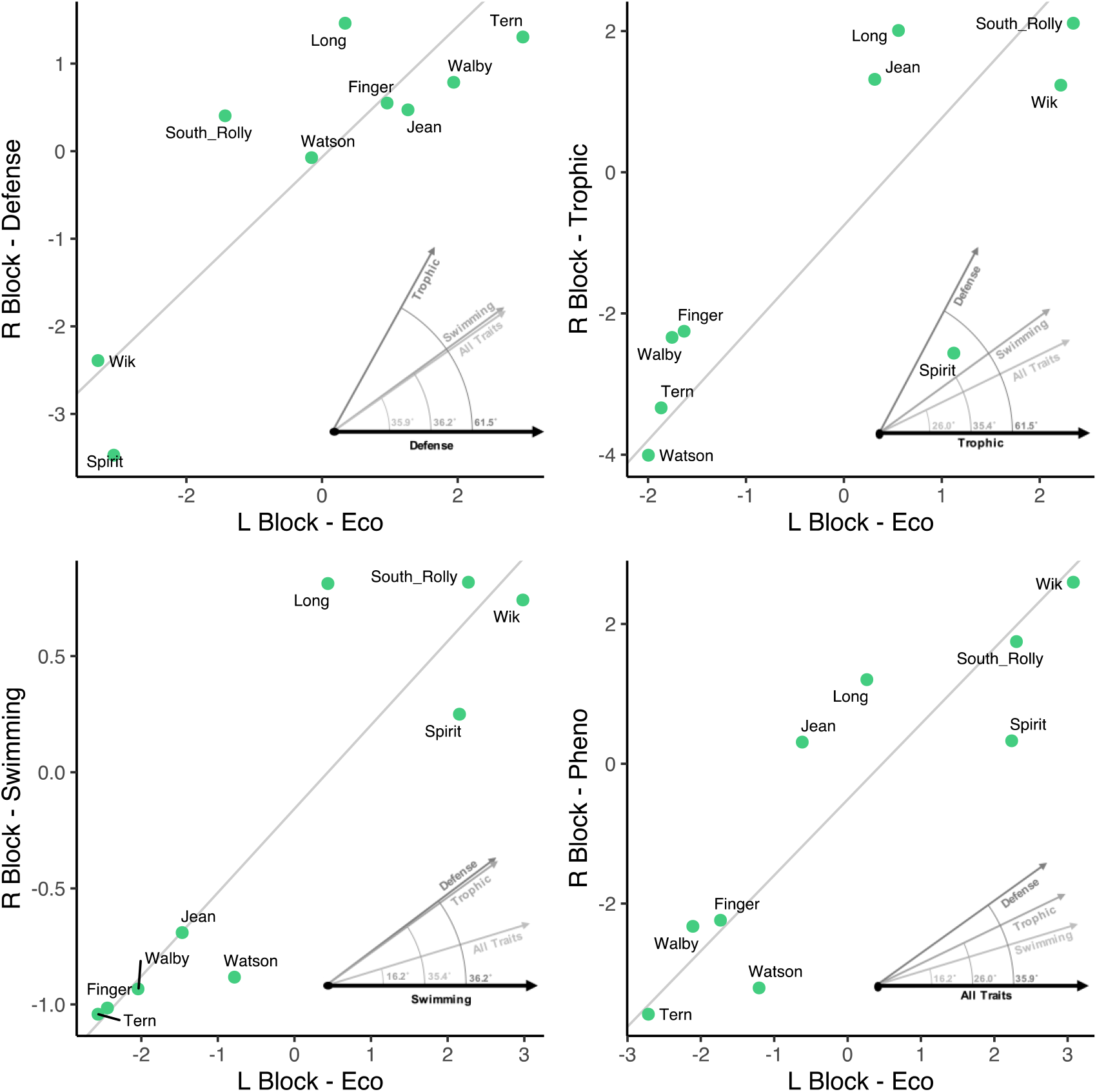
Plots of two-block PLS axes of stickleback traits against all environmental characteristics. PLS axes are plotted by trait suite, with defensive traits at the top left, trophic traits at the top right, swimming traits on the bottom left, and traits from all three of those suites at the bottom right. Inset diagrams show angles of the plotted trait suite’s primary vector through environmental space relative to the vectors of other trait groups through environmental space. The X axes of all plots represents the major axis of phenotype-ecology covariation through the environmental data block, and the Y axes represent the same through the phenotypic data block.

When the angles of the primary environmental vectors were calculated between 2B-PLS tests against different trait suites, angles involving the stickleback defensive trait suites were consistently largest. The defensive trait suite angles, especially the angles between the defensive trait suite environmental axis and the trophic trait suite environmental axis, indicate that the major axis of divergence in defensive traits among lakes occurs along an environmental gradient that is divergent from those of the other traits (Figure 7; Table 3). These angles also widen greatly with the inclusion of foraging-related environmental data, further emphasizing the distinction between the axis of divergence in defensive traits and the more congruent trophic and swimming traits.

**Table 3.**
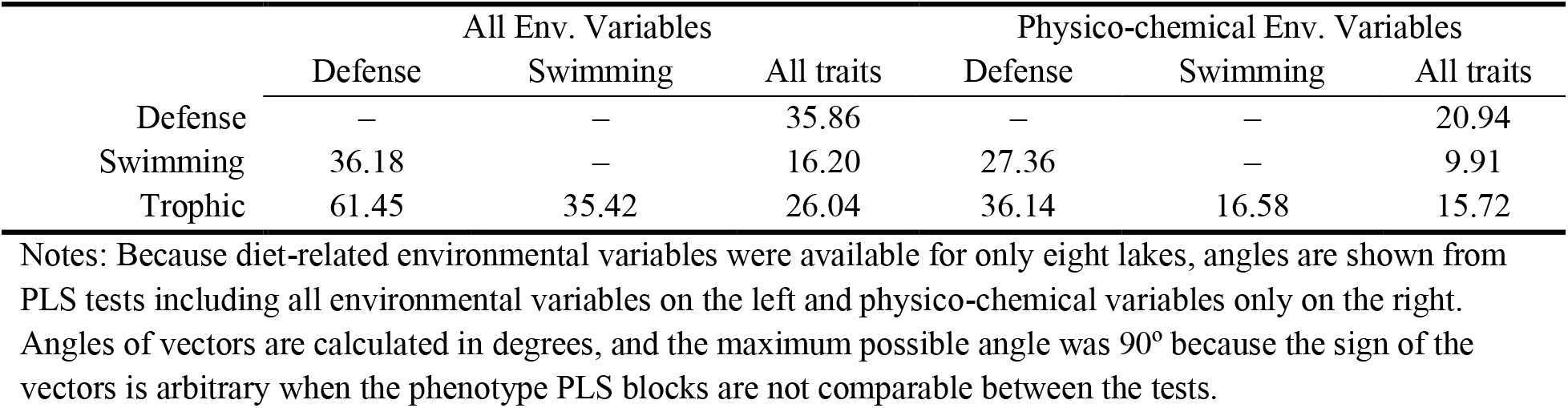
Angles of environmental vectors describing the axis of maximum covariance with traits in two-block PLS tests in relation to environmental vectors in 2B-PLS tests involving other trait suites.

## Discussion

Even on short timescales, stickleback populations can adapt in response to selection on several trait types simultaneously (Marques et al. 2018). However, because traits often covary, even when their functional relationships are not obvious or are indirect (Barrett et al. 2008; Bjærke et al. 2010), it has been difficult to establish whether populations diverge along a low-dimensional space following a genetic line of least resistance (Schluter 1996), or diverge in a higher-dimensional space as a consequence in response to multifarious selection on functionally disparate traits (Nosil et al. 2009).

We show that divergence is not a unidimensional process, but instead takes place along several axes both within and across trait suites under selection from distinct sources. This permits multiple possible evolutionary trajectories through phenotype space depending on the selection pressures experienced. While some within-lake patterns of integration between suites of traits are consistent across lakes, others vary considerably in both strength and direction between populations. Consistency and extent of trait suite modularity also varies by trait suite. The major axes of variation in the swimming and trophic trait suites also correspond to PLS axes of similar environmental variation. However, these diverged from the environmental axis along which defensive traits varied, indicating potential tradeoffs in which the optimum phenotypes of different trait suites diverge.

The most multidimensional patterns of divergence existed in body shape data and in the trophic trait suite. For the defensive trait suite, there was low dimensionality of divergence, but this was partially due to the finding that divergence along LD1, dominated by pelvic spine length, was so great that it swamped other dimensions in the calculation, rather than the lack of divergence along other axes. Importantly, while primary axes of divergence were strongly correlated between trait suites, trait suites were less strongly correlated with one another when they were considered as multivariate trait matrices. These patterns illustrate that lower-magnitude components of divergence may be more susceptible to selective pressures not shared between trait suites. This conclusion is also supported by the persistent, but relatively small-effect overall signal of modularity between trait suites.

Patterns of integration between trait suites often vary in strength and direction between populations, demonstrating that important axes of divergence not only exist in the trait values themselves, but also in the ways that trait suites are related to one another. This could potentially influence how trait suites respond to the environment via both adaptation and plasticity. In addition to our conclusions about the dimensionality of divergence within and between trait suites, we established that the defensive trait suite was the most independent of the trait suites we considered. This was evident from the inconsistent orientation of its axes of covariation with the other trait suites, and also from its high levels of modularity in all lakes, when considered independently of shape. 2B-PLS tests of trait suites against environmental variables further showed that the environmental gradient along which defensive traits varied diverged in its orientation from the environmental gradients along which the other trait suites varied. Trophic and swimming traits varied much less independently than defensive traits, although some lakes did exhibit modularity in trophic phenotypes. The axes along which they were correlated with one another were also very similar among lakes.

### General Conclusions

Three notes of caution regarding dimensionality of divergence should accompany our findings. 1) The range of habitats that we have included, while ecologically diverse for lakes with stickleback populations, represent only a small subset of the range of habitats in which stickleback are found, and consequently, probably underestimate the magnitude, and likely the dimensionality, of divergence among stickleback populations more broadly. There are also anadromous marine stickleback populations, as well as stream populations (Lavin and McPhail 1993; Bell and Foster 1994; Seebacher et al. 2016). Inclusion of either or both population types would likely have substantially increased the dimensionality calculated for the defensive trait suite, and the shape and swimming trait suites, respectively. In the former case, this is because threespine stickleback have three major lateral plate morphs: low, partial, and complete, and while low- and partial-plated morphs are common in freshwater, marine populations are typically made up almost entirely of completely-plated morph (Hagen 1967; Bell and Foster 1994; Barrett et al. 2008). Of the stickleback represented in this study, there was only one single partial-plated individual (from Spirit Lake), and the remainder were low-plated. This means that the fish we sampled only represent a small band of the possible variation in this trait, and divergence in defensive traits was determined primarily by spine length in our data. In the latter case, populations of stickleback inhabiting streams typically have trophic adaptations suited to benthivory, as benthic invertebrates are more reliably available as prey items in most streams than zooplankton (Berner et al. 2008; Berner et al. 2009). They also typically have a deeper body shape that facilitates navigating the spatially heterogeneous flow regimes present in many streams (Hendry and Taylor 2004; Izen et al. 2016). The specificity of our study to lacustrine populations likely does not adequately represent the dimensionality of trait divergence across the entire species or the diversity of integration structures among trait suites.

2) Another important consideration is that our results seem to indicate that both effective dimensionality and variability in the orientation of integration structure PLS axes generally increase with the number of traits under consideration, and this observation was supported by our rarefaction-like analysis in the appendices. This may be intuitive – after all, our swimming trait suite, which is composed of only three traits, cannot possibly have more than three effective dimensions. Neither can the number of effective dimensions of divergence exceed one fewer than the number of populations under study. Nevertheless, the intrinsic dimensionality (Del Giudice 2020) of divergence seems to not often be considered when authors assess whether an adaptive radiation is constrained to a line of least resistance (Schluter 1996). It is therefore important that researchers, when testing and making claims about the dimensionality of adaptation or divergence, measure broad ranges of traits of differing functions in populations of diverse habitat types, unless the intention is to test a narrow question relating to patterns of adaptation in a particular suite of traits. A rarefaction-like procedure similar to the one we have implemented in the appendices could help fewer researchers avoid the hazards involved in calculating effective dimensionality.

3) Finally, we advise other researchers to be conscious not just of the dimensionality of divergence in their systems, but also of the *magnitude* of divergence. The measures of effective dimensionality we have employed represent relative strength of the vectors of divergence to one another, but do not describe the extent to which they have diverged. In our case, although the populations we sampled are visibly divergent without measurement of any phenotypes, the large magnitude of the difference between the populations with pelvic spines and those without pelvic spines dwarfed other dimensions of divergence. Spine length reduced effective dimensionality of traits, and especially of the defensive trait suite, even when differences in other traits among populations were substantial, and this effect can be seen in Figure B1. Consideration of the magnitude is also necessary for measurements of divergence to be useful as adaptive responses to the environment (Stuart et al. 2017; Runquist et al. 2020), or when assessing the effect of divergence on an environment.

Our findings suggest that divergence among populations is not typically constrained by the axis of greatest covariation, even within groups of traits that might be expected to respond to similar selection pressures. This conclusion is consistent with the known examples of highly divergent conspecific sympatric morphs even within a single habitat – especially among salmonids (Skúlason et al. 1989; Jónsson 2000; Chaverie et al. 2016; Muir et al. 2016; Arostegui and Quinn 2019; Chaverie et al. 2020), and is incompatible with a simple singly-peaked or – ridged fitness landscape. Although evidence from recent post-glacial radiations – like some sticklebacks and the *Salvelinus* species – are particularly compelling because of the relatively short period of time available for the reorientation of **g_max_** in a low-dimensional model of divergence, other radiations also provide support for multi-axis divergence that is inconsistent with low-dimensional adaptation constrained by genetic covariances (Velasco and Herrel 2007; Hulsey et al. 2019; Levin et al. 2021). Further investigations into divergence as it progresses could further elucidate this process. This is especially true of the role of plasticity, which Noble et al. (2019) have shown is often aligned with **g_max_**, which would be expected to facilitate low-dimensional divergence, in spite of our results.

The implications of these findings are important to consider when attempting to assess the repeatability and predictability of evolution in nature, one of the central themes of evolutionary biology. The possibility of adaptation in a high-dimensional space certainly makes predicting the evolutionary effect of an environmental condition in any particular instance difficult, even with high-quality data about multiple aspects of the environment and when the relatedness among populations is known (Stuart et al. 2017; Yong et al. 2020). However, the potential to adapt along a number of axes may increase resilience to environmental changes, allowing a population to adapt to the change in the context of its specific habitat conditions (Reisch et al. 2015), rather constraining it to a single axis that may be perpendicular to the direction of selection. Because the dimensionality of potential divergence is likely very taxon-specific, further investigations into the genomic mechanisms and the role of plasticity through time in diverging populations will be necessary to develop useful predictive models for the evolution of species that are vulnerable, ecologically important, or important to human health.

## Acknowledgements

This work was conducted under Alaska DF&G permit #SF2018-124 and Kenai National Wildlife Refuge permit #2018-Res-GHaines-6231 for stickleback collections, and Alaska DF&G permit #SF2018-001 for macroinvertebrate collections. The limnological research was funded by a NSERC Discovery grant to AMD, and other field work was supported by an NSERC Discovery grant and CRC funding to APH. We thank Stephanie Guernon and Maxime St Martin for zooplankton analyses and Nathan Juillet for macroinvertebrate analyses at UQAM. Many thanks to Mike Bell and Rob Massengill for their guidance and knowledge of local stickleback populations, and to Blake Matthews for comments on the manuscript. Thanks also to Bailey Feddersen, Capucine Lechartre, Rebecca Pahulje, Ismail Ameen, and Greig Hospes for assistance with data collection, and Victor Frankel, Sarah Sanderson, Mariane Daneau-Lamoureux, and Michelle Packer for assistance in the field.

## Appendix A: Supplementary Methods

**Table A1.** Tests in the Methods, and the Panels of Figure 1 to which they Correspond

| *Question* | *Test* | *Figure 1 panel(s)* |
| --- | --- | --- |
| *1) To what extent do lacustrine stickleback diversify along a single axis of divergence among populations, and what is the effective dimensionality of that divergence?* | Dimensionality of traits | A1-A2 |
|  | Dimensionality of trait suites | Analogous to A1-A2 |
|  | Correlations between trait suite primary axes | B1-B2 |
|  | Multivariate correlations between trait suites (Mantel tests) | B1-B2 |
| *2) To what extent do lakes share common within-lake correlation structures between trait modules?* | 2Block-PLS tests between trait suites within populations | C1, C3 |
|  | Angle calculations of 2B-PLS axes trait suites between populations | C1-C2 |
|  | Z-tests for differences in strength of integration structures between populations | Not shown. This test focuses on strength of integration, not orientation |
| *3) To what extent do functional trait suites operate as adaptive modules?* | modularity tests | C2-C3 |
| *4) To what extent are different environmental variables, or groups of environmental variables, associated with variation in all traits and in the trait suites?* | PCA | Not shown |
|  | 2B-PLS between full trait data and environment | D1-D2 |
|  | 2B-PLS between trait suites and environment | D3 |
|  | Angle measurements between environment vectors for the trait suites | D3 |

### The Use of Linear Discriminant Analysis for Calculation of Effective Dimensionality

We use LDA, which places an emphasis on the differences between groups relative the variance within groups, rather than Principal Components Analysis (PCA) to identify axes of divergence because the primary subject of our analyses is the dimensionality of divergence between populations, not simply differences between group means. Typically dimensionality of divergence is studied using eigenvectors calculated from population means (as in Schluter 1996), but because of differing magnitudes and orientations of trait variances within populations, this does not capture the axes along which populations are actually most divergent from one another. As such, the use of eigenvectors of population means represents the process of phenotypic divergence – which can be constrained by the evolvability of traits, lineage history, and the ability of variation to persist in populations due to population size and balancing or disruptive selection – as equally possible in all directions. Effectively, it is a mathematical convenience that is agnostic towards which constraints on phenotypic change are most important (which may differ depending on study question).

### Comparison of Effective Dimensionalities of Trait Suites with Differing Numbers of Traits

Effective dimensionality of data matrices depends in part on the number of traits included in the data. For instance, the data we include in our swimming trait suite includes only three traits, and it therefore not possibly have an effective trait dimensionality higher than three. As a consequence, it was necessary to develop a method that would enable comparing trait dimensionality between trait suites at different numbers of traits. We accomplished this by implementing a rarefaction-like procedure in which traits were randomly drawn without replacement from the trait matrix for each trait suite. After each draw, an LDA was performed with the sampled traits, and dimensionality of divergence calculated using the Cangelosi and Goriely / Roy and Vetterli index presented in Table 1. This was done for each random draw of a new trait, until all traits in the trait suite had been drawn. For each trait suite, as well as for all the non-shape traits, this process was repeated for 50 permutations. A loess regression was then fitted to each trait suite to show change in the mean effective dimensionality as more traits are accumulated.

### Limnological Conditions Detailed Methods

Lake physico-chemical characteristics were measured at the deepest point of each study lake in June 2018 and June 2019. For most lakes, these data were an average of 2018 and 2019 values, but not all lakes were surveyed in both years, so data from Jean and Corcoran Lakes are from 2018 only, and data from Engineer and Ruth Lakes are from 2019. pH and conductivity (cond; μS.cm^-2^) were measured using a Professional Plus Model YSI multi-parameter sonde (model 10102030; Yellow Springs Inc.). We collected epilimnetic water samples to quantify dissolved organic carbon (DOC; mgL^-1^), total nitrogen (TN; mgL^-1^), total phosphorous (TP; μgL^-1^), chlorophyll-*a* (Chl-*a*; μgL^-1^), and dissolved calcium (Ca; mgL^-1^). Water calcium (Ca) was collected in 2018 only, except for Echo and Engineer Lakes, for which we used data from a 1990 survey (Jones et al. 2003). DOC, TN, TP, and chl-*a* samples were analyzed at the GRIL-Université du Québec à Montréal (UQAM) analytical laboratory. DOC concentrations of 0.45-μm filtered samples (surfactant-free membrane filters) were measured after acidification (phosphoric acid 5%) followed by sodium persulfate oxidation using a 1010 TOC analyzer (O.I. Analytical, College Station, TX, USA). Total Nitrogen (TN) was analyzed with a continuous flow analyzer (OI Analytical Flow Solution 3100 ©) using an alkaline persulfate digestion method, coupled with a cadmium reactor, following a standard protocol (Patton and Kryskalla 2003). Total Phosphorus (TP) was measured spectrophotometrically on the same machine by the molybdenum blue method after persulfate digestion (Griesbach and Peters 1991). Chl-*a* was quantified by passing samples through glass fiber filters (Whatman GF/F), extraction of the chl-*a* in hot ethanol and measuring the chlorophyll spectrophotometrically on a BiochromUltrospec” 2100 pro with a 10 cm quartz cuvette (Wintermans and de Mots 1965; Sartory and Grobbelaar 1984). Water Ca samples were analyzed with a Thermo ICAP-6300 Inductively Coupled Argon Plasma – Optical Emission Spectrometer (ICP-OES) following protocols described by US EPA (1994) at the University of Alberta Biogeochemical Analytical Service Laboratory (U of A – BASL; Edmonton, Alberta, Canada). Surface area and maximum depth data for lakes was obtained from previous ADF&G surveys.

### Diet-Related Environmental Data

To estimate *Daphnia* abundance in the pelagic zone of each lake, crustacean zooplankton were collected by whole water column vertical tows with a 35-cm diameter Wisconsin net from 0.5 m off the bottom to the surface. The zooplankton were sampled from four sampling stations along an open-water transect across each lake. The crustacean zooplankton were anaesthetized with bromoseltzer, preserved with 95% ethanol, and they were brought back to the laboratory at the UQAM for identification. These four samples were subsequently pooled to a single sample per lake for identification and enumeration in the laboratory. The crustacean zooplankton were identified to species level with a high-resolution dissecting microscope (6.3-126x; SZ2-IL-ST; Olympus SZ) and species were enumerated for the number of individuals per L. Crustacean zooplankton were counted using a protocol that targeted mature individuals that could be identified unambiguously to species, as well as to detect rare species (Mack et al. 2012). Subsamples (10 mL) were taken from a standardized 50 mL sample volume, until at least 200 individuals of the most abundant species were counted (excluding copepodids and nauplii), or a total of 1000 individuals were counted (excluding copepodids and nauplii), or the total sample was counted. Taxonomic keys for *Daphnia* sp. included Brooks (1957), Taylor and Hebert (1993), Witty (2004), Thorp and Covich (2010), and Haney et al. (2013).

To estimate gammarid density in the littoral zone of each lake, nearshore littoral macroinvertebrate communities were collected with a D-frame kick net at four sampling stations located around the lake in the 2018 survey, and eight sampling stations in the 2019 survey. Sampling was performed by the sweep method (“Kick & Sweep”) with a 500 μm “D-net” as recommended by the Ontario Benthos Biomonitoring Network (Jones et al. 2007) on a surface of approximatively 2 m^2^. Samples were concentrated with a 500-μm sieve, preserved in 95% ethanol, and they were brought back to the laboratory at UQAM for identification. Identification was done up to the Family level following taxonomic keys (Merritt et al. 2008; Moisan 2010). All macroinvertebrate samples were identified using SZX10 stereo microscopes (Olympus) with varying magnification (x6.3 – x10). For each sample, 100mL sub-sample was taken and counted until the 100th individual was reached. If the 100th individual was not reached within the initial 100 mL, an additional 100 mL was counted. When the 100^th^ individual was reached, the remaining part of the sub-sample was counted, and the total sub-sampled volume calculated. A ratio was then calculated between the total sub-sampled volume and the sample total volume to estimate the taxon-specific abundance of macroinvertebrates. The density of gammarid individuals per m^2^ was averaged for the 4 samples collected per lake to obtain total average gammarid density per lake in the littoral zone.

### Percent Littoral Area

To calculate the proportion littoral area for each lake, we used bathymetric maps that were available through the Alaska Department of Fish & Game. For each of the maps’ contours, we calculated the area by measuring the area of the map image bounded by the contour in ImageJ (Rasband 1997-2018). This was then converted to area using a scale indicated on the map, a total surface area indicated on the map, or, when no indication of scale was indicated on the map, the total surface area from previous surveys used in the environmental data. When maximum depth was not indicated on a map, the maximum taken during field surveys was used, unless there was a contour on the map below this depth. For Tern Lake, the only bathymetric map available used transects, rather than contours, to indicate depth. In this case, transect intervals were assumed to be proportional to area, although this likely resulted in an overestimate of the area of the deeper portions of the lake. The total area of Tern Lake as indicated on the documents accompanying the bathymetric map was also much smaller than the area used in the environmental database, likely as a result of changing lake morphology since the map was made in 1970.

**Figure A1.**
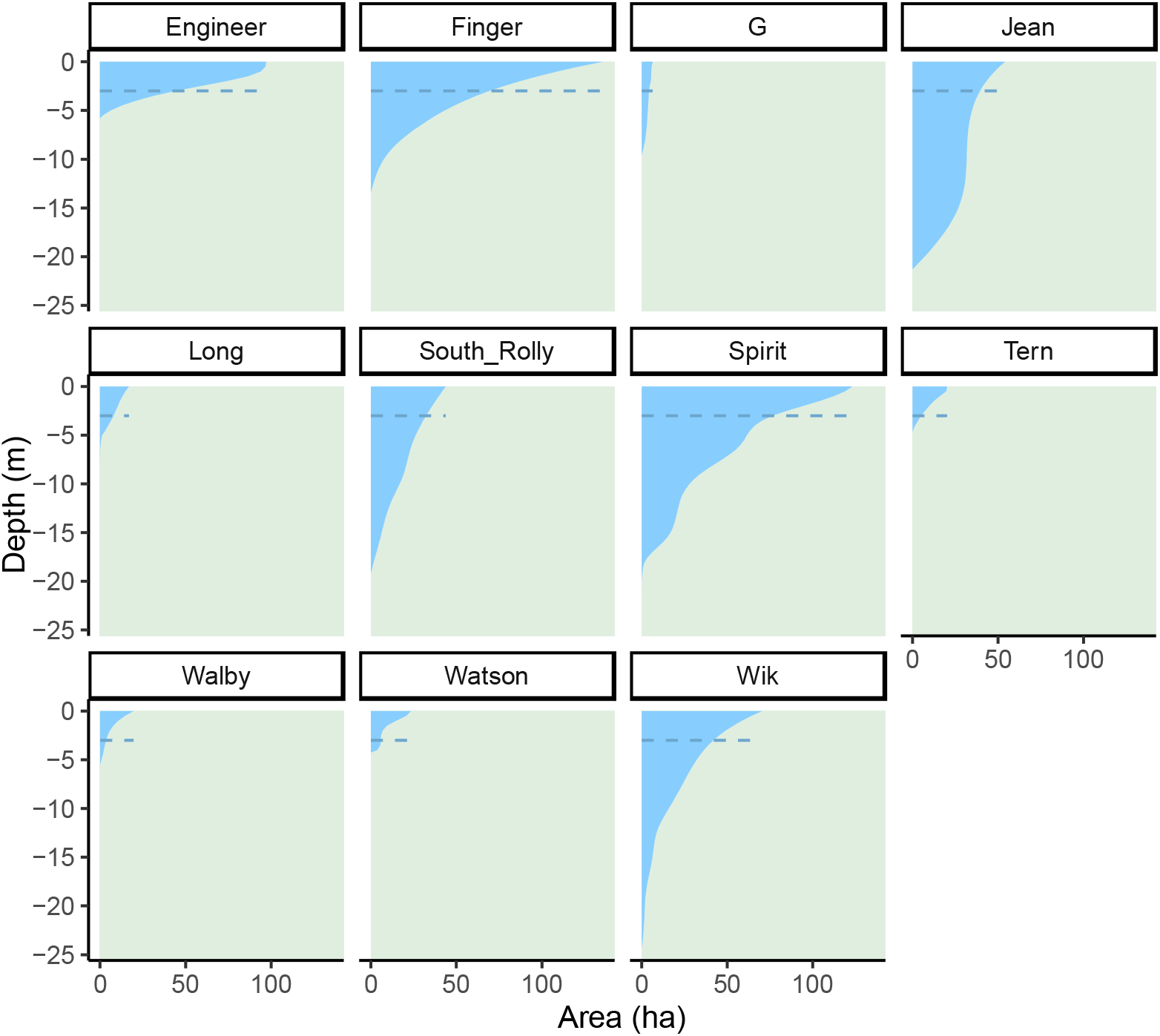
Depth profiles of lakes for which bathymetric data were available. The profiles shown are depth-area curves estimated from available data using monotonic splines. The dashed horizontal line represents a depth of 3 meters, which was used as the limit of the littoral zone for calculations of percent littoral area.

Because bathymetric maps from the different lakes had depth contours marked at inconsistent intervals, it was necessary to estimate depths at common intervals across all lakes. Therefore, after collecting depth profile data from bathymetric maps, we computed a monotonic spline for each lake using surface area as a response variable to depth. The splines used the method of Hyman (1983) as implemented in *R*’s *stats* package (R Core Team 2019), to avoid the calculation of increases in area with increasing depth. The littoral area was defined as the area of the lake having a depth of three meters or less, and the percent littoral area was calculated as 1-Area_Depth>3m_ / Area_Total_ where the area with a depth > 3 m was interpolated from the spline.

## Appendix B: Supplementary Results

### Comparison of Effective Dimensionalities of Trait Suites with Differing Numbers of Traits

The permutation procedure implemented to compare effective dimensionalities of differing trait groups showed that at low numbers of traits, the effective dimensionality of swimming and trophic trait suites is slightly lower, but similar, to that of shape (Figure B1). Divergence in the defensive trait suite is consistently much lower in effective dimensionality, as is divergence in the full non-shape trait data. However, this is primarily due to the inclusion of pelvic spine length, which results in the clear decline in divergence dimensionality in a permutation once it is included as a trait, an effect visible in the Defensive and Full data in Figure B1. Prior to inclusion of this trait in a permutation, the effective dimensionalities for both the defensive and full non-shape data are comparable to those of shape, trophic, and swimming trait suites.

**Figure B1.**
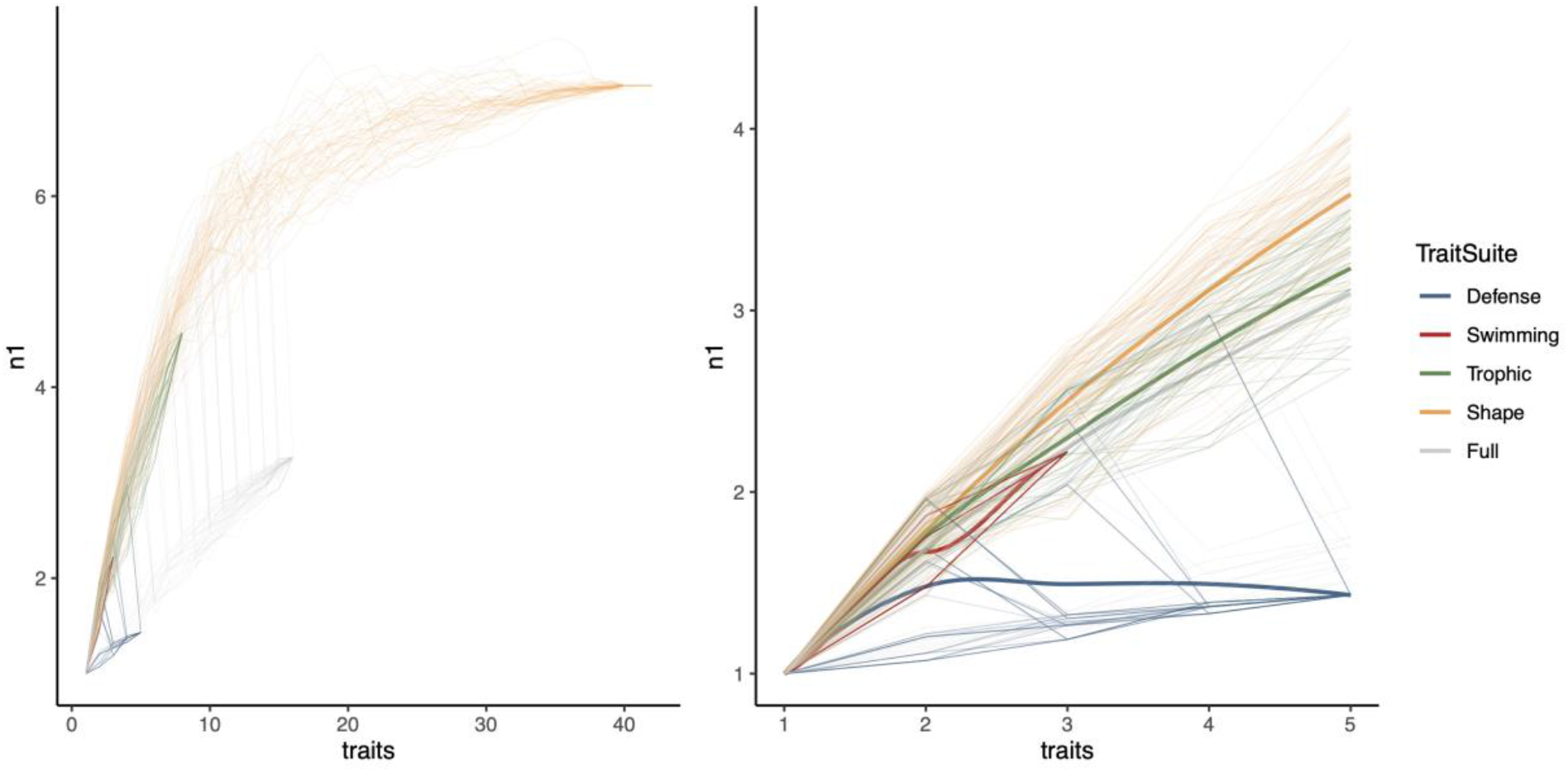
Effective dimensionality of divergence for each trait suite, as well as the full non-shape phenotype data, with the inclusion of different numbers of traits. Thin lines represent 50 permutations of randomly sampling traits until all traits are included in the dimensionality calculation of LDA vectors. The left panel includes complete data from the permutation procedures, and the right panel shows only the first five traits in the permutations along with loess regressions showing the mean effective dimensionality for each trait suite at a given number of traits.

### Diet-Related Environmental Data

Several patterns emerged from diet-related environmental data. For the *Daphnia* abundance, lakes clustered into two groups: Watson, Wik, and Spirit Lakes had over 5 *Daphnia* individuals L^-1^, and all other lakes, including Corcoran and Ruth (which are not plotted due to the absence of littoral area data), had less than 0.15 *Daphnia* L^-1^ (Figure B2). Most lakes had gammarid abundance less than 10.5 individuals per m^2^, with Long Lake having 10.5 individuals per m^2^, and Wik Lake having the fewest, with 1.0 individuals per m^2^. Tern Lake, however, had the highest abundance of gammarids, with 22.8 individuals per m^2^. Of the lakes with the highest percent littoral area – Tern (75.0%), Watson (75.6%), and Walby (82.1%) – Tern and Watson fish correspondingly appeared at the “benthic” extreme on the primary trophic trait LDA axis, although Walby Lake was more moderate in trophic morphology.

Interspecific competition likely played a role in the unusual prey abundance data for Watson Lake, which has threespine stickleback morphologies that we would expect of a benthivorous population, despite the abundance of *Daphnia* and paucity of gammarids. Of all the lakes sampled, Watson Lake was the only lake with a large ninespine stickleback (*Pungitius pungitius*) population. Wik Lake was the only other lake in which any ninespine stickleback were caught in 2018, but this was only a single individual. In a subsequent 2019 survey of Wik Lake, no ninespine stickleback were captured. State records also indicate that the two lakes immediately downstream of Watson, which are connected by short lengths of stream easily passable by fish, also have abundant sizeable populations of rainbow trout (*Oncorhynchus mykiss*) and longnose sucker (*Catostomus catostomus*) (ADF&G 2015). These species all likely compete with threespine stickleback for benthic prey, such as gammarids, and although we did not have sufficient data to formally test this the eco-evolutionary effects of competition here, this is a potentially fruitful avenue of future work.

**Figure B2.**
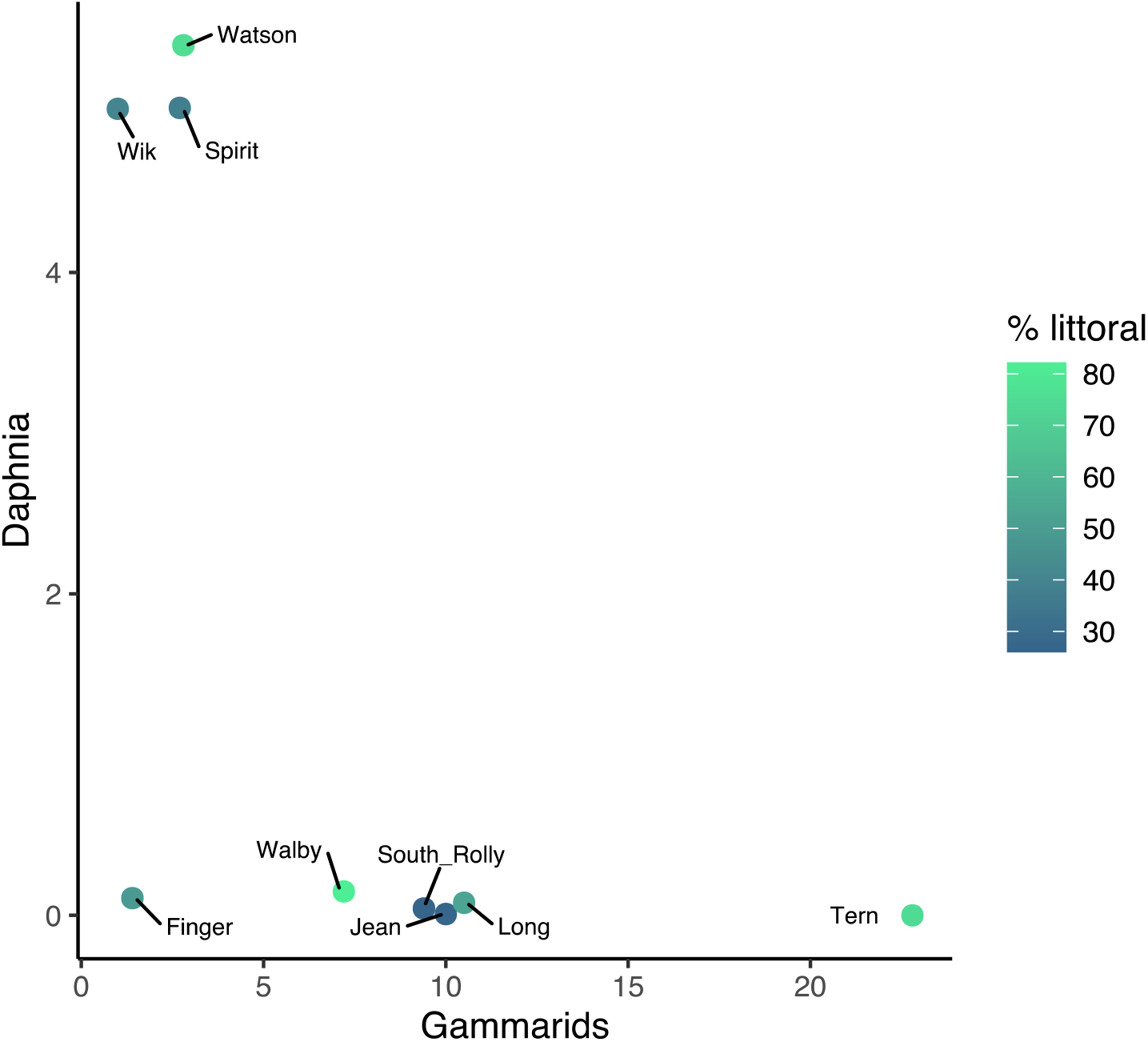
Density of *Daphnia* and Gammarids – important crustacean prey items for zooplanktivorous and benthivorous stickleback, respectively. *Daphnia* abundance is 2 shown as individuals per L, and Gammarid abundance is shown as individuals per m. Color scale indicates the proportion of each lakes surface area shallower than 3 m.

### Correlations of Environmental Variables with Phenotypes

Spearman’s correlations revealed that a large number of traits are associated with Calcium, pH, and conductivity, which are highly correlated environmental characteristics in these lakes (Figure B3). These traits include plate number and spine length, which are expected to be associated with are positively associated with Ca, pH, and conductivity, due to the dependence of these skeletal features on available calcium (Bell et al. 1993; Klepaker et al. 2013; Klepaker et al. 2016). These environmental characteristics are also negatively associated with trophic and swimming trait values suited to more benthic lifestyles. Nutrients have differing associations by trait suite, with phosphorus being negatively associated with trophic traits and nitrogen being positively associated with pelvic spine length and lateral plate number. Plate number was also the most strongly associated trait with the abundance of both *Daphnia* (negatively), and gammarids (positively), important limnetic and benthic prey, respectively, suggesting a kinematic cost of high-platedness for foraging on limnetic prey consistent with the cost of high-platedness on fast-start swimming (Bergstrom 2002).

**Figure B3.**
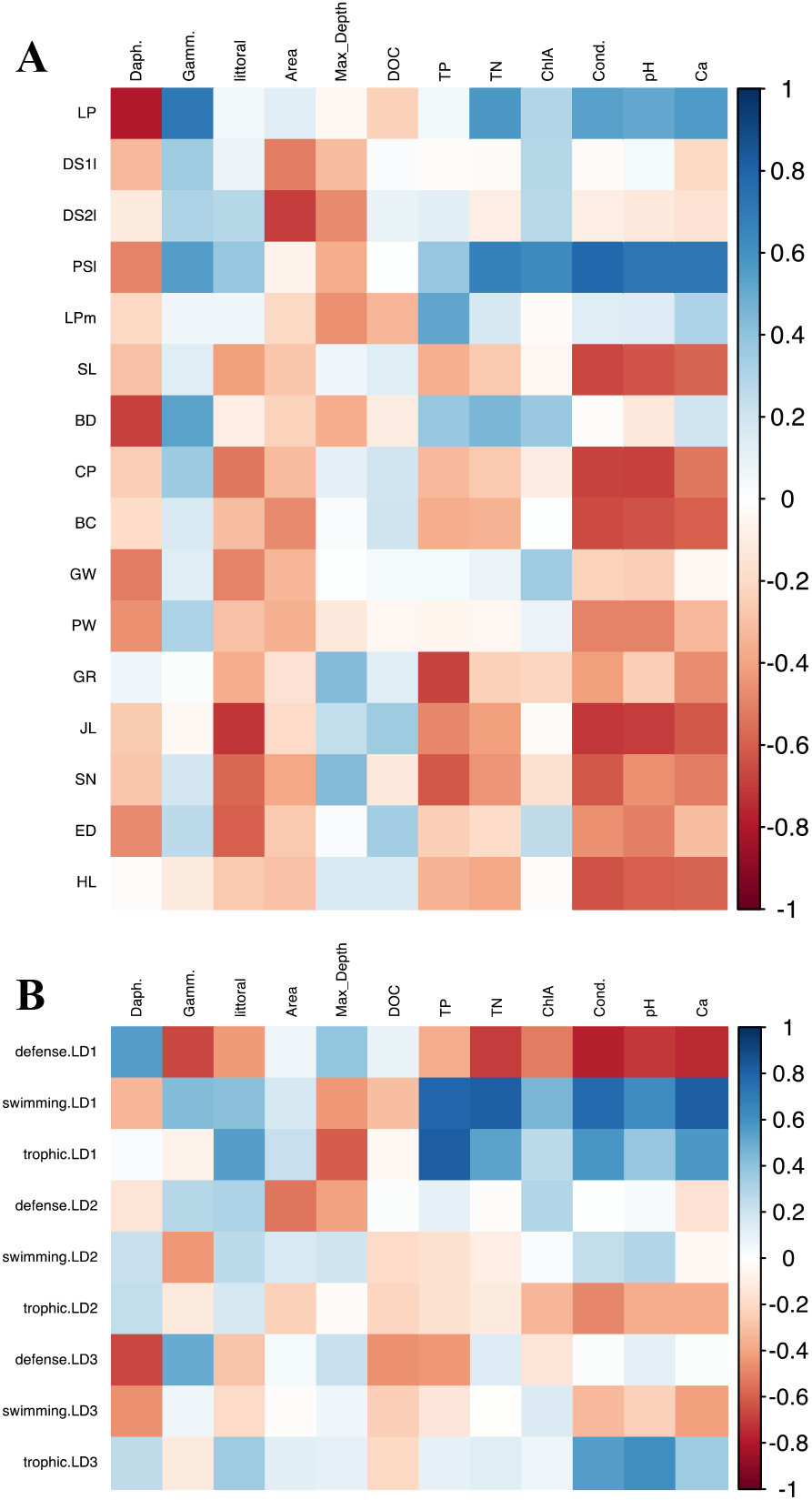
Spearman’s correlations between environmental variables and stickleback traits (A) and trait suite discriminate axes (B). Color scale indicates Spearman’s *ρ*.

In terms of the trait suite discriminant axes, the primary axis of defensive trait divergence is negatively associated with nutrient and Ca ion availability, as well as benthic prey abundance. The primary axes of trophic and swimming trait divergence are both strongly associated with lake depth and DOC concentrations, with the more “benthic” ends of these axes associated with shallow, low DOC lakes. They are associated with different prey types, however, with more “benthic” swimming values negatively associated with *Daphnia* abundance, and more “benthic” trophic values counterintuitively negatively associated with gammarid abundance. Despite the previously noted correlations in the LD axes, therefore, their environmental associations make clear some incongruities in the environmental correlates of divergence among trait suites.

## References

Adams, D., M. Collyer, and A. Kaliontzopoulou. 2019. geomorph: Geometric Morphometric Analysis of 2D/3D Landmark Data *in* D. Adams, ed, CRAN.

Adams, D. C. 2016. Evaluating modularity in morphometric data: challenges with the RV coefficient and a new test measure. Methods Ecol Evol 7:565–572.

Adams, D. C. and M. Collyer. 2019. Comparing the strength of modular signal, and evaluating alternative modular hypotheses, using covariance ratio effect sizes with morphometric data. Evolution 73:2352–2367.

Adams, D. C. and M. L. Collyer. 2016. On the comparison of the strength of morphological integration across morphometric datasets. Evolution 70:2623–2631.

ADF&G, S. F. D. 2015. The Alaska Lake Database (ALDAT). Alaska Department of Fish & Game.

Agrawal, A. F. and J. R. Stinchcombe. 2009. How much do genetic covariances alter the rate of adaptation? Proc. R. Soc. B. 276:1183–1191.

Alfaro, M. E., D. I. Bolnick, and P. C. Wainwright. 2005. Evolutionary consequences of many-to-one mapping of jaw morphology to mechanics in labrid fishes. The American Naturalist 165:E140–154.

Arnold, S. J. 1992. Constraints on phenotypic evolution. Am. Nat. 140:S85–S107.

Arostegui, M. C. and T. P. Quinn. 2019. Reliance on lakes by salmon, trout and charr (*Oncorhynchus*, *Salmo* and *Salvelinus*): An evaluation of spawning habitats, rearing strategies and trophic polymorphisms. Fish Fish. 20:775–794.

Barrett, R. D. H., S. M. Rogers, and D. Schluter. 2008. Natural selection on a major armor gene in threespine stickleback. Science 322:255–257.

Bell, M. A. 1981. Lateral plate polymorphism and ontogeny of the complete plate morph of threespine sticklebacks *(Gasterosteus aculeatus*). Evolution 35:67–74.

Bell, M. A. 1987. Interacting evolutionary constraints in pelvic reduction of threespine sticklebacks, *Gasterosteus aculeatus* (Pisces, Gasterosteidae). Biol. J. Linn. Soc. 31:347–382.

Bell, M. A. 2001. Lateral plate evolution in the threespine stickleback: getting nowhere fast. Genetica 112:445–461.

Bell, M. A. and S. A. Foster. 1994. Introduction to the evolutionary biology of the threespine stickleback. Pp. 1–27 *in* M. A. Bell, and S. A. Foster, eds. The evolutionary biology of the threespine stickleback. Oxford University Press, New York, NY.

Bell, M. A., G. Ortí, J. A. Walker, and J. P. Koenings. 1993. Evolution of pelvic reduction in threespine stickleback fish: a test of competing hypotheses. Evolution 47:906–914.

Benjamini, Y. and Y. Hochberg. 1995. Controlling the false discovery rate: a practical and powerful approach to multiple testing. Journal of the Royal Statistical Society: Series B (Methodological) 57:289–300.

Bentzen, P., M. S. Ridgway, and J. D. McPhail. 1984. Ecology and evolution of sympatric sticklebacks (*Gasterosteus*): spacial segregation and seasonal habitat shifts in the Enos Lake species pair. Canadian Journal of Zoology 62:2436–2439.

Bergstrom, C. A. 2002. Fast-start swimming performance and reduction in lateral plate number in threespine stickleback. Canadian Journal of Zoology 80:207–213.

Berner, D., D. C. Adams, A.-C. Grandchamp, and A. P. Hendry. 2008. Natural selection drives patterns of lake–stream divergence in stickleback foraging morphology. J. Evol. Biol. 21:1653–1665.

Berner, D., A.-C. Grandchamp, and A. P. Hendry. 2009. Variable Progress toward Ecological Speciation in Parapatry: Stickleback across Eight Lake-Stream Transitions. Evolution 63:1740–1753.

Bjærke, O., K. Østbye, H. M. Lampe, and L. A. Vøllestad. 2010. Covariation in shape and foraging behaviour in lateral plate morphs in the three-spined stickleback. Ecol. Freshwat. Fish 19:249–256.

Brooks, J. L. 1957. The Systematics of North American *Daphnia*. The Connecticut Academy of Arts and Sciences.

Brooks, J. L. and S. I. Dodson. 1965. The effect of a marine planktivore on lake plankton illustrates theory of size, competition, and predation. Science 150:28–35.

Cangelosi, R. and A. Goriely. 2007. Component retention in principal component analysis with application to cDNA microarray data. Biology Direct 2:2.

Chaverie, L., W. J. Harford, K. L. Howland, J. Fitzsimons, A. M. Muir, C. C. Krueger, and W. M. Tonn. 2016. Multiple generalist morphs of Lake Trout: Avoiding constraints on the evolution of intraspecific divergence? Ecology and Evolution 6:7727–7741.

Chaverie, L., K. L. Howland, L. N. Harris, C. P. Gallagher, M. J. Hansen, W. M. Tonn, A. M. Muir, and C. C. Krueger. 2020. Among-individual diet variation within a lake trout ecotype: lack of stability of niche use. Ecology and Evolution 11: 1457–1475.

Cheverud, J. M. 1982. Phenotypic, genetic, and environmental integration in the cranium. Evolution 36:499–516.

Cheverud, J. M. 2001. A simple correction for multiple comparisons in interval mapping genome scans. Heredity 87:52–58.

Corti, M., C. Fadda, S. Simson, and E. Nevo. 1996. Size and shape variation in the mandible of the fossorial rodent *Spalax ehrenbergi:* a Procrustes analysis of three dimensions. Pp. 303–320 *in* L. F. Marcus, M. Corti, A. Loy, G. J. P. Naylor, and D. E. Slice, eds. Advances in Morphometrics. Plenum Press, New York.

Del Giudice, M. 2020. Effective dimensionality: a tutorial. Multivariate Behavioral Research.

EPA. 1994. Method 200.7, Revision 4.4: Determination of metals and trace elements in water and wastes by inductively coupled plasma-atomic emission spectrometry *in* E. M. S. L. O. o. R. a. Development, ed. U.S. Environmental Protection Agency, Cincinnati, Ohio 45268.

Fadda, C. and M. Corti. 1998. Geographic variation of *Arvicanthis* (Rodentia, Muridae) in the Nile Valley. Z. Säugetierkunde 63:104–113.

Feilich, K. L. 2016. Correlated evolution of body and fin morphology in the cichlid fishes. Evolution 70:2247–2267.

Felice, R. N., J. A. Tobias, A. L. Pigot, and A. Goswami. 2019. Dietary niche and the evolution of cranial morphology in birds. Proc. R. Soc. B. 286:2018–2577.

Gelmond, O., F. A. v. Hippel, and M. S. Christy. 2009. Rapid ecological speciation in three-spined stickleback *Gasterosteus aculeatus* from Middleton Island, Alaska: the roles of selection and geographic isolation. J. Fish Biol. 75:2037–2051.

Giles, N. 1983. The possible role of environmental calcium levels during the evolution of phenotypic diversity in Outer Hebriddean populations of the three-spined stickleback, *Gasterosteus aculeatus*. J. Zool. 199:535–544.

Glazer, A. M., P. A. Cleves, P. A. Erickson, A. Y. Lam, and C. T. Miller. 2014. Parallel developmental genetic features underlie stickleback gill raker evolution. EvoDevo 5:19.

Griesbach, S. J. and R. H. Peters. 1991. The effects of analytical variations on estimates of phosphorus concentration in surface waters. Lake Reserv Manag 7:97–106.

Hagen, D. W. 1967. Isolating mechanisms in threespine sticklebacks (*Gasterosteus*). J. Fish. Res. Bd. Canada 24:1637–1692.

Haney, J. F., M. A. Aliberti, E. Allan, S. Allard, D. J. Bauer, W. Beagen, S. R. Bradt, B. Carlson, S. C. Carlson, U. M. Doan, J. Dufresne, W. T. Godkin, S. Greene, A. Kaplan, E. Maroni, S. Melillo, A. L. Murby, J. L. S. (Nowak), B. Ortman, J. E. Quist, S. Reed, T. Rowin, M. Schmuck, R. S. Stemberger, and B. Travers. 2013. An image-based key to the zooplankton of North America, version 5.0. University of New Hampshire Center for Freshwater Biology, cfb.unh.edu/cfbkey/html/.

Hansen, T. F. and D. Houle. 2008. Measuring and comparing evolvability and constraint in multivariate characters. J. Evol. Biol. 21:1201–1219.

Hendry, A. P. and E. B. Taylor. 2004. How much of the variation in adaptive divergence can be explained by gene flow? An evaluation using lake-stream stickleback pairs. Evolution 58:2319–2331.

Hine, E., K. McGuigan, and M. W. Blows. 2014. Evolutionary constraints in high-dimensional trait sets. Am. Nat. 184:119–131.

Hulsey, C. D., M. E. Alfaro, J. Zheng, A. Meyer, and R. Holzman. 2019. Pleiotropic jaw morphology links the evolution of mechanical modularity and functional feeding convergence in Lake Malawi cichlids. Proceedings of the Royal Society B: Biological Sciences 286:20182358.

Hyman, J. M. 1983. Accurate monotonicity preserving cubic interpolation. SIAM J. Sci. and Stat. Comput. 4:645–654.

Izen, R., Y. E. Stuart, Y. Jiang, and D. I. Bolnick. 2016. Coarse- and fine-grained phenotypic divergence among threespine stickleback from alternating lake and stream habitats. Evol. Ecol. Res. 17:437–457.

Jones, C., K. M. Somers, B. Craig, and T. B. Reynoldson. 2007. Ontario benthos biomonitoring network: protocol manual. Queen’s Printer for Ontario.

Jones, J. R., M. A. Bell, J. A. Baker, and J. P. Koenings. 2003. General limnology of lakes near Cook Inlet, Southcentral Alaska. Lake Reserv. Manage. 19:141–149.

Jónsson, B. 2000. Polymorphic segregation in Arctic charr *Salvelinus alpinus* (L.) from Vatnshlídarvatn, a shallow Icelandic lake. Biol. J. Linn. Soc. 69:55–74.

Kirkpatrick, M. 2009. Patterns of quantitative genetic variation in multiple dimensions. Genetica 136:271–284.

Klepaker, T., K. Østbye, and M. A. Bell. 2013. Regressive evolution of the pelvic complex in stickleback fishes: a study of convergent evolution. Evol. Ecol. Res. 15:413–435.

Klepaker, T., K. Østbye, R. Spence, M. Warren, M. Przybylski, and C. Smith. 2016. Selective agents in the adaptive radiation of Hebridean sticklebacks. Evol. Ecol. Res. 17:243–262.

Langerhans, R. B. 2018. Predictability and parallelism of multitrait adaptation. J. Hered. 109:59–70.

Lavin, P. A. and J. D. McPhail. 1993. Parapatric lake and stream sticklebacks on northern Vancouver Island: disjunct distribution or parallel evolution? Canadian Journal of Zoology 71:11–18.

Legendre, P., J. Oksanen, and C. J. F. t. Braak. 2011. Testing the significance of canonical axes in redundancy analysis. Methods in Ecology and Evolution 2:269–277.

Levin, B., E. Simonov, P. Franchini, N. Mugue, A. Golubtsov, and A. Meyer. 2021. Adaptive radiation and burst speciation of hillsteam cyprinid fish *Garra* in African river. BioRxiv.

Lipka, C. G., J. L. Gates, and S. K. Simons. 2020. Sport fisheries of the Northern Kenai Peninsula Management Area, 2016-2018, with overview for 2019. Fishery Management Report. Alaska Department of Fish and Game, Anchorage, Alaska.

MacColl, A. D. C., A. E. Nagar, and J. d. Roij. 2013. The evolutionary ecology of dwarfism in three-spined sticklebacks. J. Anim. Ecol. 82:642–652.

Mack, H. R., J. D. Conroy, K. A. Blocksom, R. A. Stein, and S. A. Ludsin. 2012. A comparative analysis of zooplankton field collection and sample enumeration methods. Limnol. Oceanogr. Methods 10:41–53.

Marques, D. A., F. C. Jones, F. D. Palma, D. M. Kingsley, and T. E. Reimchen. 2018. Experimental evidence for rapid genomic adaptation to a new niche in an adaptive radiation. Nature Ecology & Evolution 2:1128–1138.

Massengill, R., R. N. Begich, and K. Dunker. 2020. Operational Plan: Kenai Peninsula nonnative fish control, monitoring, and native fish restoration. Regional Operation Plan. Alaska Department of Fish and Game, Anchorage, Alaska.

McGee, M. D., D. Schluter, and P. C. Wainwright. 2013. Functional basis of ecological divergence in sympatric stickleback. BMC Evol. Biol. 13:277.

McPhail, J. D. 1993. Ecology and evolution of sympatric sticklebacks *(Gasterosteus):* origin of the species pairs. Can J Zool 71:515–523.

Merritt, R. W., K. W. Cummins, and M. B. Berg, eds. 2008. Introduction to the aquatic insects of North America. Kendall Hunt Publishing Company, Dubuque, IA, USA.

Mezey, J. G. and D. Houle. 2005. The dimensionality of genetic variation for wing shape in *Drosophila melanogaster*. Evolution 59:1027–1038.

Mitteroecker, P. and F. Bookstein. 2007. The conceptual and statistical relationship between modularity and morphological integration. Syst. Biol. 56:818–836.

Moisan, J. 2010. Guide d’identification des principaux macroinvertébrés benthiques d’eau douce du Québec, 2010 – Surveillance volontaire des cours d’eau peu profonds. Bibliothèque et archives nationales du Québec (CA): Direction du suivi de l’état de l’environnement, ministère du Développement durable, de l’Environnement et des Parcs..

Muir, A. M., M. J. Hansen, C. R. Bronte, and C. C. Krueger. 2016. If Arctic charr *Salvelinus alpinus* is ‘the most diverse vertebrate’, what is the lake charr *Salvelinus namaycush?* Fish Fish. 17:1194–1207.

Noble, D. W. A., R. Radersma, and T. Uller. 2019. Plastic responses to novel environments are biased toward phenotype dimensions with high additive genetic variation. Proceedings of the National Academy of Sciences 116:13452–13461.

Nosil, P., L. J. Harmon, and O. Seehausen. 2009. Ecological explanations for (incomplete) speciation. Trends Ecol. Evol. 24:145–156.

Oke, K. B., G. Rolshausen, C. LeBlond, and A. P. Hendry. 2017. How parallel is parallel evolution? a comparative analysis in fishes. Am. Nat. 190:1–16.

Palkovacs, E. P. and D. M. Post. 2008. Eco-evolutionary interactions between predators and prey: can predator-induced changes to prey communities feed back to shape predator foraging traits? Evol. Ecol. Res. 10:699–720.

Palkovacs, E. P. and D. M. Post. 2009. Experimental evidence that phenotypic divergence in predators drives community divergence in prey. Ecology 90:300–305.

Patton, C. J. and J. R. Kryskalla. 2003. Methods of analysis by the U.S. Geological Survey National Water Quality Laboratory: evaluation of alkaline persulfate digestion as an alternative to Kjeldahl digestion for determination of total and dissolved nitrogen and phosphorus in water. Water-Resources Investigations Report. US Department of the Interior, US Geological Survey.

Pirkl, R. J., K. A. Remley, and C. S. L. Patane. 2012. Reverberation chamber measurement correlation. IEEE Transactions on Electromagnetic Compatibility 54:533–545.

R Core Team. 2019. R: A language and environment for statistical computing. R Foundation for Statistical Computing, Vienna, Austria.

Rasband, W. S. 1997-2018. ImageJ. U.S. National Institutes of Health, Bethesda, Maryland, USA.

Reimchen, T. E. 1991. Trout foraging failures and the evolution of body size in stickleback. Copeia 1991:1098–1104.

Reimchen, T. E. 1994. Predators and morphological evolution in threespine stickleback. Pp. 240–276 *in* M. A. Bell, and S. A. Foster, eds. The evolutionary biology of the threespine stickleback. Oxford University Press, New York.

Reimchen, T. E., C. Bergstrom, and P. Nosil. 2013. Natural selection and the adaptive radiation of Haida Gwaii stickleback. Evol. Ecol. Res. 15:241–269.

Reisch, R., T. Easter, C. A. Layman, and R. B. Langerhans. 2015. Rapid human-induced divergence of life-history strategies in Bahamian livebearing fishes (family Poeciliidae). J. Anim. Ecol. 84:1732–1743.

Reist, J. D. 1986. An empirical evaluation of coefficients used in residual and allometric adjustment of size covariation. Can. J. Zool. 64:1363–1368.

Ripley, B., B. Venables, D. M. Bates, K. Hornik, A. Gebhardt, and D. Firth. 2020. Support Functions and Datasets for Venables and Ripley’s MASS. Pp. Functions and datasets to support Venables and Ripley, “Modern Applied Statistics with S” (4th edition, 2002). *in* B. Ripley, ed, CRAN.

Rohlf, F. J. 2018. tpsDIG2, digitize landmarks and outlines. SUNY Stony Brook.

Rohlf, F. J. and M. Corti. 2000. Use of two-block partial least-squares to study covariation in shape. Syst. Biol. 49:740–753.

Roy, O. and M. Vetterli. 2007. The effective rank: A measure of effective dimensionality. Pp. 606–610 *in* M. Domański, and R. Stasiński, eds. 15th European Signal Processing Conference.

Runquist, R. D. B., A. J. Gorton, J. B. Yoder, N. J. Deacon, J. J. Grossman, S. Kothari, M. P. Lyons, S. N. Sheth, P. Tiffin, and D. A. Moeller. 2020. Context dependence of local adaptation to abiotic and biotic environments: A quantitative and qualitative synthesis. The American Naturalist 195:412–431.

Sartory, D. P. and J. U. Grobbelaar. 1984. Extraction of chlorophyll *a* from freshwater phytoplankton for spectrophotometric analysis. Hydrobiologia 114:177–187.

Schlager, S. 2017. Morpho and Rvcg – Shape analysis in R: packages for geometric morphometrics, shape analysis and surface manipulations. Pp. 217–256 *in* G. Zheng, S. Li, and G. Szekely, eds. Statistical shape and deformation analysis. Academic Press, London, UK.

Schlager, S., G. Jefferis, and D. Ian. 2020. Morpho: Calculations and Visualisations Related to Geometric Morphometrics *in* S. Schlager, ed, CRAN.

Schluter, D. 1996. Adaptive radiation along genetic lines of least resistance. Evolution 50:1766–1774.

Schluter, D. and J. D. McPhail. 1992. Ecological character displacement and speciation in sticklebacks. Am. Nat. 140:85–108.

Schmid, D. W., M. D. McGee, R. J. Best, O. Seehausen, and B. Matthews. 2019. Rapid divergence of predator functional traits affects prey composition in aquatic communities. Am. Nat. 193:331–345.

Seebacher, F., M. M. Webster, R. S. James, J. Tallis, and A. J. W. Ward. 2016. Morphological differences between habitats are associated with physiological and behavioural trade-offs in stickleback *(Gasterosteus aculeatus)*. Royal Society Open Science 3:160316.

Skúlason, S., S. S. Snorrason, D. L. G. Noakes, M. M. Ferguson, and H. J. Malmquist. 1989. Segregation in spawning and early life history among polymorphic Arctic charr, *Salvelinus alpinus,* in Thingvallavatn, Iceland. J. Fish Biol. 35:225–232.

Spoljaric, M. A. and T. E. Reimchen. 2007. 10 000 years later: evolution of body shape in Haida Gwaii three-spined stickleback. J. Fish Biol. 70:1484–1503.

Springer, V. G. and G. D. Johnson. 2000. Use and advantages of ethanol solution of alizarin red S dye for staining bone in fishes. Copeia 2000:300–301.

Stuart, Y. E., T. Veen, J. N. Weber, D. Hanson, M. Ravinet, B. K. Lohman, C. J. Thompson, T. Tasneem, A. Doggett, R. Izen, N. Ahmed, R. D. H. Barrett, A. P. Hendry, C. L. Peichel, and D. I. Bolnick. 2017. Contrasting effects of environment and genetics generate a continuum of parallel evolution. Nature Ecology & Evolution 1:0158.

Taylor, D. J. and P. D. N. Hebert. 1993. Cryptic intercontinental hybridization in *Daphnia* (Crustacea): The ghost of introductions past. Proceedings of the Royal Society B: Biological Sciences 254:163–168.

Taylor, E. B. and J. D. McPhail. 1986. Prolonged and burst swimming in anadromous and freshwater threespine stickleback, *Gasterosteus aculeatus*. Canadian Journal of Zoology 64:416–420.

Thorp, J. H. and A. P. Covich, eds. 2010. Ecology and Classification of North American Freshwater Invertebrates. Academic Press, Burlington, MA, USA.

Velasco, J. A. and A. Herrel. 2007. Ecomorphology of *Anolis* lizards of the Choco’ region in Colombia and comparisons with Greater Antillean ecomorphs. Biol. J. Linn. Soc. 92:29–39.

Venables, W. N. and B. D. Ripley. 2002. Classification *in* W. N. Venables, and B. D. Ripley, eds. Modern applied statistics with S. Springer, New York.

Wagner, G. P. and L. Altenberg. 1996. Perpective: Complex adaptations and the evolution of evolvability. Evolution 50:967–976.

Wagner, G. P., M. Pavlicev, and J. M. Cheverud. 2007. The road to modularity. Nat. Rev. Genet. 8:921–931.

Wainwright, P. C., M. E. Alfaro, D. I. Bolnick, and C. D. Hulsey. 2005. Many-to-one mapping of form to function: A general principle in organismal design? Integr. Comp. Biol. 45:256–262.

Walker, J. A. 1997. Ecological morphology of lacustrine threespine stickleback *Gasterosteus aculeatus* L. (Gasterosteidae) body shape. Biol. J. Linn. Soc. 61:3–50.

Watanabe, J. 2021. Detecting (non)parallel evolution in multidimensional space: angles, correlations, and eigenanalysis, EcoEvoRxiv.

Welch, J. J. and D. Waxman. 2003. Modularity and the cost of complexity. Evolution 57:1723–1734.

Willacker, J. J., F. A. von Hippel, P. R. Wilton, and K. M. Walton. 2010. Classification of the threespine stickleback along the benthic-limnetic axis. Biol. J. Linn. Soc. 101:595–608.

Wintermans, J. F. and A. de Mots. 1965. Spectrophotometric characteristics of chlorophylls a and b and their pheophytins in ethanol. Biochim Biophys Acta. 109:448–453.

Witty, L. M. 2004. Practical guide to identifying freshwater crustacean zooplankton, 2nd edition *in* L. U. Cooperative Freshwater Ecology Unit – Department of Biology, ed.

Yong, L., D. P. Croft, J. Troscianko, I. Ramnarine, and A. Wilson. 2020. sensory-based quantification of male colour patterns in Trinidadian guppies reveals nonparallel phenotypic evolution across an ecological transition in multivariate trait space. BioRxiv.

Zelditch, M. L., D. L. Swiderski, and H. D. Sheets. 2012. General linear models. Pp. 225-260. Geometric morphometrics for biologists: a primer. Academic Press, Waltham, Mass., USA.

