## SupplementalFigures for "Dimensionality and modularity of adaptive variation: Divergence in threespine stickleback from diverse environments"

### Supplementary Figures

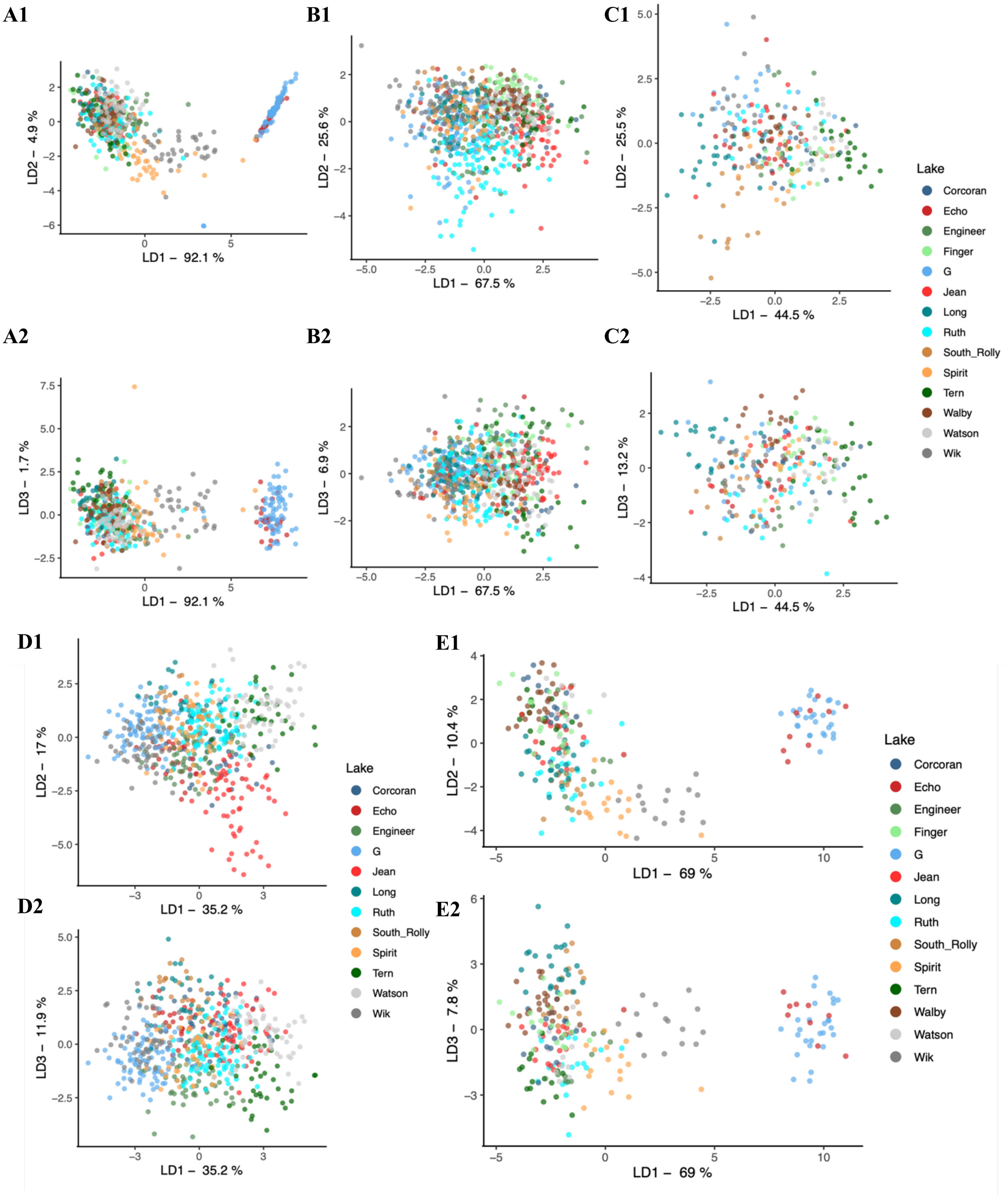


Figure S1. Scatter plots showing the positions of individuals in spaces defined by discriminant scores for Defensive traits (A1 & A2), Swimming traits (B1 & B2), Trophic traits (C1 & C2), and Shape (D1 & D2). Panels E1 and E2 show discriminant axes of all traits excluding shape. For each trait suite, the first panel shows LD1 and LD2, and the second panel shows LD1 and LD3. Panels are sized differently to maintain 1:1 scaling of units on the X and Y axes.

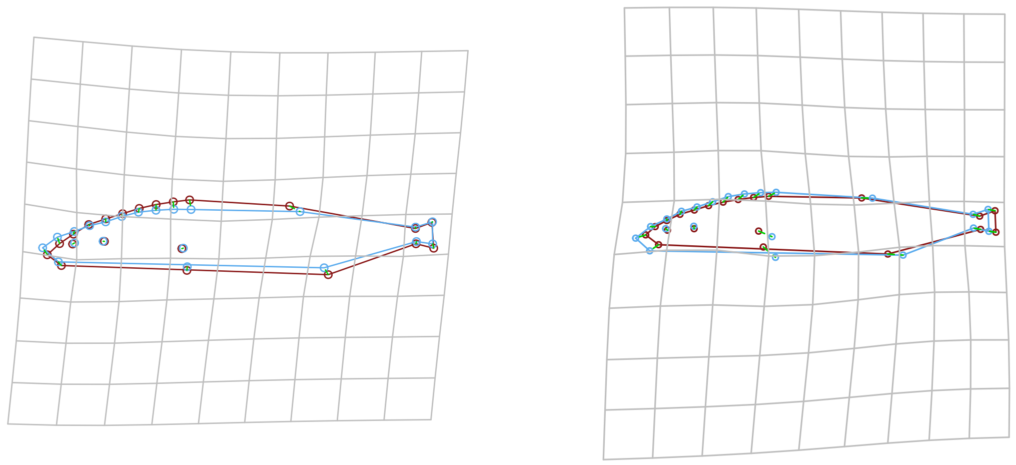

CV1 ± 5 CV2 ± 5

Figure S2. Differences between stickleback shapes along CVA axes 1 and 2. Shapes shown in red represent shape at CV values of +5, and shapes shown in blue represent CV values of -5.

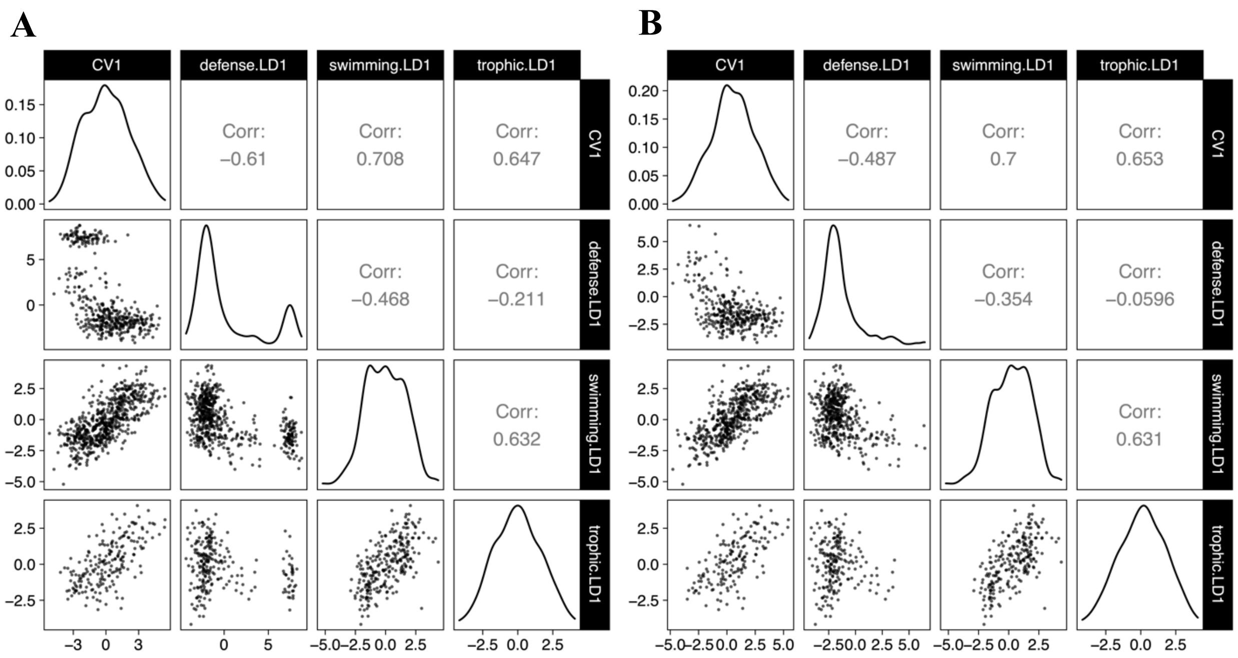


Figure S3. Correlation matrices of major linear discriminant axes of trait suites. Panel A shows all lakes, while panel B excludes individuals from Echo and G lakes because their extreme Defense LD1 values resulted in a bimodal distribution (and thus severe departure from a normal distribution). Correlation coefficients shown are Pearson's *r*.

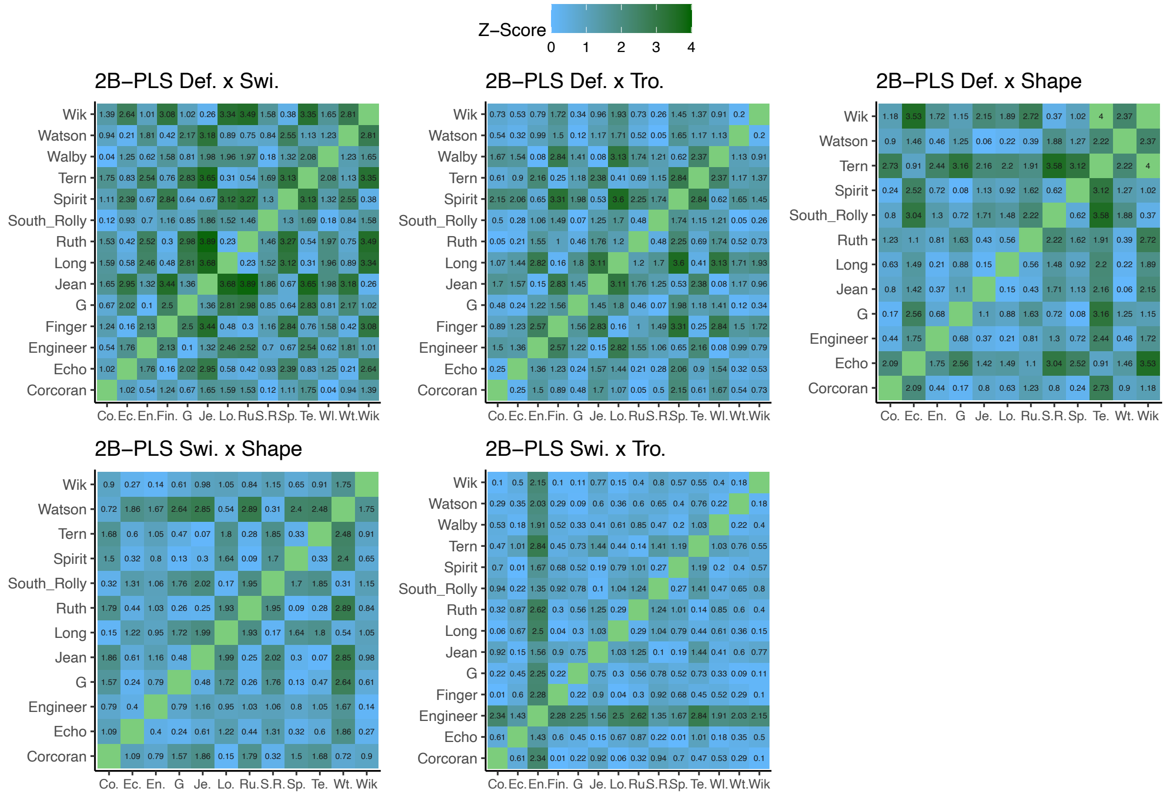


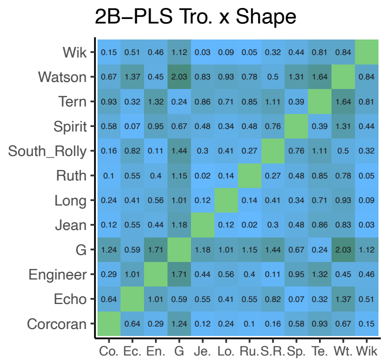


Figure S4. Effect sizes of pairwise two-sample Z-tests of integration strength between lakes, showing differences in strength of integration between lake pairs. Each matrix represents a combination of trait suites whose integration was assessed using 2-block PLS correlation. Effect sizes being compared are effect sizes representing strength of integration between the two blocks in each lake.

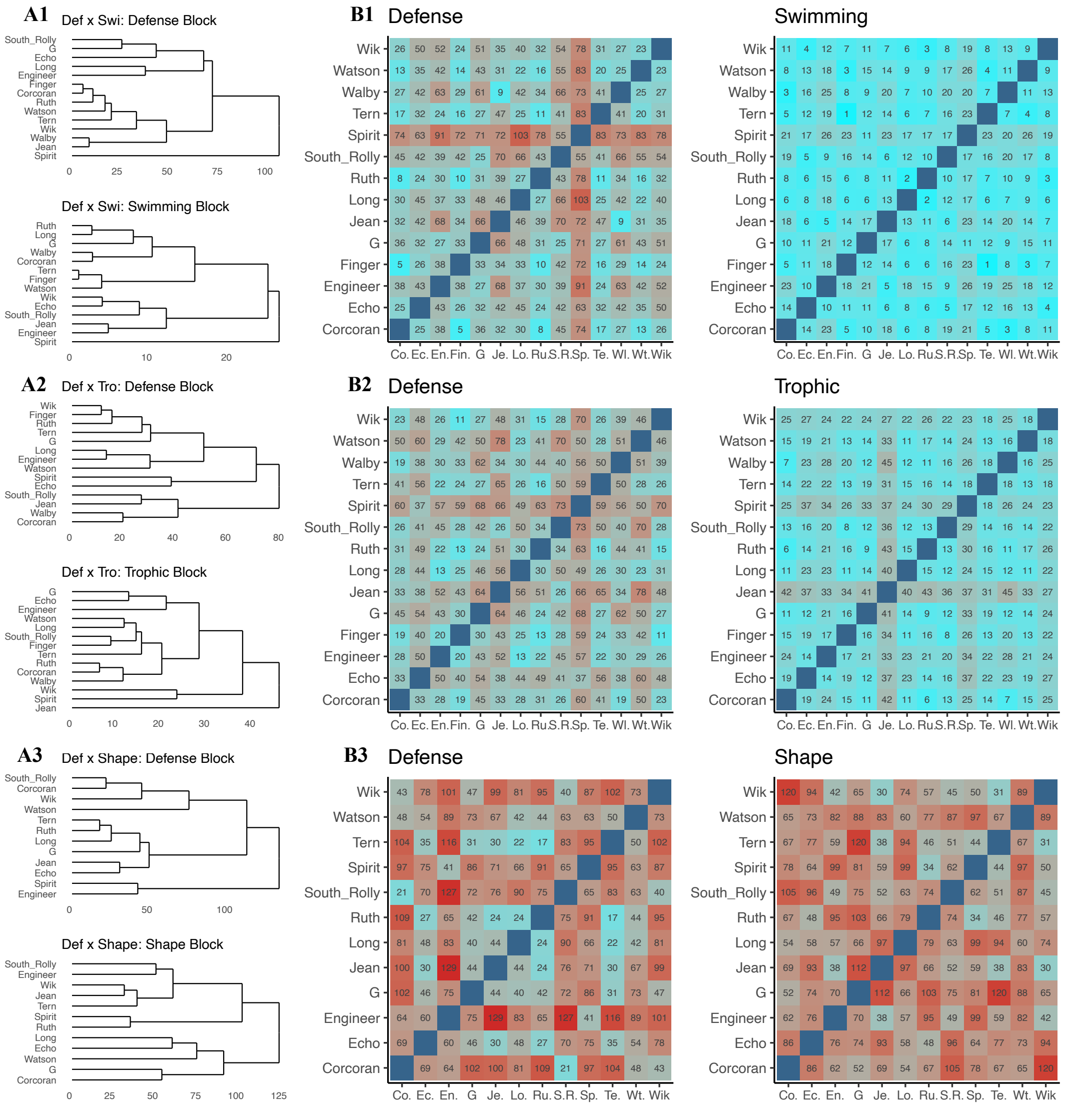


Figure S5. Angles of two-block PLS vectors, in pairwise comparisons between lakes. Dendrograms in panels A1-A6 show hierarchical clustering of angle matrices shown in panels B1-B6, using angle as the distance metric. Matrices in panels B1-B6 correspond to major vectors through left and right data blocks for trait suites in 2B-PLS tests of integration, and numbers in cells indicate angle in degrees.

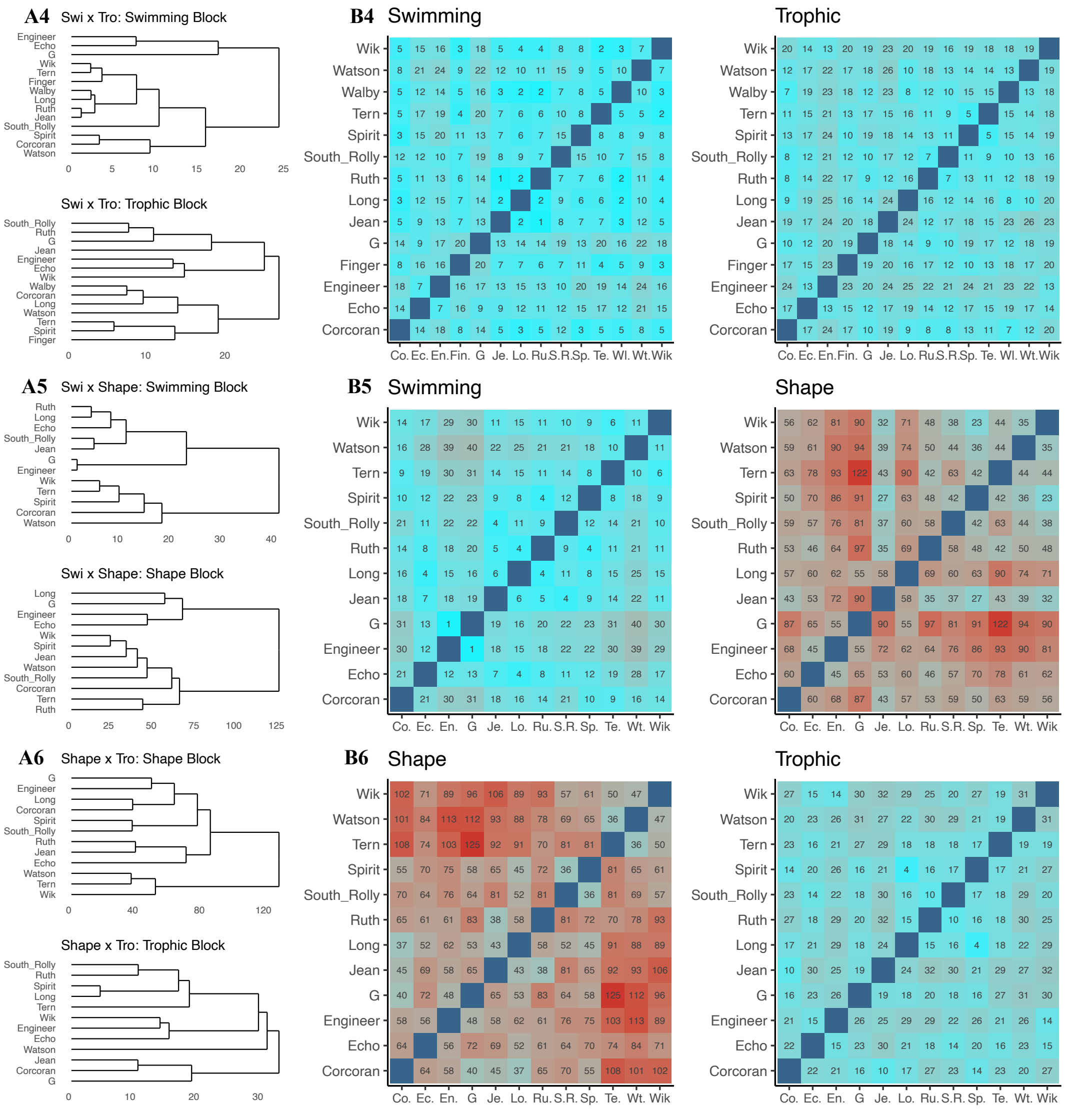


Figure S5 cont'd.
