## SupplementalTables for "Dimensionality and modularity of adaptive variation: Divergence in threespine stickleback from diverse environments"

### Supplementary Tables

Table S1. Phenotypic traits used for analysis, the trait suites to which they were assigned, the abbreviations used for them in the text, and a brief description of each trait. Traits indicated with an * were excluded from all multidimensional analyses.

| Assigned Trait Suite / Group | Trait | Abbreviation | Description | Support for Trait Suite Assignment |
|---|---|---|---|---|
| **Defense** | Lateral Plate Count | LP | The number of lateral plates, if the number differed on R and L sides, mean number was used | Reimchen (2000), Bell et al. (2004), Stuart et al. (2017) |
| | Mean Lateral Plate Position | $LP_M$ | Mean plate position, calculated from position of first and last plates on both sides | Reimchen (1983) |
| | Pelvic Spine Length | $PS_L$ | Mean length of both pelvic spines from base to tip of spine. If one was broken, only intact spine was used | Reimchen (2000), Stuart et al. (2017), Marques et al. (2018) |
| | First Dorsal Spine Length | $DS1_L$ | From base to tip of spine | Stuart et al. (2017), Marques et al. (2018) |
| | Second Dorsal Spine Length | $DS2_L$ | From base to tip of spine | Stuart et al. (2017), Marques et al. (2018) |
| **Swimming** | Standard Length | SL | Length from tip of snout to base of posterior-most caudal fin ray | Walker (1997), Stuart et al. (2017) |
| | Body Depth | BD | Measured from base of DS1 to anterior of pelvis | Walker (1997), Stuart et al. (2017) |
| | Caudal Peduncle Width | CP | Top to bottom of caudal peduncle at narrowest point | Walker (1997), Stuart et al. (2017) |
| **Trophic** | Jaw Length | JL | Tip of snout to posterior tip of maxilla | Reimchen and Nosil (2006), Marques et al. (2018) |
| | Gape Width | GW | Maximum width between tips of maxillae | McGee and Wainwright (2013), Stuart et al. (2017) |
| | Pterotic Width | PW | Measured at widest portion of posterior process | Anker (1974) |
| | Buccal Cavity Length | BC | Measured from tip of snout to anterior extent of ectocoracoid | Caldecutt and Adams (1998), Stuart et al. (2017) |
| | Gill Raker Count | GR | Count of gill rakers on distal row of 1st right branchial arch | Hendry and Taylor (2004), Berner et al. (2009), Stuart et al. (2017) |
| | Snout Length | SN | Measured from tip of snout to anterior edge of eye | Reimchen et al. (2016), Caldecutt and Adams (1998) |
| | Head Length | HL | Measured from tip of snout to the posterior margin of operculum | Reimchen et al. (2016), Caldecutt and Adams (1998) |
| | Eye Diameter | ED | Measured from anterior to posterior edge of eye | Reimchen and Nosil (2006), Reimchen et al. (2016) |
| **Shape** | Landmark 1 | | Tip of snout | |
| | Landmark 2 | | Posterior tip of maxilla | |
| | Landmark 3 | | Anterior edge of eye | |
| | Landmark 4 | | Posterior edge of eye | |
| | Landmark 5 | | Dorsal insertion of pectoral fin | |
| | Landmark 6 | | Ventral insertion of pectoral fin | |
| | Landmark 7 | | Anterior insertion of anal fin | |
| | Landmark 8 | | Ventral edge at narrowest point of caudal peduncle | |
| | Landmark 9 | | Insertion of ventral-most caudal fin ray | |
| | Landmark 10 | | Insertion of dorsal-most caudal fin ray | |
| | Landmark 11 | | Dorsal edge at narrowest point of caudal peduncle | |
| | Landmark 12 | | Anterior insertion of dorsal fin | |
| | Landmark 13 | | Base of first dorsal spine | |

| | | | |
|---|---|---|---|
| | 8 Semi-landmarks | | Evenly spaced between Landmarks 1 and 13 |
| **Environment** | Surface Area | Area | hectares |
| | Maximum Depth | MaxDepth | meters |
| | Chlorophyll-a | Chl-a | $\mu gL^{-1}$ |
| | Total Nitrogen | TN | $mgL^{-1}$ |
| | Total Phosphorus | TP | $\mu gL^{-1}$ |
| | Dissolved Calcium | Ca | $mgL^{-1}$ |
| | Conductivity | Cond | $\mu S.cm^{-2}$ |
| | Dissolved Organic Carbon | DOC | $mgL^{-1}$ |
| | pH | pH | |
| | % Littoral Area | Litt. | |
| | *Daphnia* Abundance | *Daph.* | individuals $L^{-1}$ |
| | Gammarid Abundance | Gamm. | individuals $(m^2)^{-1}$ |


Anker, G. C. 1974. Morphology and kinetics of the head of the stickleback, *Gasterosteus aculeatus*. Transactions of the Zoological Society of London 32:311-416.

Bell, M. A., W. E. Aguirre, and N. J. Buck. 2004. Twelve years of contemporary armor evolution in a threespine stickleback population. Evolution 58:814-824.

Berner, D., A.-C. Grandchamp, and A. P. Hendry. 2009. Variable Progress toward Ecological Speciation in Parapatry: Stickleback across Eight Lake-Stream Transitions. Evolution 63:1740-1753.

Caldecutt, W. J. and D. C. Adams. 1998. Morphometrics of trophic osteology in the threespine stickleback, *Gasterosteus aculeatus*. Copeia 4:827-838.

Hendry, A. P. and E. B. Taylor. 2004. How much of the variation in adaptive divergence can be explained by gene flow? An evaluation using lake-stream stickleback pairs. Evolution 58:2319-2331.

Marques, D. A., F. C. Jones, F. D. Palma, D. M. Kingsley, and T. E. Reimchen. 2018. Experimental evidence for rapid genomic adaptation to a new niche in an adaptive radiation. Nature Ecology & Evolution 2:1128-1138.

McGee, M. D. and P. C. Wainwright. 2013. Convergent evolution as a generator of phenotypic diversity in threespine stickleback. Evolution 64:1204-1208.

Reimchen, T. E. 1983. Structural relationships between spines and lateral plates in threespine stickleback (*Gasterosteus aculeatus*). Evolution 37:931-946.

Reimchen, T. E. 2000. Predator handling failures of lateral plate morphs in gasterosteus aculeatus: Functional implications for the ancestral plate condition. Behaviour 137:1081-1096.

Reimchen, T. E. and P. Nosil. 2006. Replicated ecological landscapes and the evolution of morphological diversity among *Gasterosteus* populations from an archipelago on the west coast of Canada. Canadian Journal of Zoology 84:643-654.

Reimchen, T. E., D. Steeves, and C. A. Bergstrom. 2016. Sex matters for defense and trophic traits of threespine stickleback. Evol. Ecol. Res. 17:459-485.

Stuart, Y. E., T. Veen, J. N. Weber, D. Hanson, M. Ravinet, B. K. Lohman, C. J. Thompson, T. Tasneem, A. Doggett, R. Izen, N. Ahmed, R. D. H. Barrett, A. P. Hendry, C. L. Peichel, and D. I. Bolnick. 2017. Contrasting effects of environment and genetics generate a continuum of parallel evolution. Nature Ecology & Evolution 1:0158.

Walker, J. A. 1997. Ecological morphology of lacustrine threespine stickleback *Gasterosteus aculeatus* L. (Gasterosteidae) body shape. Biol. J. Linn. Soc. 61:3-50.


Table S2. Sample sizes for each lake and trait before interpolation of missing values ($n_I$) and the number of individuals for which values were interpolated ($n_+$). Values for some traits, particularly meristic traits, were not interpolated: LP, $LP_M$, and GR. Shape and LDA columns show sample sizes for each lake.

| Lake | Defense | | | | | | | | | | Swimming | | | | | |
|---|---|---|---|---|---|---|---|---|---|---|---|---|---|---|---|---|
| | LP | | $DS1_L$ | | $DS2_L$ | | $PS_L$ | | $LP_M$ | | SL | | BD | | CP | |
| | $n_I$ | $n_+$ | $n_I$ | $n_+$ | $n_I$ | $n_+$ | $n_I$ | $n_+$ | $n_I$ | $n_+$ | $n_I$ | $n_+$ | $n_I$ | $n_+$ | $n_I$ | $n_+$ |
| *Corcoran* | 21 | - | 21 | - | 21 | - | 21 | - | 21 | - | 41 | - | 41 | - | 41 | - |
| *Echo* | 28 | - | 24 | 3 | 27 | - | 28 | - | 28 | - | 28 | - | 28 | - | 27 | 1 |
| *Engineer* | 41 | - | 35 | 6 | 41 | - | 40 | 1 | 41 | - | 60 | - | 60 | - | 60 | - |
| *Finger* | 42 | - | 38 | 4 | 42 | - | 42 | - | 42 | - | 60 | - | 60 | - | 59 | 1 |
| *G* | 67 | - | 66 | 1 | 67 | - | 67 | - | 67 | - | 100 | - | 100 | - | 100 | - |
| *Jean* | 37 | - | 36 | 1 | 37 | - | 37 | - | 37 | - | 60 | - | 60 | - | 60 | - |
| *Long* | 32 | - | 31 | 1 | 31 | 1 | 32 | - | 32 | - | 64 | - | 64 | - | 63 | 1 |
| *Ruth* | 61 | - | 58 | 3 | 59 | 2 | 61 | - | 61 | - | 99 | - | 99 | - | 99 | - |
| *South_Rolly* | 23 | - | 22 | 1 | 23 | - | 23 | - | 23 | - | 23 | - | 23 | - | 23 | - |
| *Spirit* | 37 | - | 33 | 4 | 36 | 1 | 36 | 1 | 37 | - | 50 | - | 50 | - | 50 | - |
| *Tern* | 25 | - | 23 | 2 | 25 | - | 25 | - | 25 | - | 37 | - | 37 | - | 37 | - |
| *Walby* | 44 | - | 43 | 1 | 44 | - | 44 | - | 44 | - | 60 | - | 60 | - | 59 | 1 |
| *Watson* | 56 | - | 53 | 3 | 55 | 1 | 56 | - | 56 | - | 60 | - | 60 | - | 60 | - |
| *Wik* | 38 | - | 34 | 4 | 38 | - | 38 | - | 38 | - | 50 | - | 50 | - | 50 | - |

| Lake | Trophic | | | | | | | | | | | | | | | | | | Shape | LDAs | | | | |
|---|---|---|---|---|---|---|---|---|---|---|---|---|---|---|---|---|---|---|---|---|---|---|---|---|
| | BC | | GW | | PW | | GR | | JL | | SN | | ED | | HL | | | | | Def | Swi | Tro | | Shape |
| | $n_I$ | $n_+$ | $n_I$ | $n_+$ | $n_I$ | $n_+$ | $n_I$ | $n_+$ | $n_I$ | $n_+$ | $n_I$ | $n_+$ | $n_I$ | $n_+$ | $n_I$ | $n_+$ | | | | | | w/GR | no GR | |
| *Corcoran* | 41 | - | 19 | 22 | 19 | 22 | 19 | - | 41 | - | 41 | - | 41 | - | 41 | - | | | 41 | 21 | 41 | 19 | 41 | 41 |
| *Echo* | 28 | - | 11 | 17 | 11 | 17 | 11 | - | 27 | - | 27 | - | 27 | - | 27 | - | | | 27 | 27 | 28 | 10 | 27 | 27 |
| *Engineer* | 60 | - | 15 | 45 | 15 | 45 | 10 | - | 60 | - | 60 | - | 60 | - | 60 | - | | | 55 | 41 | 60 | 10 | 60 | 55 |
| *Finger* | 60 | - | 20 | 40 | 20 | 40 | 20 | - | 59 | 1 | 59 | 1 | 59 | 1 | 59 | 1 | | | 0 | 42 | 60 | 20 | 60 | - |
| *G* | 100 | - | 32 | 68 | 32 | 68 | 27 | - | 100 | - | 100 | - | 100 | - | 100 | - | | | 92 | 67 | 100 | 27 | 100 | 92 |
| *Jean* | 60 | - | 20 | 40 | 20 | 40 | 20 | - | 60 | - | 60 | - | 60 | - | 60 | - | | | 59 | 37 | 60 | 20 | 60 | 59 |
| *Long* | 64 | - | 25 | 38 | 25 | 39 | 25 | - | 62 | 1 | 62 | 1 | 63 | - | 62 | 1 | | | 33 | 32 | 64 | 25 | 63 | 33 |
| *Ruth* | 99 | - | 20 | 79 | 20 | 79 | 20 | - | 99 | - | 99 | - | 99 | - | 99 | - | | | 96 | 61 | 99 | 20 | 99 | 96 |
| *South_Rolly* | 23 | - | 15 | 8 | 15 | 8 | 15 | - | 23 | - | 23 | - | 23 | - | 23 | - | | | 22 | 23 | 23 | 15 | 23 | 22 |
| *Spirit* | 50 | - | 22 | 28 | 22 | 28 | 18 | - | 50 | - | 50 | - | 50 | - | 50 | - | | | 47 | 37 | 50 | 18 | 50 | 47 |
| *Tern* | 37 | - | 33 | 4 | 33 | 4 | 23 | - | 37 | - | 37 | - | 37 | - | 37 | - | | | 36 | 25 | 37 | 23 | 37 | 36 |
| *Walby* | 60 | - | 24 | 35 | 24 | 35 | 23 | - | 59 | - | 59 | - | 59 | - | 59 | - | | | 0 | 44 | 60 | 23 | 59 | - |
| *Watson* | 60 | - | 24 | 36 | 24 | 36 | 19 | - | 60 | - | 60 | - | 60 | - | 60 | - | | | 59 | 56 | 60 | 19 | 60 | 59 |
| *Wik* | 50 | - | 17 | 33 | 17 | 33 | 17 | - | 50 | - | 50 | - | 50 | - | 50 | - | | | 49 | 38 | 50 | 17 | 50 | 49 |

Table S3. Proportion of variance explained by first five linear discriminant axes for each trait suite and the first three trait suites combined. Also included are the sample size of each LDA (*N*), and the effective dimensionality of trait divergence ($n_D$).

| **Trait Suite** | LD1 | LD2 | LD3 | LD4 | LD5 | Sum LDs 1-5 | N |
|---|---|---|---|---|---|---|---|

| | | | | | | | |
|---|---|---|---|---|---|---|---|
| **Defense** | 0.921 | 0.049 | 0.017 | 0.009 | 0.005 | 1 | 551 |
| **Swimming** | 0.675 | 0.256 | 0.069 | NA | NA | 1 | 792 |
| **Trophic** | 0.445 | 0.255 | 0.132 | 0.073 | 0.035 | 0.940 | 266 |
| **Trophic no GR** | 0.429 | 0.292 | 0.122 | 0.082 | 0.044 | 0.970 | 789 |
| **Shape** | 0.352 | 0.170 | 0.119 | 0.103 | 0.073 | 0.818 | 616 |
| **Total** | 0.690 | 0.104 | 0.078 | 0.040 | 0.032 | 0.945 | 241 |
| **Total Excluding GR** | 0.714 | 0.092 | 0.075 | 0.034 | 0.027 | 0.941 | 549 |

Table S4. Trait loadings on the first 5 discriminant axis in combined Defensive, Swimming, and Trophic trait linear discriminant analysis.

| **Trait** | LD1 | LD2 | LD3 | LD4 | LD5 |
|---|---|---|---|---|---|
| **LP** | -0.226 | 0.038 | -0.016 | 0.245 | 0.327 |
| **$DS1_L$** | 0.458 | 0.366 | 0.352 | 0.476 | 0.406 |
| **$DS2_L$** | 0.610 | 1.430 | 0.491 | 0.540 | -0.432 |
| **$PS_L$** | -4.215 | -0.438 | 0.282 | -0.085 | -0.544 |
| **$LP_M$** | 0.005 | 0.254 | -0.460 | 0.196 | 0.285 |
| **SL** | 1.010 | -1.736 | 0.631 | 0.098 | 0.409 |
| **BD** | -0.310 | 0.908 | -0.730 | -0.381 | -0.251 |
| **CP** | 0.058 | -0.673 | -1.008 | 2.174 | -0.86 |
| **BC** | -0.356 | 0.443 | 0.236 | -0.315 | -0.064 |
| **GW** | -0.012 | 0.543 | 0.178 | -0.886 | 0.448 |
| **PW** | -0.033 | 0.139 | -1.279 | -0.223 | -0.593 |
| **GR** | -0.012 | -0.193 | 0.342 | 0.138 | -0.304 |
| **JL** | 0.248 | -0.472 | -0.120 | -0.211 | 0.636 |
| **SN** | 0.330 | 0.295 | 0.890 | -0.408 | -0.68 |
| **ED** | -0.347 | -0.098 | 0.057 | 1.025 | 1.785 |
| **HL** | -0.338 | -0.540 | 0.578 | -0.758 | -0.953 |
| **Cumulative proportion** | **0.690** | **0.794** | **0.872** | **0.912** | **0.944** |

Table S5. Trait loadings in Defensive trait linear discriminant analysis.

| **Trait** | LD1 | LD2 | LD3 | LD4 | LD5 |
|---|---|---|---|---|---|
| **$PS_L$** | -3.698 | 0.034 | -0.461 | -0.044 | 0.142 |
| **LP** | -0.079 | 0.010 | 1.092 | 0.395 | -0.179 |
| **$LP_M$** | 0.003 | 0.030 | -0.222 | 0.897 | 0.524 |
| **$DS1_L$** | 0.304 | 0.768 | 0.448 | -0.705 | 1.648 |
| **$DS2_L$** | 0.474 | 0.553 | -0.369 | 0.635 | -1.766 |
| **Cumulative proportion** | **0.921** | **0.970** | **0.986** | **0.995** | **1** |

Table S6. Trait loadings in Swimming trait linear discriminant analysis

| **Trait** | LD1 | LD2 | LD3 |
|---|---|---|---|
| **SL** | -1.973 | 1.203 | 1.073 |
| **BD** | 2.285 | -0.118 | 0.697 |
| **CP** | -0.478 | -1.940 | -0.865 |
| **Cumulative proportion** | **0.675** | **0.931** | **1** |

Table S7. Trait loadings in Trophic trait linear discriminant analysis

| **Trait** | LD1 | LD2 | LD3 | LD4 | LD5 | LD6 | LD7 | LD8 |
|---|---|---|---|---|---|---|---|---|
| **BC** | 0.008 | 0.310 | -0.226 | -0.566 | -1.379 | -1.428 | -0.230 | -1.614 |
| **GW** | 0.389 | -0.268 | 0.805 | 0.601 | 0.227 | 1.027 | 1.021 | -1.203 |
| **PW** | 1.275 | 1.479 | -0.628 | 0.934 | -0.848 | -0.328 | -0.800 | 1.859 |
| **GR** | -0.529 | -0.063 | 0.290 | 0.126 | -0.612 | 0.008 | 0.701 | 0.483 |
| **JL** | -0.043 | -0.267 | -1.418 | 0.977 | 0.763 | -0.800 | 0.699 | -0.134 |
| **SN** | -1.108 | 0.310 | 1.905 | 1.765 | 0.307 | 0.239 | -1.565 | 0.312 |
| **ED** | 0.069 | -2.184 | -0.077 | 0.079 | -0.653 | 0.377 | -0.680 | 0.517 |

| | | | | | | | | |
|---|---|---|---|---|---|---|---|---|
| **HL** | -1.137 | 0.821 | -1.115 | -3.162 | 1.425 | 1.330 | 1.191 | 0.015 |
| **Cumulative proportion** | **0.445** | **0.700** | **0.832** | **0.904** | **0.940** | **0.964** | **0.983** | **1** |

Table S8. Correlations between primary axes of divergence in trait suites across populations. Sample sizes represent the overlap in the number of individuals with complete data from both trait suites in the comparison.

| LDA primary axes | Pearson's *r* | p | Spearman's ρ | p | N |
|---|---|---|---|---|---|
| **Defense x Swimming** | -0.47 | <.001 | -0.42 | <.001 | 551 |
| **Defense x Trophic** | -0.21 | 0.001 | -0.10 | 0.109 | 241 |
| **Defense x Shape** | -0.61 | <.001 | -0.59 | <.001 | 432 |
| **Swimming x Trophic** | 0.63 | <.001 | 0.65 | <.001 | 266 |
| **Swimming x Shape** | 0.71 | <.001 | 0.72 | <.001 | 616 |
| **Trophic x Shape** | 0.65 | <.001 | 0.65 | <.001 | 199 |

Table S9. Distance matrix correlations of standardized trait values calculated using mantel tests. All linear measurements were adjusted for size before Z-transformation except for standard length. All mantel tests used 5000 permutations.

| Trait Suites | Pearson's *r* | p | Spearman's ρ | p |
|---|---|---|---|---|
| **Defense x Swimming** | 0.121 | <2e-04 | 0.112 | <2e-04 |
| **Defense x Trophic** | 0.078 | 0.0046 | 0.082 | <2e-04 |
| **Defense x Shape** | 0.033 | 0.1104 | 0.037 | 0.055 |
| **Swimming x Trophic** | 0.789 | <2e-04 | 0.705 | <2e-04 |
| **Swimming x Shape** | 0.115 | <2e-04 | 0.124 | <2e-04 |
| **Trophic x Shape** | 0.092 | <2e-04 | 0.101 | <2e-04 |

Table S10. Statistical results of two-block PLS correlation tests between trait suites within each lake. P-values are FDR-adjusted for multiple comparisons. Tests with adjusted p-values ≤ 0.05 are bold and italicized. Negative Z-scores indicate that trait suites are less integrated than would be expected given random associations of traits.

| Lake | Defense x Swimming | | | Defense x Trophic | | | Defense x Shape | | | Swimming x Trophic | | | Swimming x Shape | | | Trophic x Shape | | |
|---|---|---|---|---|---|---|---|---|---|---|---|---|---|---|---|---|---|---|
| | r-PLS | p | Z | r-PLS | p | Z | r-PLS | p | Z | r-PLS | p | Z | r-PLS | p | Z | r-PLS | p | Z |
| **Corcoran** | **0.719** | **0.003** | **3.00** | **0.775** | **0.021** | **2.40** | 0.584 | 0.689 | 0.08 | **0.949** | **0.001** | **6.07** | **0.619** | **0.003** | **2.99** | 0.682 | 0.056 | 1.72 |
| **Echo** | **0.837** | **0.002** | **4.91** | **0.868** | **0.021** | **2.40** | **0.794** | **0.006** | **3.24** | **0.986** | **0.001** | **3.80** | **0.839** | **0.002** | **4.22** | **0.949** | **0.012** | **2.69** |
| **Engineer** | **0.612** | **0.004** | **3.12** | 0.708 | 0.427 | 0.43 | 0.502 | 0.379 | 0.74 | **0.855** | **0.011** | **2.36** | **0.662** | **0.002** | **4.70** | **0.927** | **0.041** | **1.76** |
| **Finger** | **0.768** | **0.002** | **6.22** | **0.887** | **0.007** | **4.13** | - | - | - | **0.972** | **0.001** | **5.71** | - | - | - | - | - | - |
| **G** | **0.462** | **0.003** | **3.82** | **0.582** | **0.034** | **2.44** | 0.337 | 0.689 | -0.18 | **0.910** | **0.001** | **6.49** | **0.656** | **0.002** | **6.93** | **0.849** | **0.012** | **3.68** |
| **Jean** | 0.385 | 0.129 | 1.22 | 0.570 | 0.429 | 0.25 | 0.552 | 0.226 | 1.32 | **0.912** | **0.001** | **5.27** | **0.770** | **0.002** | **6.29** | **0.767** | **0.034** | **2.07** |
| **Long** | **0.871** | **0.002** | **6.06** | **0.872** | **0.007** | **4.68** | 0.795 | 0.362 | 0.99 | **0.954** | **0.001** | **6.83** | **0.681** | **0.008** | **2.71** | **0.926** | **0.024** | **2.10** |
| **Ruth** | **0.776** | **0.002** | **7.27** | **0.711** | **0.021** | **2.63** | 0.552 | 0.08 | 2.16 | **0.988** | **0.001** | **6.34** | **0.686** | **0.002** | **7.32** | **0.772** | **0.035** | **1.91** |
| **South Rolly** | **0.693** | **0.003** | **3.38** | 0.751 | 0.054 | 1.93 | 0.487 | 0.95 | -1.10 | **0.920** | **0.001** | **4.32** | **0.740** | **0.014** | **2.14** | 0.821 | 0.055 | 1.54 |
| **Spirit** | **0.426** | **0.048** | **2.24** | 0.443 | 0.672 | -0.49 | 0.384 | 0.689 | -0.28 | **0.899** | **0.001** | **5.05** | **0.726** | **0.002** | **5.33** | **0.804** | **0.024** | **2.66** |
| **Tern** | **0.914** | **0.002** | **5.50** | **0.836** | **0.014** | **3.31** | **0.770** | **0.006** | **3.70** | **0.971** | **0.001** | **7.01** | **0.764** | **0.002** | **4.83** | **0.716** | **0.031** | **2.91** |
| **Walby** | **0.613** | **0.004** | **4.26** | 0.533 | 0.427 | 0.38 | - | - | - | **0.924** | **0.001** | **5.72** | - | - | - | - | - | - |
| **Watson** | **0.709** | **0.002** | **6.20** | 0.702 | 0.078 | 1.78 | 0.511 | 0.171 | 1.64 | **0.955** | **0.001** | **5.47** | **0.518** | **0.012** | **2.50** | 0.670 | 0.168 | 0.97 |
| **Wik** | 0.434 | 0.080 | 1.52 | 0.632 | 0.120 | 1.50 | 0.333 | 0.962 | -1.71 | **0.963** | **0.001** | **5.41** | **0.688** | **0.002** | **4.60** | **0.776** | **0.034** | **2.02** |

Table S11. Results of complete modularity test, including partitions between defense, swimming, and trophic traits. CR is the mean of covariance ratios between assigned modules. P-values are FDR-adjusted for multiple comparisons.

| Lake | CR | 95% CI Lower | 95% CI Upper | p | Z |
|---|---|---|---|---|---|
| Corcoran | **1.072** | **0.931** | **1.229** | **0.022** | **-1.108** |
| Echo | **1.083** | **0.932** | **1.358** | **0.041** | **-1.341** |
| Engineer | 1.093 | 0.948 | 1.766 | 0.076 | -1.039 |
| Finger | **1.101** | **0.978** | **1.191** | **0.022** | **-0.939** |
| G | **0.869** | **0.742** | **1.052** | **0.017** | **-1.002** |
| Jean | **0.839** | **0.768** | **1.114** | **0.017** | **-2.092** |
| Long | **1.112** | **1.026** | **1.172** | **0.017** | **-1.461** |
| Ruth | **1.062** | **0.843** | **1.197** | **0.024** | **-1.211** |
| South_Rolly | **1.019** | **0.790** | **1.181** | **0.017** | **-1.881** |
| Spirit | **0.659** | **0.647** | **1.047** | **0.017** | **-2.351** |
| Tern | **1.100** | **1.026** | **1.179** | **0.017** | **-1.522** |
| Walby | **0.867** | **0.702** | **1.129** | **0.017** | **-2.378** |
| Watson | **0.995** | **0.823** | **1.180** | **0.017** | **-2.137** |
| Wik | **0.837** | **0.689** | **1.166** | **0.017** | **-2.552** |

**Table S12.** Results of focused modularity tests for defense, swimming, and trophic trait suites when a single partition is set between the trait suite of interest and all other traits. P-values are FDR-adjusted to account for multiple comparisons, and significant adjusted p-values are indicated in bold typeface.

| Lake | Defense | | | | | Swimming | | | | | Trophic | | | | |
|---|---|---|---|---|---|---|---|---|---|---|---|---|---|---|---|
| | CR | 95% CI lower | 95% CI upper | p | Z | CR | 95% CI lower | 95% CI upper | p | Z | CR | 95% CI lower | 95% CI upper | p | Z |
| Corcoran | **0.959** | **0.767** | **1.246** | **0.002** | **-4.353** | 1.150 | 1.071 | 1.194 | 0.281 | -0.427 | **1.020** | **0.915** | **1.128** | **0.038** | **-3.73** |
| Echo | **0.984** | **0.757** | **1.371** | **0.005** | **-3.514** | 1.171 | 1.130 | 1.229 | 0.281 | -0.473 | 1.139 | 1.069 | 1.173 | 0.806 | 0.806 |
| Engineer | **0.974** | **0.767** | **1.340** | **0.019** | **-2.166** | 1.188 | 1.088 | 1.888 | 0.399 | -0.32 | 1.024 | 0.869 | 1.22 | 0.124 | -1.716 |
| Finger | **1.029** | **0.872** | **1.142** | **0.003** | **-2.901** | 1.127 | 1.082 | 1.161 | 0.243 | -0.543 | 1.095 | 1.054 | 1.133 | 0.270 | -0.668 |
| G | **0.701** | **0.521** | **0.968** | **0.001** | **-7.636** | 1.121 | 1.036 | 1.200 | 0.243 | -0.302 | **1.017** | **0.906** | **1.106** | **0.038** | **-3.030** |
| Jean | **0.700** | **0.591** | **1.085** | **0.001** | **-4.394** | 1.072 | 0.927 | 1.190 | 0.281 | -0.714 | 1.012 | 0.864 | 1.183 | 0.177 | -1.234 |
| Long | **1.046** | **0.906** | **1.135** | **0.001** | **-2.993** | 1.131 | 1.102 | 1.159 | 0.243 | -0.726 | **1.056** | **0.997** | **1.092** | **0.038** | **-2.998** |
| Ruth | **0.927** | **0.606** | **1.120** | **0.001** | **-4.568** | 1.179 | 1.167 | 1.207 | 0.302 | -0.457 | 1.120 | 1.083 | 1.151 | 0.683 | 0.350 |
| South_Rolly | **0.931** | **0.645** | **1.160** | **0.001** | **-4.319** | 1.115 | 0.999 | 1.172 | 0.243 | -0.815 | 1.079 | 0.980 | 1.133 | 0.177 | -1.362 |
| Spirit | **0.420** | **0.398** | **0.978** | **0.001** | **-8.984** | 1.067 | 0.982 | 1.191 | 0.243 | -0.657 | **0.982** | **0.780** | **1.115** | **0.038** | **-3.042** |
| Tern | **1.014** | **0.913** | **1.115** | **0.002** | **-4.197** | 1.156 | 1.127 | 1.179 | 0.281 | -0.557 | 1.071 | 1.010 | 1.114 | 0.067 | -2.690 |
| Walby | **0.730** | **0.533** | **1.097** | **0.001** | **-5.574** | 1.092 | 0.948 | 1.138 | 0.243 | -0.691 | 1.058 | 0.932 | 1.124 | 0.219 | -0.941 |
| Watson | **0.870** | **0.656** | **1.111** | **0.001** | **-5.699** | 1.144 | 1.066 | 1.206 | 0.281 | -0.504 | 1.076 | 1.005 | 1.149 | 0.216 | -0.985 |
| Wik | **0.654** | **0.443** | **1.130** | **0.001** | **-8.787** | 1.146 | 0.992 | 1.180 | 0.243 | -0.621 | 1.112 | 0.956 | 1.153 | 0.177 | -1.226 |

**Table S13.** Results of complete modularity test, including partitions between defense, swimming, trophic and shape traits. CR is the mean of covariance ratios between assigned modules. P-values are FDR-adjusted for multiple comparisons.

| Lake | CR | 95% CI lower | 95% CI upper | p | Z |
|---|---|---|---|---|---|
| Corcoran | **0.842** | **0.820** | **1.063** | **0.006** | **-1.861** |
| Echo | 0.982 | 0.900 | 1.193 | 0.053 | -1.353 |
| Engineer | 1.010 | 0.934 | 1.535 | 0.084 | -1.113 |
| G | **0.791** | **0.727** | **0.963** | **0.002** | **-1.496** |
| Jean | **0.764** | **0.753** | **1.044** | **0.001** | **-2.268** |
| Long | **0.943** | **0.885** | **1.150** | **0.029** | **-1.663** |
| Ruth | **0.870** | **0.758** | **1.024** | **0.029** | **-1.395** |
| South_Rolly | **0.928** | **0.802** | **1.095** | **0.029** | **-1.468** |
| Spirit | **0.717** | **0.709** | **1.016** | **0.001** | **-2.731** |
| Tern | 0.977 | 0.873 | 1.102 | 0.053 | -1.393 |
| Watson | **0.804** | **0.753** | **1.048** | **0.005** | **-1.709** |
| Wik | **0.783** | **0.709** | **1.053** | **0.002** | **-2.254** |

Table S14**.** Results of focused modularity tests for Defense, Swimming, Trophic, and Shape traits when a single partition is set between the trait suite of interest and all other traits. P-values are FDR-adjusted to account for multiple comparisons, and significant adjusted p-values are indicated in bold typeface.

| Lake | Defense | | | | | Swimming | | | | | Trophic | | | | | Shape | | | | |
|---|---|---|---|---|---|---|---|---|---|---|---|---|---|---|---|---|---|---|---|---|
| | CR | 95% CI lower | 95% CI upper | p | Z | CR | 95% CI lower | 95% CI upper | p | Z | CR | 95% CI lower | 95% CI upper | p | Z | CR | 95% CI lower | 95% CI upper | p | Z |
| Corcoran | 0.905 | 0.790 | 1.408 | 0.091 | -1.564 | 0.779 | 0.664 | 1.001 | 0.055 | -1.411 | **0.696** | **0.619** | **0.930** | **0.001** | **-4.798** | **0.587** | **0.559** | **0.917** | **< 0.001** | **-10.25** |
| Echo | 0.982 | 0.840 | 1.352 | 0.154 | -1.069 | 1.025 | 0.901 | 1.158 | 0.218 | -0.779 | 0.945 | 0.848 | 1.052 | 0.051 | -1.848 | **0.838** | **0.746** | **1.023** | **< 0.001** | **-5.719** |
| Engineer | 1.004 | 0.859 | 1.355 | 0.226 | -0.851 | 1.061 | 0.899 | 1.929 | 0.268 | -0.64 | **0.851** | **0.771** | **1.101** | **0.006** | **-2.924** | **0.863** | **0.836** | **1.039** | **0.001** | **-4.054** |
| G | **0.702** | **0.608** | **0.997** | **0.002** | **-3.228** | 0.896 | 0.761 | 1.026 | 0.13 | -0.733 | **0.846** | **0.763** | **0.954** | **0.008** | **-3.065** | **0.797** | **0.720** | **0.931** | **< 0.001** | **-6.082** |
| Jean | 0.879 | 0.771 | 1.272 | 0.067 | -1.764 | **0.709** | **0.643** | **0.965** | **0.034** | **-1.619** | **0.772** | **0.716** | **0.992** | **0.002** | **-3.945** | **0.762** | **0.725** | **0.966** | **< 0.001** | **-6.563** |
| Long | 0.952 | 0.838 | 1.175 | 0.111 | -1.406 | 0.931 | 0.778 | 1.111 | 0.130 | -1.159 | **0.899** | **0.799** | **1.072** | **0.016** | **-2.563** | **0.772** | **0.707** | **1.032** | **< 0.001** | **-7.124** |
| Ruth | 0.983 | 0.803 | 1.144 | 0.259 | -0.708 | 1.084 | 0.936 | 1.135 | 0.349 | -0.497 | 0.827 | 0.686 | 0.954 | 0.064 | -1.944 | **0.580** | **0.519** | **0.790** | **< 0.001** | **-8.420** |
| South_Rolly | 0.908 | 0.765 | 1.194 | 0.083 | -1.642 | 0.936 | 0.797 | 1.061 | 0.131 | -0.971 | **0.863** | **0.776** | **0.995** | **0.010** | **-2.836** | **0.798** | **0.732** | **0.955** | **< 0.001** | **-5.967** |

| | | | | | | | | | | | | | | | | | | | | |
|---|---|---|---|---|---|---|---|---|---|---|---|---|---|---|---|---|---|---|---|---|
| **Spirit** | **0.638** | **0.570** | **1.126** | **0.002** | **-4.009** | 0.795 | 0.675 | 1.071 | 0.055 | -1.478 | **0.803** | **0.726** | **0.976** | **0.002** | **-3.847** | **0.757** | **0.712** | **0.946** | **< 0.001** | **-7.230** |
| **Tern** | 1.057 | 0.968 | 1.135 | 0.251 | -0.586 | 1.076 | 0.953 | 1.123 | 0.268 | -0.608 | **0.863** | **0.712** | **1.010** | **0.017** | **-2.832** | **0.761** | **0.591** | **0.977** | **< 0.001** | **-8.224** |
| **Watson** | **0.801** | **0.733** | **1.229** | **0.033** | **-1.94** | 0.797 | 0.672 | 1.016 | 0.056 | -1.145 | **0.619** | **0.562** | **0.871** | **0.001** | **-4.732** | **0.541** | **0.527** | **0.858** | **< 0.001** | **-9.314** |
| **Wik** | **0.679** | **0.561** | **1.145** | **0.002** | **-3.294** | 0.989 | 0.776 | 1.093 | 0.218 | -0.823 | **0.817** | **0.674** | **0.983** | **0.006** | **-3.343** | **0.741** | **0.643** | **0.939** | **< 0.001** | **-7.398** |

Table S15. Eigenvalues, singular values, lake values of principal components, and PC loadings for an environmental PCA including only physico-chemical environmental variables.

| | PC1 | PC2 | PC3 | PC4 | PC5 | PC6 | PC7 | PC8 | PC9 |
|---|---|---|---|---|---|---|---|---|---|
| Eigenvalues | 4.528 | 1.57 | 1.373 | 0.754 | 0.306 | 0.257 | 0.131 | 0.073 | 0.007 |
| Proportion Explained | 0.503 | 0.174 | 0.153 | 0.084 | 0.034 | 0.029 | 0.015 | 0.008 | 0.001 |
| Corcoran | -0.419 | -0.093 | 0.28 | -0.447 | 0.197 | 0.018 | 0.024 | -0.091 | 0.202 |
| Engineer | -0.052 | 0.296 | 0.419 | 0.59 | -0.017 | -0.304 | -0.264 | -0.359 | 0.08 |
| Finger | -0.365 | 0.511 | -0.161 | 0.121 | 0.295 | 0.12 | 0.519 | 0.126 | -0.197 |
| G | 0.414 | -0.308 | 0.178 | 0.112 | 0.39 | -0.207 | 0.389 | 0.274 | 0.313 |
| Jean | -0.256 | -0.07 | -0.571 | -0.072 | 0.202 | -0.368 | -0.457 | 0.089 | 0.266 |
| Long | 0.115 | -0.233 | 0.044 | -0.188 | -0.203 | -0.011 | 0.219 | -0.465 | 0.203 |
| South_Rolly | 0.324 | 0.171 | 0.103 | -0.187 | -0.175 | -0.377 | -0.14 | 0.464 | -0.379 |
| Spirit | 0.265 | 0.45 | -0.098 | -0.107 | -0.372 | 0.414 | -0.086 | 0.147 | 0.506 |
| Tern | -0.192 | -0.466 | -0.241 | 0.524 | -0.325 | 0.244 | 0.092 | 0.217 | -0.061 |
| Walby | -0.223 | -0.1 | 0.097 | -0.242 | -0.49 | -0.201 | 0.162 | -0.062 | -0.293 |
| Watson | -0.031 | -0.174 | 0.343 | -0.052 | 0.274 | 0.531 | -0.436 | 0.149 | -0.25 |
| Wik | 0.42 | 0.017 | -0.395 | -0.052 | 0.225 | 0.141 | -0.022 | -0.488 | -0.39 |
| Area | 0.001 | 0.673 | -0.262 | 0.436 | -0.072 | 0.42 | 0.151 | -0.116 | 0.265 |
| Max Depth | 0.231 | 0.347 | -0.584 | -0.273 | 0.166 | -0.176 | -0.403 | 0.281 | -0.332 |
| DOC | 0.056 | 0.516 | 0.575 | -0.201 | -0.459 | -0.209 | -0.08 | 0.314 | -0.021 |
| TP | -0.369 | 0.107 | 0.392 | 0.148 | 0.474 | 0.353 | -0.517 | 0.046 | -0.24 |
| TN | -0.34 | -0.125 | -0.166 | 0.66 | -0.374 | -0.375 | -0.178 | 0.133 | -0.276 |
| Chl-a | -0.38 | 0.328 | 0.02 | -0.059 | 0.437 | -0.639 | 0.188 | -0.27 | 0.191 |
| Cond. | -0.441 | 0.058 | -0.1 | -0.198 | 0.026 | 0.22 | 0.611 | 0.35 | -0.456 |
| pH | -0.404 | 0.057 | -0.162 | -0.401 | -0.442 | 0.147 | -0.237 | -0.599 | -0.124 |
| Ca | -0.435 | -0.139 | -0.205 | -0.187 | -0.079 | 0.073 | -0.209 | 0.485 | 0.651 |

Table S16. Eigenvalues, singular values, lake values of principal components, and PC loadings for an environmental PCA including physico-chemical and foraging-associated environmental variables.

| | PC1 | PC2 | PC3 | PC4 | PC5 | PC6 | PC7 | PC8 |
|---|---|---|---|---|---|---|---|---|
| Eigenvalues | 5.545 | 2.506 | 1.805 | 0.948 | 0.558 | 0.371 | 0.213 | 0.054 |
| Proportion Explained | 0.462 | 0.209 | 0.15 | 0.079 | 0.046 | 0.031 | 0.018 | 0.004 |
| Finger | 0.324 | -0.674 | 0.129 | 0.023 | 0.319 | 0.1 | 0.443 | 0.067 |
| Jean | 0.291 | -0.245 | -0.518 | -0.221 | -0.494 | 0.206 | -0.372 | 0.046 |
| Long | -0.045 | 0.296 | -0.074 | 0.311 | -0.193 | -0.096 | 0.285 | 0.755 |
| South_Rolly | -0.362 | 0.054 | -0.247 | 0.583 | 0.169 | 0.457 | -0.036 | -0.34 |
| Spirit | -0.396 | -0.259 | 0.115 | -0.076 | 0.343 | -0.365 | -0.584 | 0.234 |
| Tern | 0.386 | 0.535 | -0.232 | -0.293 | 0.552 | -0.025 | -0.014 | -0.093 |
| Walby | 0.323 | 0.083 | 0.258 | 0.382 | -0.293 | -0.551 | -0.058 | -0.415 |
| Watson | -0.003 | 0.186 | 0.705 | -0.243 | -0.191 | 0.494 | -0.135 | 0.014 |
| Wik | -0.517 | 0.025 | -0.135 | -0.466 | -0.211 | -0.219 | 0.47 | -0.268 |
| Area | -0.075 | -0.493 | -0.033 | -0.265 | 0.685 | -0.26 | 0.052 | 0.218 |
| Max Depth | -0.266 | -0.344 | -0.37 | -0.152 | -0.187 | -0.018 | -0.196 | -0.507 |
| DOC | -0.17 | -0.199 | 0.33 | 0.699 | 0.249 | -0.037 | -0.451 | -0.237 |
| TP | 0.279 | -0.138 | 0.433 | -0.15 | 0.059 | 0.648 | -0.116 | 0.087 |
| TN | 0.384 | 0.076 | -0.225 | -0.169 | 0.238 | -0.014 | -0.122 | -0.51 |
| Chl-a | 0.271 | -0.452 | -0.065 | 0.152 | -0.192 | 0.155 | 0.308 | -0.173 |
| Cond. | 0.382 | -0.249 | 0.09 | 0.009 | 0.147 | -0.091 | 0.191 | 0.01 |
| pH | 0.359 | -0.17 | 0.014 | 0.066 | -0.362 | -0.535 | -0.324 | 0.358 |
| Ca | 0.404 | -0.075 | -0.073 | -0.13 | -0.132 | 0.079 | -0.449 | -0.046 |
| Daph. | -0.267 | -0.018 | 0.387 | -0.56 | -0.073 | -0.073 | -0.376 | -0.062 |
| Gamm. | 0.197 | 0.418 | -0.36 | 0.068 | 0.394 | 0.094 | -0.296 | 0.129 |
| % littoral area | 0.219 | 0.313 | 0.467 | -0.061 | 0.082 | -0.414 | 0.246 | -0.438 |

Table S17. Results of two-block PLS tests of phenotypic trait groups against environmental variables. Adjusted p-values are adjusted for multiple comparisons using the FDR method. Because diet-related environmental variables were available for only eight lakes, statistics are shown from PLS tests including all environmental variables on the left, and those including physico-chemical environmental variables only on the right.

| | All Env. Variables | | | | Physico-chemical Env. Variables | | | |
|---|---|---|---|---|---|---|---|---|
| | $r_{PLS}$ | p | Z | adj. p | $r_{PLS}$ | p | Z | adj. p |
| Defense | 0.852 | 0.096 | 1.259 | 0.144 | 0.821 | 0.044 | 1.632 | 0.066 |
| Swimming | 0.904 | 0.049 | 1.463 | 0.144 | 0.822 | 0.031 | 1.922 | 0.066 |
| Trophic | 0.787 | 0.298 | 0.639 | 0.298 | 0.544 | 0.698 | -0.523 | 0.698 |
| All traits | 0.893 | 0.066 | 1.412 | NA | 0.799 | 0.096 | 1.337 | NA |

Table S18. Singular values, eigenvalues, and vector loadings of environmental variables and stickleback traits in Two-Block PLS correlations between environmental and trait suite data. PLS correlation is calculated using singular value decomposition rather than eigen decomposition of matrices, but because singular values are square roots of eigenvalues, eigenvalues are provided for comparison with LDA and RDA results for which eigenvalues are provided.

| | Dim1 | Dim2 | Dim3 | Dim4 | Dim5 | Dim1 | Dim2 | Dim3 | Dim1 | Dim2 | Dim3 | Dim4 | Dim5 | Dim6 | Dim7 | Dim8 |
|---|---|---|---|---|---|---|---|---|---|---|---|---|---|---|---|---|
| Singular Values | 2.258 | 1.023 | 0.531 | 0.363 | 0.022 | 1.464 | 0.821 | 0.092 | 2.357 | 0.778 | 0.302 | 0.276 | 0.211 | 0.111 | 0.072 | 0.016 |
| $SV^2$ | 5.098 | 1.046 | 0.282 | 0.132 | 0 | 2.143 | 0.674 | 0.008 | 5.554 | 0.605 | 0.091 | 0.076 | 0.044 | 0.012 | 0.005 | 0 |
| Area | -0.321 | 0.602 | 0.394 | 0.276 | -0.237 | -0.056 | 0.149 | -0.038 | -0.179 | 0.322 | 0.14 | 0.629 | -0.285 | 0.401 | 0.376 | 0.258 |
| Max_Depth | -0.434 | 0.452 | -0.084 | -0.039 | 0.509 | 0.22 | 0.31 | -0.092 | 0.296 | 0.115 | -0.443 | 0.443 | -0.299 | -0.552 | -0.006 | -0.228 |
| DOC | -0.105 | -0.052 | 0.499 | -0.578 | -0.477 | 0.08 | 0.176 | -0.19 | 0.06 | 0.406 | 0.007 | 0.365 | 0.776 | -0.041 | -0.299 | -0.005 |
| TP | 0.347 | -0.102 | 0.69 | 0.439 | 0.273 | -0.415 | -0.173 | -0.3 | -0.472 | -0.231 | 0.369 | 0.141 | 0.245 | -0.489 | 0.46 | -0.217 |
| TN | 0.381 | 0.241 | -0.22 | 0.118 | -0.224 | -0.357 | -0.544 | -0.297 | -0.307 | -0.469 | 0.107 | 0.41 | -0.147 | 0.199 | -0.61 | -0.267 |
| ChlA | 0.304 | 0.001 | 0.199 | -0.383 | 0.521 | -0.289 | -0.363 | 0.604 | -0.055 | -0.494 | -0.659 | 0.137 | 0.34 | 0.246 | 0.328 | 0.113 |
| Cond. | 0.366 | 0.287 | -0.106 | -0.084 | -0.049 | -0.479 | 0.411 | 0.494 | -0.504 | 0.301 | -0.28 | -0.236 | 0.024 | 0.125 | -0.096 | 0.011 |
| pH | 0.224 | 0.417 | 0.037 | -0.451 | 0.123 | -0.335 | 0.408 | -0.082 | -0.313 | 0.327 | -0.274 | -0.115 | -0.098 | 0.122 | 0.084 | -0.646 |
| Ca | 0.395 | 0.326 | -0.109 | 0.157 | -0.209 | -0.466 | 0.239 | -0.401 | -0.452 | 0.059 | -0.221 | -0.011 | -0.137 | -0.404 | -0.250 | 0.578 |
| LP | 0.411 | 0.485 | -0.769 | 0.041 | -0.056 | | | | | | | | | | | |
| $DS1_L$ | 0.413 | -0.533 | -0.174 | -0.138 | 0.704 | | | | | | | | | | | |
| $DS2_L$ | 0.497 | -0.496 | -0.003 | -0.149 | -0.697 | | | | | | | | | | | |
| $PS_L$ | 0.519 | 0.483 | 0.552 | -0.424 | 0.115 | | | | | | | | | | | |
| $LP_M$ | 0.379 | 0.043 | 0.273 | 0.882 | 0.050 | | | | | | | | | | | |
| SL | | | | | | 0.672 | -0.413 | 0.615 | | | | | | | | |
| BD | | | | | | -0.539 | -0.842 | 0.022 | | | | | | | | |
| CP | | | | | | 0.508 | -0.346 | -0.789 | | | | | | | | |
| BC | | | | | | | | | 0.378 | -0.216 | 0.421 | -0.506 | 0.499 | -0.239 | 0.19 | 0.188 |
| GW | | | | | | | | | -0.037 | -0.632 | -0.662 | -0.096 | 0.043 | -0.036 | 0.384 | 0.024 |
| PW | | | | | | | | | 0.111 | -0.622 | 0.364 | 0.124 | -0.234 | 0.532 | -0.29 | 0.175 |
| GR | | | | | | | | | 0.444 | 0.352 | -0.305 | -0.248 | 0.14 | 0.705 | 0.086 | -0.033 |
| JL | | | | | | | | | 0.46 | 0.084 | 0.083 | 0.652 | -0.071 | -0.094 | 0.468 | 0.34 |
| SN | | | | | | | | | 0.464 | 0.037 | -0.278 | -0.25 | -0.477 | -0.356 | -0.437 | 0.314 |
| ED | | | | | | | | | 0.235 | -0.122 | -0.234 | 0.412 | 0.606 | -0.104 | -0.546 | -0.17 |
| HL | | | | | | | | | 0.406 | -0.139 | 0.141 | 0.007 | -0.274 | -0.123 | 0.128 | -0.83 |

Table S19. Singular values, eigenvalues, and vector loadings of environmental variables and the traits in defensive, swimming, and trophic trait suites in Two-Block PLS correlations between environmental and trait data.

| | Dim1 | Dim2 | Dim3 | Dim4 | Dim5 | Dim6 | Dim7 | Dim8 | Dim9 |
|---|---|---|---|---|---|---|---|---|---|
| Singular Values | 3.434 | 1.55 | 0.813 | 0.735 | 0.51 | 0.285 | 0.162 | 0.1 | 0.04 |
| $SV^2$ | 11.793 | 2.401 | 0.661 | 0.54 | 0.26 | 0.081 | 0.026 | 0.01 | 0.002 |
| Area | -0.026 | 0.654 | 0.509 | -0.299 | -0.047 | 0.147 | -0.354 | 0.031 | -0.272 |
| Max_Depth | -0.34 | 0.385 | 0.286 | 0.392 | -0.161 | -0.415 | 0.395 | 0.118 | 0.365 |
| DOC | -0.089 | 0.149 | -0.299 | -0.238 | -0.649 | 0.446 | 0.35 | 0.285 | 0.001 |
| TP | 0.431 | -0.06 | 0.153 | -0.647 | -0.145 | -0.436 | 0.239 | -0.158 | 0.274 |
| TN | 0.369 | -0.202 | 0.569 | 0.199 | 0.113 | 0.55 | 0.259 | 0.06 | 0.272 |
| ChlA | 0.206 | -0.332 | 0.26 | 0.304 | -0.666 | -0.245 | -0.375 | 0.093 | -0.185 |
| Cond. | 0.465 | 0.305 | -0.315 | 0.139 | 0.1 | -0.043 | -0.355 | 0.503 | 0.426 |
| pH | 0.299 | 0.346 | -0.228 | 0.291 | -0.212 | 0.139 | -0.019 | -0.76 | 0.108 |
| Ca | 0.454 | 0.186 | -0.069 | 0.211 | 0.135 | -0.184 | 0.452 | 0.19 | -0.646 |
| LP | 0.277 | 0.01 | 0.069 | 0.752 | 0.451 | 0.28 | 0.12 | 0.091 | -0.005 |
| $DS1_L$ | 0.202 | -0.497 | -0.394 | -0.015 | 0.058 | 0.02 | -0.026 | 0.052 | 0.19 |
| $DS2_L$ | 0.261 | -0.501 | -0.4 | -0.072 | -0.052 | 0.083 | 0.04 | 0.05 | -0.087 |
| $PS_L$ | 0.368 | 0.046 | 0.025 | 0.193 | -0.636 | 0.162 | 0.164 | -0.49 | -0.045 |
| $LP_M$ | 0.253 | -0.14 | 0.11 | -0.207 | 0.225 | -0.382 | 0.265 | 0.074 | -0.539 |
| SL | -0.278 | -0.241 | 0.135 | -0.141 | 0.128 | 0.336 | -0.265 | 0.066 | 0.117 |
| BD | 0.237 | -0.307 | 0.543 | -0.07 | -0.166 | 0.265 | -0.262 | 0.133 | -0.116 |
| CP | -0.208 | -0.184 | 0.143 | -0.159 | 0.15 | 0.348 | 0.377 | -0.243 | -0.382 |
| BC | -0.237 | -0.286 | -0.088 | 0.001 | -0.012 | -0.131 | 0.089 | -0.221 | 0.175 |
| GW | 0.059 | -0.232 | 0.284 | 0.25 | -0.157 | -0.564 | -0.276 | 0.089 | -0.041 |
| PW | -0.051 | -0.283 | 0.374 | 0.02 | 0.137 | -0.017 | -0.06 | -0.239 | 0.147 |
| GR | -0.303 | -0.032 | -0.279 | 0.249 | -0.197 | 0.153 | -0.503 | 0.02 | -0.518 |
| JL | -0.315 | -0.079 | 0.098 | 0.104 | -0.226 | 0.038 | 0.302 | 0.126 | -0.177 |
| SN | -0.305 | -0.139 | -0.081 | 0.351 | 0.027 | -0.234 | 0.112 | -0.099 | -0.182 |

| | | | | | | | | | |
|---|---|---|---|---|---|---|---|---|---|
| ED | -0.147 | -0.129 | 0.073 | 0.122 | -0.368 | 0.071 | 0.385 | 0.638 | 0.174 |
| HL | -0.265 | -0.188 | 0.073 | 0.155 | 0.016 | -0.134 | 0.116 | -0.336 | 0.272 |

Table S20. Singular values, eigenvalues, and vector loadings of environmental variables and stickleback traits in Two-Block PLS correlations between environmental and trait suite data. PLS correlation is calculated using singular value decomposition rather than eigen decomposition of matrices, but because singular values are square roots of eigenvalues, eigenvalues are provided for comparison with LDA and RDA results for which eigenvalues are provided.

| | Dim1 | Dim2 | Dim3 | Dim4 | Dim5 | Dim1 | Dim2 | Dim3 | Dim1 | Dim2 | Dim3 | Dim4 | Dim5 | Dim6 | Dim7 | Dim8 |
|---|---|---|---|---|---|---|---|---|---|---|---|---|---|---|---|---|
| Singular Values | 3.119 | 1.004 | 0.717 | 0.172 | 0.043 | 1.676 | 1.209 | 0.192 | 3.468 | 1.247 | 0.508 | 0.41 | 0.32 | 0.096 | 0.061 | 0.003 |
| SV$^2$ | 9.725 | 1.008 | 0.514 | 0.03 | 0.002 | 2.809 | 1.461 | 0.037 | 12.024 | 1.554 | 0.259 | 0.168 | 0.103 | 0.009 | 0.004 | 0 |
| Area | -0.306 | 0.491 | -0.232 | -0.499 | -0.11 | -0.012 | -0.198 | 0.009 | -0.131 | 0.077 | 0.035 | -0.302 | 0.732 | 0.39 | 0.305 | 0.123 |
| Max_Depth | -0.366 | 0.131 | -0.232 | 0.003 | 0.052 | 0.287 | 0.227 | 0.122 | 0.424 | -0.176 | -0.257 | -0.097 | 0.332 | -0.283 | -0.12 | -0.455 |
| DOC | -0.13 | -0.34 | 0.094 | -0.721 | 0.04 | 0.135 | -0.474 | 0.008 | -0.04 | 0.587 | 0.534 | -0.181 | 0.208 | -0.274 | -0.396 | -0.217 |
| TP | 0.198 | 0.187 | 0.56 | -0.037 | -0.613 | -0.405 | -0.238 | 0.146 | -0.474 | -0.074 | 0.102 | -0.301 | -0.135 | -0.551 | 0.322 | 0.172 |
| TN | 0.365 | 0.457 | -0.191 | -0.022 | 0.099 | -0.417 | 0.26 | 0.202 | -0.251 | -0.472 | 0.042 | -0.224 | 0.068 | 0.076 | -0.15 | -0.495 |
| ChlA | 0.209 | -0.162 | 0.104 | -0.188 | -0.154 | -0.279 | 0.098 | -0.459 | -0.029 | -0.235 | 0.21 | 0.393 | 0.234 | -0.321 | 0.248 | -0.139 |
| Litt_Area | 0.215 | 0.057 | 0.309 | 0.065 | 0.606 | -0.318 | -0.439 | -0.154 | -0.54 | 0.239 | -0.151 | 0.27 | -0.169 | 0.295 | -0.049 | -0.501 |
| Cond. | 0.26 | 0.116 | 0.096 | -0.28 | 0.042 | -0.401 | -0.071 | -0.201 | -0.316 | -0.135 | 0.16 | 0.217 | 0.263 | 0.06 | 0.092 | 0.037 |
| pH | 0.135 | 0.009 | 0.093 | -0.259 | 0.402 | -0.204 | -0.02 | 0.121 | -0.117 | -0.055 | -0.058 | 0.385 | 0.24 | 0.061 | -0.494 | 0.368 |
| Ca | 0.287 | 0.283 | 0.14 | -0.168 | 0.08 | -0.398 | 0.129 | 0.389 | -0.24 | -0.342 | -0.035 | 0.003 | 0.13 | -0.223 | -0.458 | 0.185 |
| Daph. | -0.469 | 0.456 | 0.449 | 0.126 | 0.149 | 0.109 | -0.437 | 0.596 | -0.226 | 0.238 | -0.677 | -0.294 | 0.113 | -0.178 | -0.142 | 0.078 |
| Gamm. | 0.323 | 0.226 | -0.434 | 0.031 | -0.127 | -0.098 | 0.382 | 0.357 | 0 | -0.288 | 0.282 | -0.465 | -0.22 | 0.322 | -0.253 | 0.093 |
| LP | 0.503 | 0.21 | -0.834 | 0.02 | 0.084 | | | | | | | | | | | |
| DS1$_{L}$ | 0.527 | -0.295 | 0.197 | 0.502 | -0.586 | | | | | | | | | | | |
| DS2$_{L}$ | 0.565 | -0.178 | 0.368 | -0.006 | 0.717 | | | | | | | | | | | |
| PS$_{L}$ | 0.343 | -0.034 | 0.142 | -0.855 | -0.359 | | | | | | | | | | | |
| LP$_{M}$ | 0.178 | 0.914 | 0.332 | 0.125 | -0.082 | | | | | | | | | | | |
| SL | | | | | | 0.59 | 0.498 | -0.635 | | | | | | | | |
| BD | | | | | | -0.652 | 0.758 | -0.011 | | | | | | | | |
| CP | | | | | | 0.476 | 0.421 | 0.772 | | | | | | | | |
| BC | | | | | | | | | 0.339 | 0.036 | -0.182 | 0.067 | -0.608 | -0.05 | 0.247 | -0.643 |
| GW | | | | | | | | | 0.163 | -0.701 | 0.081 | 0.231 | 0.423 | -0.183 | 0.422 | -0.175 |
| PW | | | | | | | | | 0.144 | -0.556 | -0.22 | 0.065 | -0.238 | 0.608 | -0.416 | 0.131 |
| GR | | | | | | | | | 0.367 | 0.385 | 0.279 | 0.607 | 0.185 | 0.463 | 0.144 | 0.044 |
| JL | | | | | | | | | 0.501 | 0.165 | -0.296 | -0.6 | 0.443 | 0.241 | 0.079 | -0.112 |
| SN | | | | | | | | | 0.413 | 0.036 | -0.116 | 0.274 | 0.178 | -0.5 | -0.668 | -0.105 |
| ED | | | | | | | | | 0.369 | -0.146 | 0.796 | -0.358 | -0.232 | -0.072 | -0.091 | 0.114 |
| HL | | | | | | | | | 0.38 | -0.006 | -0.315 | 0.072 | -0.28 | -0.257 | 0.326 | 0.707 |

Table S21. Singular values, eigenvalues, and vector loadings of environmental variables and the traits in defensive, swimming, and trophic trait suites in Two-Block PLS correlations between environmental and trait data. Seven dimensions with singular values <0.0001 are excluded from this table.

| | Dim1 | Dim2 | Dim3 | Dim4 | Dim5 | Dim6 | Dim7 | Dim8 |
|---|---|---|---|---|---|---|---|---|
| Singular Values | 4.37 | 2.788 | 1.409 | 0.645 | 0.53 | 0.347 | 0.176 | 0.07 |
| SV$^2$ | 19.092 | 7.773 | 1.986 | 0.417 | 0.281 | 0.12 | 0.031 | 0.005 |
| Area | 0.045 | -0.36 | 0.409 | 0.379 | 0.563 | -0.154 | -0.264 | -0.245 |
| Max_Depth | 0.431 | 0.08 | 0.372 | -0.194 | 0.182 | 0.074 | 0.167 | 0.513 |
| DOC | 0.064 | -0.276 | -0.52 | 0.337 | 0.408 | 0.351 | 0.399 | 0.271 |
| TP | -0.421 | -0.224 | 0.054 | -0.153 | -0.083 | 0.678 | -0.247 | -0.067 |
| TN | -0.367 | 0.208 | 0.39 | 0.204 | 0.046 | -0.01 | 0.056 | 0.489 |
| ChlA | -0.146 | 0.194 | -0.054 | -0.437 | 0.419 | 0.071 | -0.199 | 0.143 |
| Litt_Area | -0.442 | -0.325 | -0.219 | 0.05 | -0.168 | -0.498 | -0.05 | 0.427 |
| Cond. | -0.35 | -0.013 | 0.031 | -0.085 | 0.335 | -0.126 | -0.082 | -0.085 |
| pH | -0.154 | 0.024 | -0.017 | -0.213 | 0.185 | -0.277 | 0.548 | -0.358 |
| Ca | -0.324 | 0.11 | 0.246 | -0.151 | 0.062 | 0.145 | 0.501 | -0.099 |
| Daph | 0.072 | -0.644 | 0.366 | -0.124 | -0.299 | 0.065 | 0.248 | 0.005 |
| Gama | -0.15 | 0.352 | 0.152 | 0.597 | -0.177 | 0.128 | 0.134 | -0.11 |
| LP | -0.249 | 0.435 | 0.061 | 0.633 | 0.124 | -0.463 | 0.014 | 0.296 |
| DS1$_{L}$ | -0.293 | 0.337 | -0.365 | -0.163 | -0.257 | 0.115 | -0.158 | -0.041 |
| DS2$_{L}$ | -0.34 | 0.313 | -0.336 | -0.189 | -0.182 | 0.1 | 0.202 | 0.056 |
| PS$_{L}$ | -0.211 | 0.186 | -0.162 | 0.012 | 0.149 | 0.158 | 0.269 | -0.168 |
| LP$_{M}$ | -0.193 | -0.023 | 0.486 | 0.119 | -0.202 | 0.104 | 0.287 | -0.027 |
| SL | 0.238 | 0.172 | -0.001 | 0.173 | -0.29 | -0.158 | -0.532 | -0.299 |
| BD | -0.211 | 0.343 | 0.348 | -0.202 | 0.204 | 0.047 | -0.039 | -0.366 |
| CP | 0.191 | 0.125 | 0.101 | 0.242 | -0.424 | 0.075 | 0.365 | -0.536 |
| BC | 0.244 | 0.18 | -0.072 | -0.111 | -0.341 | -0.028 | -0.078 | 0.244 |
| GW | 0.051 | 0.306 | 0.362 | -0.343 | 0.313 | 0 | -0.214 | 0.008 |
| PW | 0.053 | 0.248 | 0.305 | -0.226 | -0.208 | -0.16 | -0.034 | 0.08 |
| GR | 0.285 | 0.113 | -0.341 | -0.051 | 0.354 | -0.404 | 0.099 | -0.43 |
| JL | 0.391 | 0.167 | 0.049 | 0.165 | 0.059 | 0.177 | 0.249 | 0.171 |
| SN | 0.299 | 0.206 | -0.048 | -0.225 | 0.128 | -0.124 | 0.445 | 0.184 |
| ED | 0.231 | 0.305 | -0.074 | 0.304 | 0.25 | 0.673 | -0.197 | 0.044 |
| HL | 0.276 | 0.194 | 0.002 | -0.198 | -0.228 | -0.068 | 0.016 | 0.228 |
